# Comparative analysis of *Glycine BBX* gene family reveals lineage-specific evolution and expansion

**DOI:** 10.1101/2022.11.16.516718

**Authors:** Reena Deshmukh, Sourav Datta

## Abstract

*BBX* genes are associated with photomorphogenesis, hormone response and seed gemination. Although, *BBX* gene family is reported in *G. max*, little is known about its classification and expansion. Similarly, no information is available for the *BBX* gene family in its closest relative *Glycine soja* (Siebold & Zucc.). With extensive genome diversity, *G. soja* can be used as an effective genetic reservoir for introgressing important agronomic traits in *G. max*. In the present study, we carried out a comprehensive comparative genome-wide analysis of *BBX* gene family in *G. max* and *G. soja*, to identify their evolutionary relationship and origin in plant lineage. Our results show an ancient *BBX* gene family expansion through segmental duplication, further suggesting, that the *GmBBX* members are the out-paralogs. These genes show lineage-specific evolution and expansion in the ancestral *Glycine* genome supported by the incidences of microsynteny between *G. max* and *G. soja*. The two genomes also showed interesting evidence of conserved linkages which might be due to their common ancestor descendance, with minimum horizontal expansion in *G. max* after its split from *G. soja*. Our study suggests that the *BBX* gene family diverged before the split of *G. max* and *G. soja*. As the two genomes share several regions of synteny, the paralogous members in *G. max* could have been a result of segmental duplications.

## 1. Introduction

Soybean is amongst the very few legumes to have been sequenced. With the release of its genome sequence, the era of legume genomics has been revolutionized (Schmutz et al., 2010). Plant genomes have been constantly evolving and are much more complex than the animal genomes (Jiao and Paterson, 2014). Instances of polyploidy has considerably contributed towards the plant genome complexity (Adams et al., 2005). Polyploidy can be derived either from genome multiplication through interspecies crosses, called as allopolyploidy or genome multiplication through autopoplyploidy involving Whole Genome Duplication (WGD) (del Pozo et al., 2015). These events result in the emergence of multiple copies of genes leading to the expansion of gene families after the last speciation event or the last common ancestor split. Allopolyploidy and autopolyploid significantly cause abrupt gene family expansion that may or may not be similar in all descendants of a monophyletic clade. Although polyploidy have played a significant role in the domestication of several crop species, it has however resulted in multiple copies of genes that has increased genetic redundancy and sometimes to gene losses (Adams et al., 2005). A well-known example of such genome multiplication through allopolyploidy is in the *Brassicaceae* family, which includes the model eudicot plant, *Arabidopsis thaliana*. Contrastingly, genus *Glycine*, is the result of genome autoploploidy, a paleopolyploid with two rounds of WGD. The first WGD is reported to have occurred around 59 million years ago (Mya), while the second WGD at around 13 Mya, followed by chromosomal rearrangements (Shoemaker et al., 2006; Schmutz et al., 2010).

The genus *Glycine* is supposed to have diverged into two sub-genera viz., *Glycine and Soja*, around 5 to 10 Mya (Doyle and Egan, 2010; Zhuang et al., 2022). The sub-genus *soja* includes the present domesticated species *Glycine soja max* and its wild progenitor *Glycine soja soja* (Sedivy et al., 2017; Zhuang et al., 2022). *G. max* evolved as a result of extensive multiple domestications events in *G. soja* that spread across various geographical regions in different time periods ranging between 5,000 and 9,000 years ago, demonstrating the two species cross-compatibility (Sedivy et al., 2017). Although, their crosses often result in huge amounts of linkage drag associated with candidate gene linkage disequilibrium (Carter et al., 2004; Kim et al., 2012; Kofsky et al., 2018), rigorous selection procedures in *G. soja* subjective to local environment, led to slow and continuous variations in their phenotypic and genotypic behavior (Carter et al., 2004; Kim et al., 2010; 2012). Both the genomes share similar diploid number (*2n=40*), a comparable genome of approximately 1.1Gb, several analogous protein coding genes and syntenic regions. However, the two species vary considerably in their phenotypic traits that are suggested to be controlled by both quantitative and qualitative genetic mechanisms. Some of these differences includes seed size and color, oil and protein percentage in seeds, pod and seed number and general growth habit (Joshi et al., 2013; Xie et al., 2019). The genome of *G. soja* is much more diverse in comparison to *G. max* which has lost it genetic diversity owing to domestication, and stringent advanced breeding (Hyten et al., 2006; Kofsky et al., 2018). The diversity in *G. soja* offers large and wider range of genetic variations, that can be used to introgress important agronomic traits. Thus, analyzing the gene families in the two species, will provide an understanding of varied background for the improvement of cultivated soybean.

*BBX* genes have been identified in several plant species (Crocco and Botto, 2013). These The BBX TFs belong to the zinc-finger binding domain protein, with a conserved one or two B-Box domain(s) with or without CCT (CONSTANS, CO-like and TOC1) domain (Torok and Etkin, 2001; Khanna et al., 2009; Gangappa and Botto, 2014). Several independent *BBX* genes have been known to regulate seedling photomorphogenesis, flowering and circadian clock in plants (Vaishak et al., 2019). *AtBBX32* regulates flowering in Arabidopsis (Tripathi et al., 2017), and *AtBBX24* induces root growth under high salt conditions (Nagaoka et al., 2003). *AtBBX21* regulates photomorphogenesis, anthocyanin accumulation, while *AtBBX18* and *AtBBX23* regulates thermomorphogenesis (Xu et al., 2016; Ding et al., 2018; Xu et al., 2018). Moreover, the *BBX* genes are also associated with the abiotic response in a hormonal regulation. In Arabidopsis, *BBX* genes viz., *BBX1, BBX5* and *BBX21* regulate abscisic acid (ABA) in drought stress (Xu et al., 2014; Min et al., 2015; Riboni et al., 2016). Similarly, *BBX21* interaction with few other genes, modulates the gibberellin metabolism, while its integration with *BBX22* plays significant role in shade avoidance syndrome (Crocco et al., 2010; Xu et al., 2017). Previous studies suggests that clade IV of *AtBBX* genes is actively involved in plant development network (Khanna et al., 2009).

The *BBX* genes have been identified in Arabidopsis and several non-leguminous plants like grapes, tomato, rice, pear, etc. (Chu et al., 2016; Bai et al., 2019; Shalmani et al., 2019; Wei et al., 2020). However, not many leguminous crop species are known with the identified *BBX* gene family members. Hence, we performed a comparative cross-species analysis of *BBX* genes in *G. max* and *G. soja*, studying their evolution and expansion.

## 2. Materials and Methods

### 2.1. Identification of GmBBX and GsBBX gene family members

To detect potential homologous *GmBBX* (*G. max* BBX) and *GsBBX* (*G. soja* BBX) genes, we carried out the genome-wide identification of members in the newest assembly of *G. max* (Glycine_max_v4.0 reference, Annotation Release 104) and *G. soja* (ASM419377v2 reference Annotation Release 100), at the NCBI and Phytozome v13 database (https://phytozome-next.jgi.doe.gov/). The *AtBBX* (Arabidopsis BBX) were used to identify all possible homologs in the two species. The *AtBBX* sequences were downloaded from TAIR (The Arabidopsis Information Resource; http://www.arabidopsis.org/; Rhee et al., 2003; Swarbreck et al., 2008; Lamesch et al., 2012). We used the all-against-all BLASTp search for every AtBBX protein in both the genomes (Altschul et al., 1990) to screen the non-redundant protein database of *G. max* and *G. soja*. Owing to the closeness between the two genomes, we also performed a reciprocal best hit (RBH) to identify orthologs in *G. soja* using *G. max* as query sequences with E-value and sequence coverage as filtering parameters (Moreno-Hagelsieb and Latimer, 2008). We initially filtered the non-redundant sequences for maximum genome span with highly similar sequences at an E-value <10^−5^ and a sequence identity of more than 50% and query coverage greater than 60%. We also did a basic bit score to maximal bit score distribution through simple representative comparison to further refine our RBH ortholog search (Pushker et al., 2004). This was done by sorting out the sequences in ascending order of bit score and/or E-value, with the first sequence being identified as the best scoring match. Sequences with more than one similar score were considered as another ortholog or simply an isoform (depending upon their precise genome location and protein coding sequence; Moreno-Hagelsieb and Latimer, 2008). For this analysis, we selected the GmBBX members with only BBX1 and BBX2 domain. The predicted sequences were again confirmed for non-redundancy. Only the full-length peptides were included in the future annotations.

The probable *BBX* gene family included all the non-redundant sequences harboring BBX domain(s) and/or CCT domain. The identified protein sequence was further confirmed for BBX domain by scanning each homolog in Pfam (Mistry et al., 2021), InterProScan (Jones et al., 2014) and NCBI Conserved Domain database (CDD) for BBX and CCT domains. The corresponding gene sequence of the redundant protein with the overlapping or incomplete sequences were filtered by their chromosome number and relative chromosomal location on the genome browser. Any alternative splice sequence was filtered out to reduce overlapping information of a protein/gene sequence. To identify paralogous genes, we again performed the BLASTp search using each protein sequence against its own genome. However, in-here, to have maximum coverage and to fetch every individual sequence with BBX domain, we restrained the percent identity criterion to >60% as against >50% while using AtBBX as query.

### 2.2. Chromosome localization, nomenclature and gene structure prediction

The genomic/protein sequence and the gene location of *GmBBX* and *GsBBX* gene family members were obtained from the genome sequence database of *G. max* and *G. soja*, respectively from NCBI. These locations were physically mapped on the respective genomes using the MapInspect v1.0 (https://mapinspect.software.informer.com/). For the ease of interpretation, the chromosomes of *G. max* have been abbreviated as Gm01 to Gm20, while that of *G. soja* are referred to as Gs01 to Gs20. The *GmBBX* and *GsBBX* members were named based upon their corresponding location on the chromosome, as adopted for other crop species like cucumber, cotton, rice, etc. (Huang et al., 2012; Gao et al., 2020; Obel et al., 2022; Zhu et al., 2022). The consecutive members on the same chromosome were numbered using the chromosome number followed by the alphabet as GmBBX1(x)-20(x), and GsBBX1(x)-20(x), where x, is the English alphabet in the linear order. This nomenclature methodology was reasonably more convenient to compare the *BBX* family in the two species, especially while performing synteny and conserved linkage analyses. The gene structure of individual *GmBBX* and *GsBBX* genes were predicted using Fgenesh server (http://www.softberry.com), while the gene structure of all the genes with complete sequence were visualized using Gene Structure Display Server (GSDS; http://gsds.gao-lab.org/) (Hu et al., 2015).

### 2.3. Syntenic blocks and gene duplication events in G. max

To analyze the events of gene duplication events in *BBX* gene family in the *G. max* genome, we studied the pattern of segmental and tandem gene duplication. The gene(s) was considered tandemly duplicated if one or more *GmBBX* genes were located on the same chromosome with less than 10 intervening genes. To identify segmentally duplicated *BBX* genes, we used RBH to best identify sequences, that shared >90% sequence identity and along the stretch of >1kb. To further investigate the evolutionary pattern of *BBX* genes in *G. max* we compared the *GmBBX* and *GsBBX* sequences to identify their homologous behavior. As evident from the studies conducted by Cannon and Shoemaker, 2012, we also attempted to identify the incidences of internal syntenic blocks in *G. max* in and around the *BBX* genes. To identify these regions or synteny, if any, shared between the local genic regions of the chromosomes carrying the *BBX* genes, we used the mVISTA (Tools for Comparative Genomics, Frazer et al., 2004), with window size of 100bp. The *BBX* genes/clusters were distributed in a co-linear fashion in the two genomes and hence we suspected incidences of conserved linkages. To identify conserved linkages and pattern of genome evolution, we carried out microsynteny analysis between the chromosomes of the two genomes around the regions carrying *BBX* genes using GEvo visualizer in a CoGe platform (https://genomevolution.org/coge/; Lyons and Freeling, 2008). To perform the GEvo analysis, we manually submitted individual chromosomal sequences with *G. max* as a reference sequence using *Lagan* alignment algorithm.

### 2.4. Structural characterization of GmBBX and GsBBX family members

The conserved motifs of the two gene families were analyzed using MEME suite v5.4.1 (Bailey and Elkan, 1994; Bailey et al., 2015) and WebLogo (Crooks et al., 2004). While other parameters were set to default, the number of motifs were standardized to 7, keeping the motif width ranging between 40 and 50, while we restricted the sites per motif between 20 and 53. Subcellular localization of respective proteins in both the families were predicted using the CELLO v.2.5 server (Yu et al., 2006). Several other chemical and physical parameters of GmBBX and GsBBX proteins were identified through the ProtParam Expasy tool, which includes parameters like aliphatic index, theoretical pI, grand average of hydropathicity (GRAVY), etc.

### 2.5. Protein sequence analysis and phylogenetic studies

The multiple sequence alignment of the *GmBBX* and *GsBBX* gene family members was performed using CLC sequence viewer v8.0 (www.clcbio.com). The ortholog behavior between the two closely related species was predicted through the phylogenetic studies. To study the *BBX* gene family expansion pattern in the two species, gene loss and the genetic divergence between the paralogs and the orthologs, we used MEGA 11 (Kumar et al., 2018) to construct an evolutionary phylogenetic tree using the Maximum likelihood (ML) with Tamura-Nei model of nucleotide substitution and Bayesian statistics. The clades were statistical supported with 1000 bootstrap replications in an equal input model. The rates among the sites were kept uniform, while the alignment file was created using ClustalW. To study the relative divergence, we performed the molecular clock test using ML tree topologies in MEGA 11.

## 3. Results

### 3.1. BBX gene family identification and distribution in G. max and G. soja

The searches were done using the conserved Hidden Markov Models for the BBX domain(s). Our few BLAST searches within the *G. max* and *G. soja* genomes using GmBBX and GsBBX members respectively, carrying BBX and CCT domains returned several other sequences with only CCT domain, without BBX domain. Hence, for maximum genome coverage with reduced redundancy, we used BBX members with only BBX domain(s). Using both the RBH and bit score matrices, we obtained similar number of homologs, along with few partial sequences. These partial sequences were later overlapped to form a complete protein sequence.

We identified almost equal number of *BBX* genes in the two species, sharing similar features within the homologous pairs. A total of 54 *GmBBX* and 55 *GsBBX* orthologs were identified in *G. max* and *G. soja* (Supplementary Table S1). Two *BBX* members in *G. soja* viz., *GsBBX12b* and *GsBBX15c* had an incomplete sequence and they could not be seen overlapping with any other GsBBX protein. They scored high both in the RBH and bit score matrices, but lacked BBX domains. Hence, due to inconsistency in sequence information and lack of domain, we did not include these members in our future studies. Their exclusion did not seem to interfere with the interpretation of our results, due to relative nomenclature of *GsBBX* members and tight conjunction between orthologous pairs. The *BBX* members were renamed in *G. max* and named in *G. soja* based upon their chromosomal locations as described in materials and methods. The distribution of *GmBBX* and *GsBBX* members mapped using MapInspect 1.0 have been illustrated in the Fig. 1A, 1B, 2A, 2B. The nucleotide sequence of the corresponding *BBX* genes were downloaded using the tBLASTn. The average gene size of *GmBBX* and *GsBBX* were comparable with values equivalent to 4011bp and 3648bp, respectively (Supplementary Table S1).

**Fig. 1.**
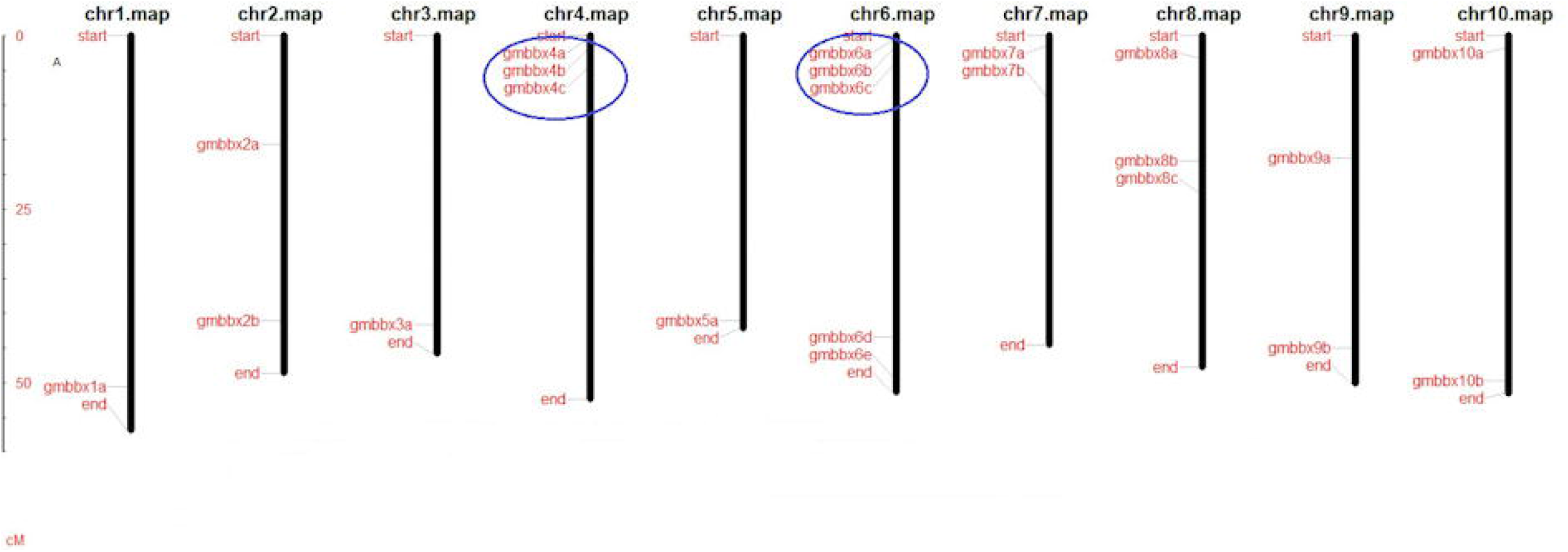
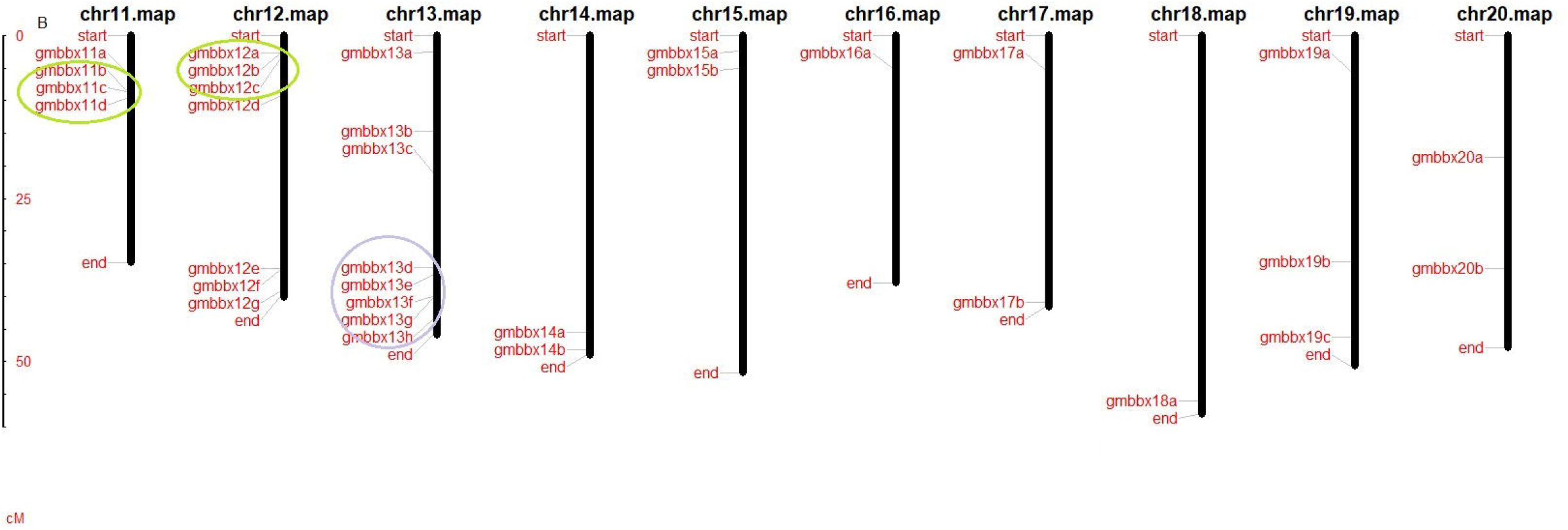
The physical localization of *GmBBX* genes illustrated using MapInspect 1.0. (A) The chromosomal localization of *GmBBX* genes located on Gm01-Gm10. (B) The chromosomal localization of *GmBBX* genes located on Gm11-Gm20. The scale on the left is in ten million bases. The chromosome number is indicated on top of each bar. Note: The consecutive *BBX* gene clusters on *G. max* genome are marked with blue and green. The pink-coloured block represent similar cluster on *G. max* and *G. soja* genomes.

The *BBX* genes in the respective genomes were found to be randomly distributed on all 20 chromosomes. In *G. max* most of the *BBX* genes (8) were localized on Gm13 whilst maximum number of *BBX* genes (9) in *G. soja* were concentrated on Gs11. Interestingly, we observed that all the *GmBBX* and *GsBBX* orthologs shared similar chromosome localization on their respective genomes. The *GsBBX* members that have been excluded from further analysis due to incomplete sequence information, nevertheless, positioned at an approximately similar location on the respective chromosome as their consecutive ortholog member in *G. max* (Fig. 1A, 1B). *BBX* members with similar gene structure were found to group together in a cluster in both the families. While most of *BBX* genes were structured with two or three exons, three *BBX* genes, viz., *GmBBX7a, 8b* and *13d* were intron-less, along with their corresponding orthologs *GsBBX7a, 8b* and *11e*.

Most of the gene pairs of the consecutive syntenic chromosome regions were found to share more than 90% of sequence conservation along >1kb region. However, we could not detect any tandem duplications in the *GmBBX* or the *GsBBX* gene family. To analyze the *BBX* gene family expansion in *G. max*, we first studied if, any, incidence of possible local segmental duplication events in its genome.

### 3.2. Segmental duplications and conserved linkages

The chromosomal distribution of *GmBBX* genes revealed the occurrence of equally spaced *BBX* gene clusters on few chromosomes. These gene clusters showed linear order conservation, with equidistant localization between *BBX* genes of the clusters on chromosomal pairs. Hence, we looked out for the duplicated regions carrying *BBX* genes in mVISTA and compared *BBX* gene regions on consecutive chromosome pairs, Gm04 vs Gm06 and Gm11 vs Gm12 (Supplementary Fig. S1A, 1B). To identify these syntenic regions within the *G. max* genome, we first compared the genic differences between all possible clustered gene(s) within and across the other chromosomes (Supplementary Table S1). The gene pair(s) (within the genome) with comparable distance paired paralogous to each other, with >90% similarity and similar predicted genetic and physiochemical characteristics. These gene pairs when subjected to mVISTA revealed long stretches of regions varying in size between 2-12Mb sequences on different chromosomes. For example, Gm04 and Gm06 shared similarity over 3.58Mb region bearing genes *GmBBX4a,b,c* and *GmBBX6a,b,c*. Similarly, the Gm11 and Gm12 carrying genes *GmBBX11b,c*,d and *GmBBX12a,b*,c respectively, shared approximately a region of 4Mb. Similar incidences were also seen in *G. soja*. These regions apparently appeared to be segmentally duplicated and translocated across the genome.

The genomes of *G. max* and *G. soja* share relative closeness along chromosomal stretches (Kim et al., 2010). In the present study the two genomes displayed almost identical physical maps of the *BBX* gene distribution. Moreover, the representative distribution of *GmBBX* and *GsBBX* seemed to have a conserved gene order of the orthologs in their respective genomes, supposedly demonstrating conserved linkages. These conserved linkages visualized through GEvo identified several microsyntenic regions across the two genomes shared between the collinear chromosomes harboring *BBX* genes. The visualization of orthologous Gm04 vs Gs04 chromosome with BBX region is represented in Supplementary Fig. S2 (the other chromosomes data not shown), except for the region on Gm13 (discussed later).

### 3.3. Domain-based classification and in silico structural characterization of GmBBX and GsBBX proteins

The BBX members in both the species were classified into four distinct groups based upon the number and types of domain present in each (Supplementary Table S2). Members with only CCT domain were filtered out. These four distinct groups classification however, did not coincide with the four-clade classification in a phylogenetic tree. An overall of 39 GmBBXs and 36 GsBBX proteins had a conserved two BBX domains with 20 GmBBX and 18 GsBBX members also having CCT domain. Only seven GmBBXs and GsBBXs each carried one BBX and a CCT domain, ten GmBBXs and GsBBXs each had a single BBX domain (Supplementary Fig. S3).

The predicted molecular weight of GmBBX and GsBBX varied substantially, but they showed remarkable similarity in the individual orthologous nodes with *GmBBX*:*GsBBX* gene pair. The average molecular weight in two BBX families was similar where they shared a close value of 33.3 kDa in GmBBX, while 33.6 kDa in GsBBXs. Likewise, the two families shared similar isoelectric points (pI) with a minimum of 4.2 in GmBBX13d and GsBBX11e, and a maximum of 9.72 shared by GmBBX7a and GsBBX7a. Most of the BBX proteins were predicted to be unstable except for three members viz., BBX8c, BBX12c and BBX18a in the two families which were predicted as stable proteins. Similarly, while most of the GmBBX and GsBBX members were localized in the nucleus, ten GmBBX and GsBBX pairs were localized extracellularly (Supplementary Table S3).

### 3.4. Sequence and motif analysis

We performed an individual and comparative alignment of the two families using CLC sequence viewer. The output was optimized to 80 residues per line. Each orthologous pair shared highly conserved amino acid sequence. Although BBX1 and BBX2 showed similar conserved sequence, the individual domain was almost identical among the orthologous pair (Supplementary Fig. S6). The consensus BBX1 sequence in the two families was identified as C-X_2_-C-X_8_-C-X_2_-D-X-A-X-LC-X_2_-CD-X_3_-H-X_8_-H while BBX2 had a conserved sequence C-X_2_-C-X_8_-C-X_6_-LC-X_2_-CD-X_3_-H-X_8_/X_6_-H. The clade-specific motif identified through MEME and WebLogo3 generated seven motifs. Motif 1 and 3 corresponded to the BBX1 and BBX2 domain, respectively (Fig. 2). Motif 1 was highly conserved in all the BBX members, which was seen in all the clades. The BBX domain members were distributed according to their clade classification (Supplementary Fig. S4). Except clade II, all the three clades shared similar conserved regions in all the members. However, as the clade II was sub-divided into groups of members with one and two BBX domains, the motif was much more staggered (Supplementary Fig. S4). The phylogeny of *Glycine* BBX protein classification clade I included all the members with BBX1:BBX2 domains. Similarly, clade III and IV comprised of members with only BBX1 and BBX1:BBX2:CCT domains respectively. However, clade II consists of members containing BBX1:CCT and BBX1:BBX2:CCT domains (Supplementary Fig. S5A). The protein sequence analysis revealed that the BBX members of clade II shared greater sequence similarity to the clade I members, rather than the members of clade IV, suggesting a greater BBX domain sequence conservation between the clade I and II. Further, we used relative time tree analysis of *Glycine* BBX proteins using AtBBX26 as an outgroup, wherein *Glycine* BBXs were divided into two major clusters, which further divided into four clades. Cluster II seems to be older than cluster I, where clade I possibly evolved earlier than clade II (Supplementary Fig. S5B). Motif sequence conservation was seen to be prominent in the orthologous GmBBX:GsBBX pair, with identical amino acid in the ‘X’ of the consensus sequence, in comparison to the GmBBX or GsBBX paralogs (Supplementary Fig. S6).

**Fig. 2.**
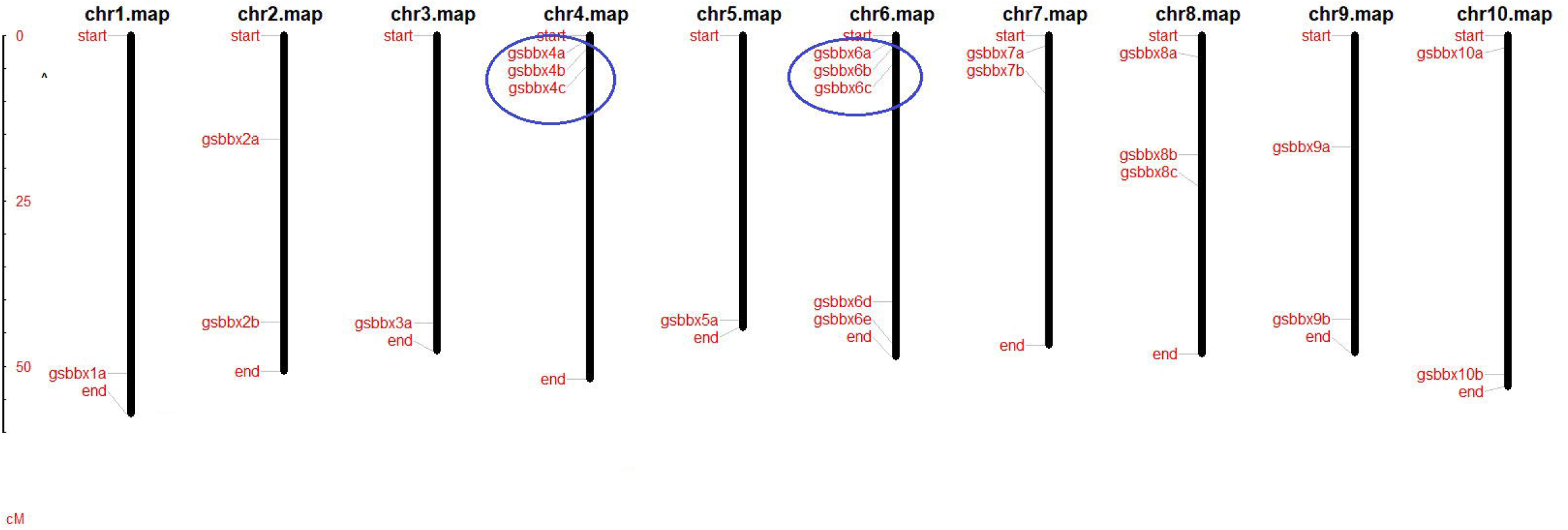
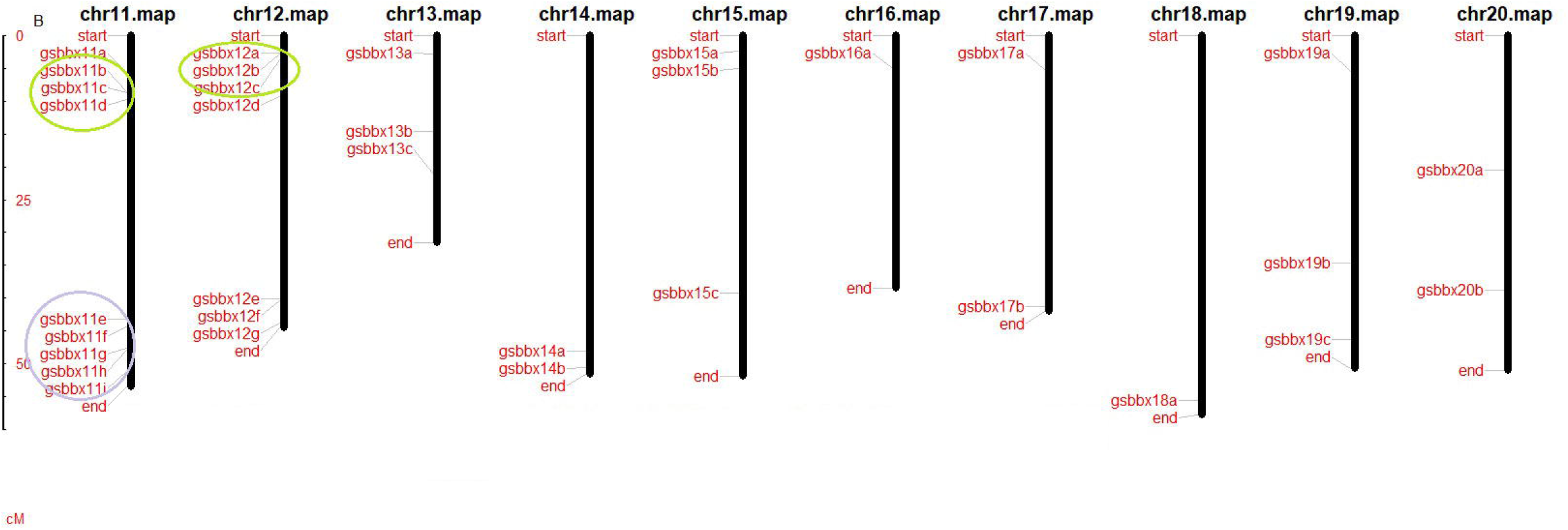
The physical localization of *GsBBX* genes illustrated using MapInspect 1.0. (A) The chromosomal localization of *GsBBX* genes located on Gs01-Gs10. (B) The chromosomal localization of *GsBBX* genes located on Gs11-Gs20. The scale on the left is in ten million bases. The chromosome number is indicated on top of each bar. Note: The consecutive *BBX* gene clusters on *G. soja* genome are marked with yellow and orange. The pink-coloured block represent similar cluster on *G. max* and *G. soja* genomes.

### 3.5. Phylogenetic classification and ancestral evolution of Glycine BBX gene family

Motif analyses of *BBX* genes in *Glycine* genus gave interesting insights into their diversification. However, due to absence of concrete sequence data of *BBX* genes in other species of the *Glycine* genus, the exact origin of the GmBBX may be difficult to predict. Nevertheless, as the sub-genus *soja* has only two species, the prediction of *G. max* can be ascertained through this study. Concisely, both the methods resulted in strong evidential support for the evolution of the *BBX* gene family in *G. max* and *G. soja* through descendance from a common ancestor without any divergence at speciation level, yet (Fig. 3). Similar evolutionary topologies were obtained using both the ML and Bayesian model of their DNA sequences. To gain our clearer understanding into the *BBX* gene family in *G. max*, we analyzed the two families closely by determining their individual paralogous divergence. Although, incidences on few chromosomes like Gm04 and Gm06, etc. suggests expansion of *BBX* genes *G. max* through segmental duplication, followed by chromosomal rearrangements, WGD played a major role in its expansion (Pagel et al., 2004: Schmutz et al., 2010). Hence, to further study the role of WGD in the evolution of *GmBBX*, we classified the *GmBBX* and *GsBBX* together to ascertain their clustering pattern. Interestingly, none of two species-specific paralogs grouped together, and each leaf node was occupied by *GmBBX*:*GsBBX* ortholog pair in 1:1 clustering. For example, *GmBBX4c* always paired with *GsBBX4c* irrespective of their protein or DNA sequences. Simultaneously, although the *GmBBX4c* and *GmBBX6c* paralogs shared >90% sequence identity, the *BBX4c* orthologs were always closer than their corresponding paralog in respective genomes. This type of clustering suggest origination of *BBX4c* orthologs from common ancestor.

**Fig. 3.**
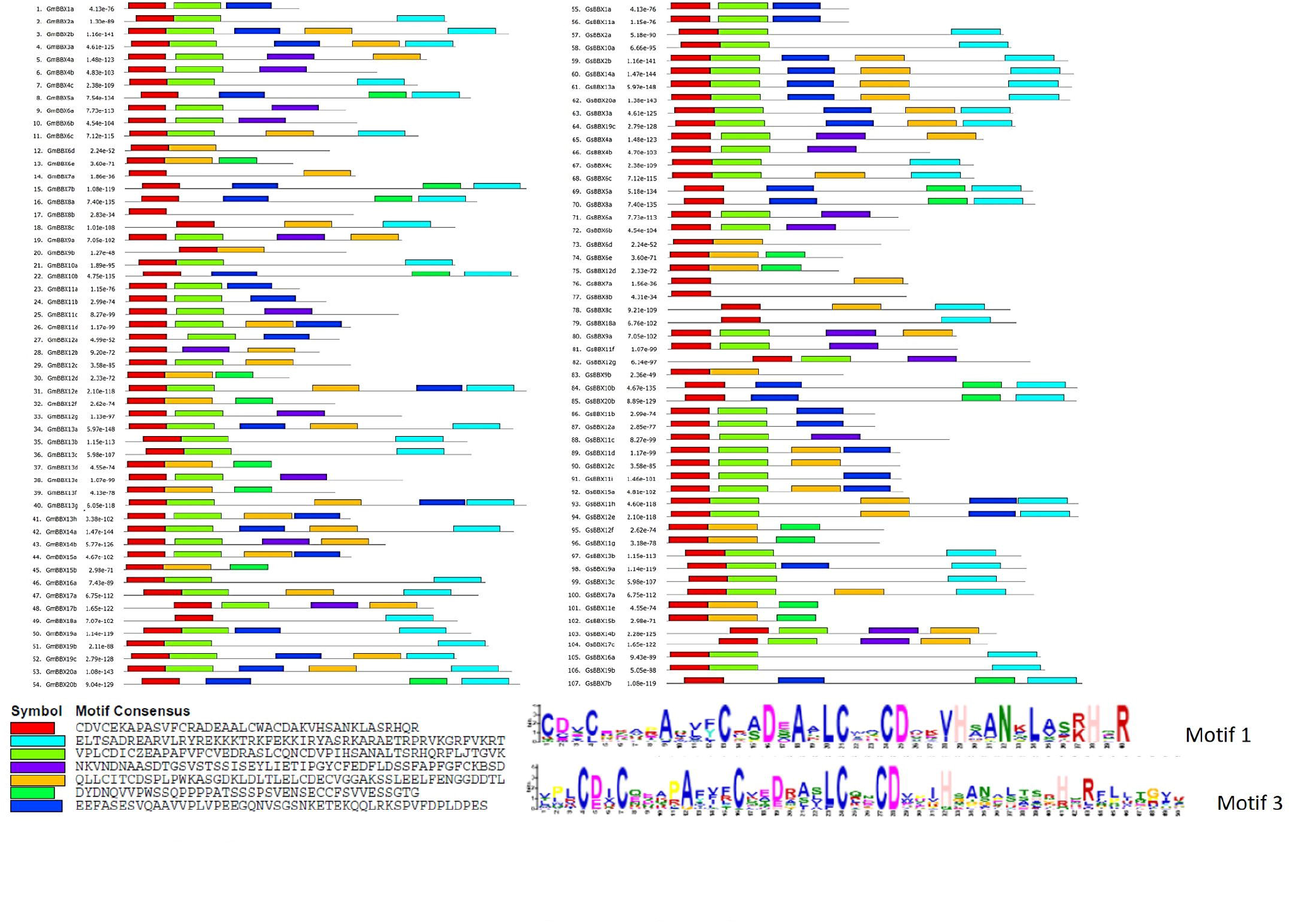
Conserved motif analysis of GmBBX and GsBBX proteins using MEME. The parameters were adjusted to retrieve seven conserved motifs at a motif width between 40 and 50. The sequence logo of each motif is shown at the bottom. Motif 1 and motif 2 correspond to the BBX1 and BBX2 domains of the proteins, respectively.

To further gain our understanding into the evolution and expansion of *BBX* gene family in *Glycine*, we used *AtBBX* to analyze and compare our results. We classified the three *BBX* families using similar statistical parameters in a phylogenetic tree. With respect to the genus *Glycine*, similar *GmBBX*:*GsBBX* clustering towards the leaf was observed, without any family-specific classification. The *BBX* genes clustered together in all the three species. According to the ML clustering, the three *BBX* gene families were divided into four clades, excluding the isolated *AtBBX* members (Fig. 4). These clades were further divided into sub-clades diverging *AtBBX* and *Glycine BBXs*. Homologs from *Glycine* formed monophyletic sub-clades in all the clades (Fig. 4). Using molecular clock test, we identified that these four different clades evolved at different evolutionary rates (Supplementary Fig. S7), although clade II and III have likely to have been evolved at a closer rate (null hypothesis of equal evolutionary rate throughout the tree was rejected at a 5% significance level, *P* = 2.174E-191) (Supplementary Fig. S7). *BBX* members *AtBBX19, 23, 26, 27* and *32*, appeared to be early divergents. According to the ML cladistics, *AtBBX27* may have emerged before clade I, while *AtBBX32* emerged before clade II, and *AtBBX26* separated before clade III and IV comprising of all three *BBX* gene family. While *AtBBX19* and *AtBBX23* seem to have diverged in-parallel, they might have evolved even before *AtBBX27* and *AtBBX32*, i.e., before the split of clade I and II, and could possibly be representative of the immediate Arabidopsis ancestral sequence. Apparently, these members evolved in a lineage-specific manner following speciation in Arabidopsis. Thus, phylogenetic tree revealed that *AtBBX19, 23, 26, 27* and *32* diverged independently from their corresponding neighboring clades and we can assume that their corresponding *Glycine BBX* orthologs were lost during first WGD in the Papilionoideae clade. The nodes of divergence can be understood with respect to their taxonomic divergence, for the above given clades. For instance, *AtBBX27* emerged separately from clade I members and it could be possible that, either it did not diverge further or lost its paralogs during Arabidopsis evolution. This study suggests that different evolutionary rates have played an essential role in sequence diversification, especially after the expansion into four phylogenetic clades. Therefore, conclusively, the phylogenetic studies for the *BBX* gene family suggests that clade diversification occurred before the split of Rosids and Asterids and the additional members in *Glycine* genomes has been horizontally evolved from both segmental duplication and WGD.

**Fig. 4.**
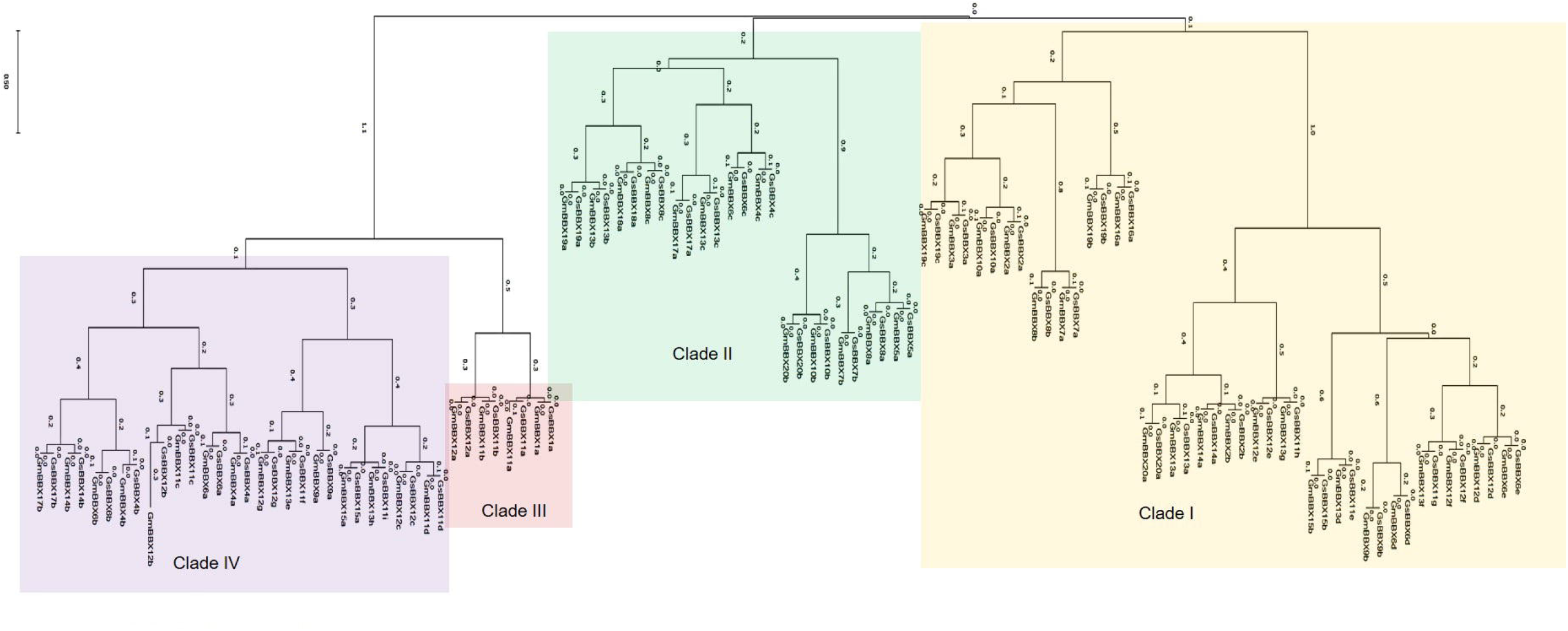
Phylogenetic analysis of Glycine BBXs. The phylogenetic tree of *GmBBX* and *GsBBX* genes was classified into four major clades with *GmBBX:GsBBX* orthologous clustering to the terminal node.

## 4. Discussion

### 4.1. Diversification and origin of BBX gene family in Glycine

*BBX* genes are ubiquitously present in the plant kingdom. Different *BBX* genes have been attributed to several functions like, acting as a regulator of UV-induced photomorphogenesis, circadian cycle, hormone related photomorphic responses, abiotic stress and jasmonic acid-related biotic stress response (Shalmani et al., 2019; Valverde 2011; Gangappa and Botto 2014; Lyu et al., 2020; Zhang et al., 2020). Soybean is an important oilseed crop worldwide with high protein content. Its genome sequencing has made it one of the most popular legume species for molecular studies, owing to its wide geographical cultivation. Soybean evolved after two rounds of WGD as an autoplolyploid, which makes it an interesting candidate to study gene family evolution and diversification. With more than 1.1Gb genome size, *G. max* is expected to carry huge number of *BBX* genes. Although, *BBX* genes have been previously reported in soybean (Preuss et al., 2012), there has been no attempts to classify the family in soybean and there is no evidence for their basis of divergence. Therefore, we performed a comparative genome-wide identification of *BBX* genes in the *Glycine* sub-genus *soja* species to classify the *BBX* gene family and also to ascertain their ancestral origin. We identified a close sequence homology between the *AtBBXs* members with that of *Glycine BBXs*. We also tracked an early diversification in *BBX* gene family, older than the split of Brassicaceae and Papilionoideae, with an ancestral duplication in the genus *Glycine* leading to the expansion of *BBX* genes in soybean.

*Glycine* genome has been subjected to second round of WGD post-Papilionoideae split around 13 Mya (Schmutz et al., 2010). Several gene families have been reported and/or are expected to have been doubled in the *Glycine* genome in comparison to its other homologous genera of the tribe *Phaseoleae*. Arabidopsis genome harbors 32 *BBX* genes with its diploid number 10. Although Brassicaceae and Fabaceae diverged around 125-136 Mya (Hyung et al., 2015), the Arabidopsis evolved from its ancestor recently around 10 to 15 Mya which is approximately around the same time as the second WGD in the genus *Glycine* (Mitchell-Olds, 2001; Hyung et al., 2014). This might suggest that the *AtBBX* and second duplicated *GmBBX* genes would have evolved around the same zoic era. This statement can be extended to that the genes of the *Phaseoleae* family might be relatively older than those of the *Camelineae* (Arabidopsis) family (Frazer et al., 2003; Zhuang et al., 2022). Nevertheless, despite wide lineage or era old separation, several orthologous gene families have maintained their functions and structures in distantly related families. Concurrently, keeping in view of the evolution rate in Arabidopsis and *Glycine*, we expected similar rate of *BBX* gene family expansion with almost four times the *GmBBX* genes as that of Arabidopsis.

Due to recent WGD in the *Glycine* genome, as expected, we identified almost equal number of *BBX* genes i.e., 54 *GmBBX* and 55 *GsBBX* in the genomes of *G. max* and *G. soja*, respectively. Although the two species are suspected to have a shorter time of divergence, around 0.13 to 0.62 Mya (Zhuang et al., 2022), the *G. soja* and *G. max* have shown wide genic variations, which has been also documented in the perennial and annual species of *Brachypodium* (Li et al., 2018). We identified the orthologs using the RBH for better stringency and removal of non-redundant sequences. This type of identical phylogeny shows a clear representation of paralog divergence in the last common ancestor. The *BBX* genes in the two genomes shared similar chromosomal gene organization pattern in the two compatible genomes. This seems to be quite proportionate with respect to the genome organization the two species share.

A family of 62 *GmBBX* genes have been initially identified and has been named from *GmBBX1* to *GmBBX62* (Preuss et al., 2012). However, the present study was encouraged by two major hypotheses. Firstly, as there has been several updates post the Preuss et al., 2012 studies on the genome assembly of *G. max*, there was an immense opportunity for either a deletion due to the general rearrangements of the genome sequence of the unassembled scaffolds and contigs, that might have led to deletion/overlapping of a member(s) or an addition of a family member(s). Secondly, *BBX* gene family members have not been identified in *G. soja*, which could provide a better understanding of the divergence/expansion of *GmBBXs*, and explain its paralogy. This may lead to identification of a better structured evolutionary pattern of the *BBX* gene family in the genus *Glycine*. Hence, in the present study, we specifically emphasized on the identification of *BBXs* in the *G. soja* genome assembly along with the *GmBBXs*.

### 4.2. Structural conservation in GmBBX:GsBBX orthologs

The exon-intron organization was more or less identical in the corresponding *GmBBX:GsBBX* orthologs. Nevertheless, the *GmBBX:GsBBX* orthologs shared similarity, the *GmBBX* and *GsBBX* paralogs showed considerable variation amongst the *GmBBX* and *GsBBX* family, respectively. However, before we propose their functional similarity, further biochemical and sequencing studies needs to be done to establish their orthologous structural and functional nature. Paralogous genes are the result of duplication events, either segmental, tandem or WGD within a species (Koonin 2005). These are horizontally evolved and are highly susceptible to sub-functionalization, neo-functionalization and/or gene loss. The duplicated copy is free to mutate and hence is under lesser selective pressure (Torgerson and Singh 2004). This allows duplicated gene to take up new functions and also undergo substantial sequence variation. Contrastingly, the orthologs are under stricter selection pressure to retain similar function in different species. Orthologs originate through speciation event and descend vertically with greater sequence and function conservation (Altenhoff et al., 2012; Stamboulian et al., 2020).

The BBX2 domain duplicated from BBX1, and hence BBX2 can be expected to have been under lower selection pressure which showed higher variation within *Glycine* BBX paralogs in comparison to BBX1. Despite sharing similar domain architecture, *BBX* genes control different plant functions. Although the differential binding in different clades is not well established, few studies have identified the presence of VP motif in CCT domain of clade IV members of AtBBX family (Gangappa et al., 2014). The VP motif impart functional variations through variable transcriptional binding of HY5 (Jiang et al., 2012; Yadukrishnan et al., 2018). The absence or presence of VP motif has been associated to the occurrence of contrasting traits among the BBX members of different clade (Job et al., 2018). However, whether any other conserved motif in BBX domain is also associated with clade-specific trait in Arabidopsis or other plants, is largely unknown. Moreover, our study grouped few genes with one BBX domain and few genes with two BBX domains in clade II, which was not in agreement with the AtBBX classification (Supplementary Fig. S3). Hence, it presumably appears that the members of these clusters might have evolved from the same ancestral sequence and divergence probably could have resulted due to an internal deletion of the BBX2 domain of the ancestral sequence in clade II (Griffiths et al., 2003). The motifs identified in *Glycine* BBXs were also seen to be conserved in AtBBXs, indicating their conservation in the ancient clade (monophyletic clade in *Glycine* and Arabidopsis BBXs).

### 4.3. Origin of BBX gene family expansion in Glycine

We identified an ancient *BBX* gene family expansion, which could be seen while clustering *AtBBX* genes with *Glycine BBX* genes presumably, a vertical inheritance pattern from their last common ancestor. Every clade comprised of diverged *BBX* genes, which suggests that the diversification in *BBX* gene family has an early origin. We suspect that the *BBX* genes in Arabidopsis and *Glycine* sharing similar clusters, can be perceived as genes being the descendants of the same ancestral sequences. The several members in *Glycine* genus seemed to have evolved horizontally (Zhuang et al., 2022), following two rounds of WGD. Several gene family in plants show lineage and family-specific separation indicating their vertical inheritance (orthologs) while their expansion shows horizontal duplication (paralogs). However, as represented in Fig. 4, we reckon, that *AtBBX19* and *AtBBX23* are the earliest divergents amongst the three *BBX* gene families, without or loss of orthologous *Glycine BBXs*.

There were at least 11 instances detected, where *AtBBX* and *GmBBXs* were seen to have been descended from their common ancestor (for reference we only used *GmBBX* as inclusion of *GsBBX* cluttered the figure, which share similar pairing). For most of the vertical divergence clusters for each *AtBBX* gene, there were at least four copies of *GmBBXs*. This can be presumably suggested due to two rounds of WGD in *G. max*, which is possibly the lineage-specific gene gain in *G. max* (Fig. 5). This can be understood as during the evolution of Arabidopsis, the *Glycine* genome was already doubled due to first WGD, leading to duplicate *BBX* genes. However, during the course of Arabidopsis evolution, *Glycine* genome additionally underwent second WGD, which quadrupled the *BBX* genes. After the two rounds of WGD, *Glycine* genome underwent further genic rearrangement in its chromosomes including several gene losses (Cannon and Shoemaker, 2012). This was interestingly observed in the *GmBBX* family as well. For instance, in the branch 8, 9, 10 and 11 of *AtBBX* and *GmBBX* divergence, two corresponding *AtBBX* orthologs were missing in the *G. max* genome. Similarly, in the sub-branch of branch 11, corresponding *GmBBX* ortholog seemed to have been lost after first WGD. These gene loss or gain events in *G. max* can be well understood by the fact that the second WGD is specific to the *Glycine* genome and has not only resulted in genome doubling but also suffered gene shuffling (Cannon and Shoemaker, 2012).

**Fig. 5.**
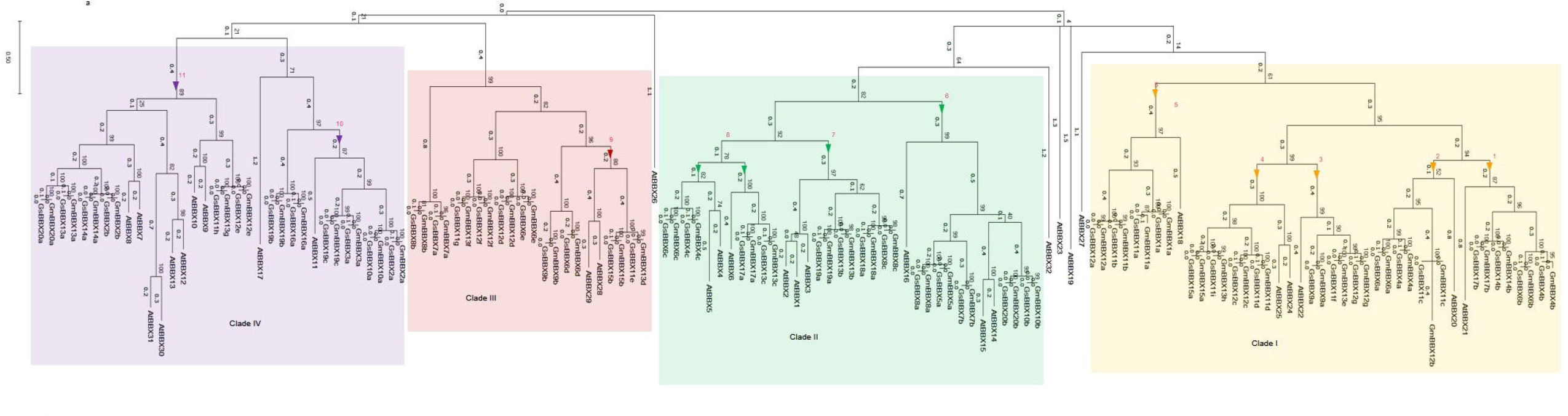
The phylogenetic analysis of *AtBBX, GmBBX* and *GsBBX* genes. The BBX family encodes 54 *GmBBX* and 55 *GsBBX* genes. Branch lengths are proportional to mean substitutions per site. The *BBX* gene clusters in each clade comprised of all three species BBX family, indicating a probable family expansion before the divergence of Arabidopsis and Glycine. The coloured arrows identifies the separation point of Arabidopsis and Glycine *BBXs* illustrated in detail in Fig. 6.

**Fig. 6.**
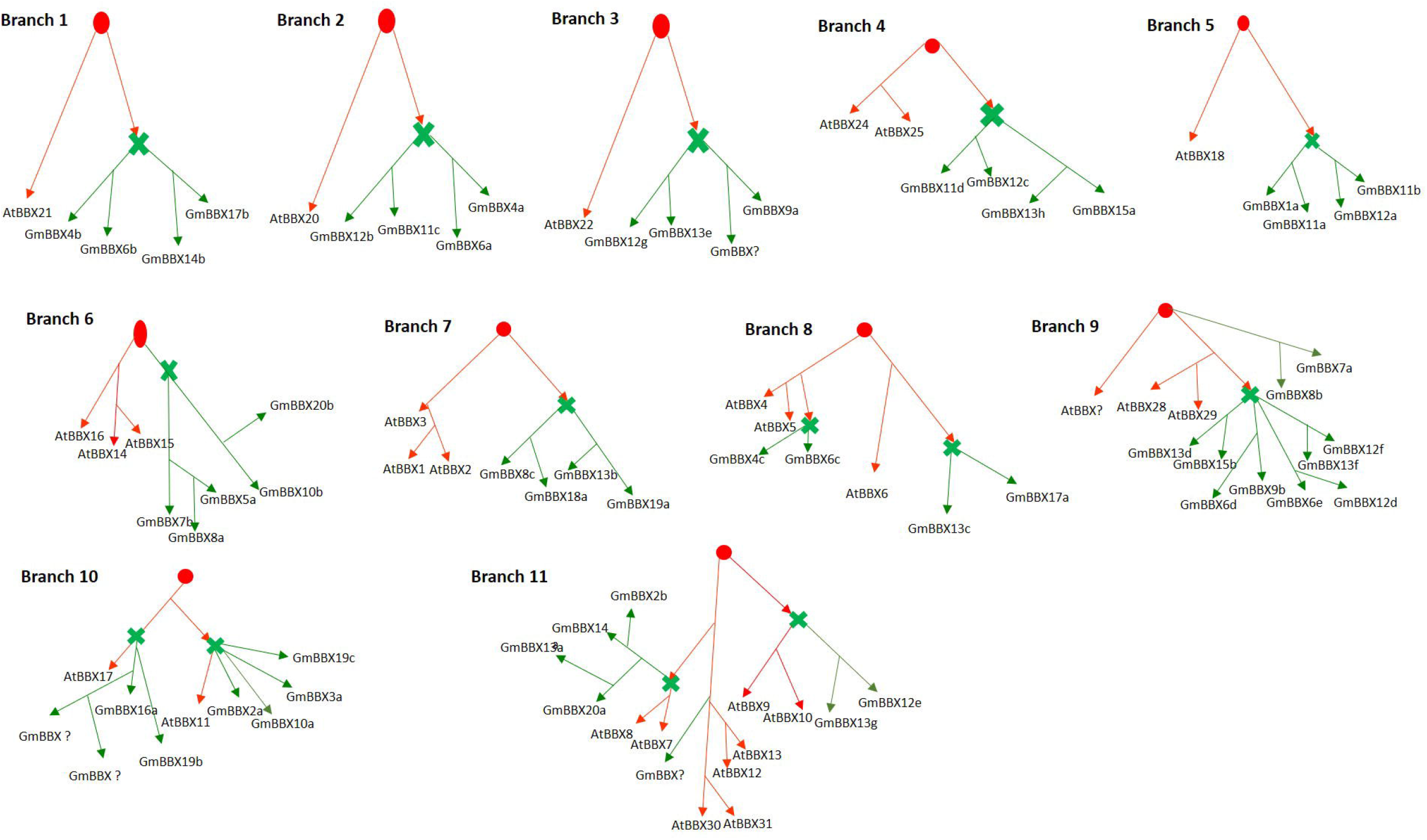
Probable gene duplication events in GmBBX in relation to AtBBX. The branches are derived from Fig. 5. The red colour illustrates divergence between *AtBBX* and *GmBBXs*. The red line identifies speciation in Arabidopsis, while the green line indicates speciation in *G. max*. The green star shows split of *BBX* in Arabidopsis and *G. max*.

### 4.4. Conserved linkages and Glycine paralogy

*Glycine* is a paleopolyploid genome, shared with other legumes as an event of first WGD that occurred 59 Mya in Papilionoideae clade. However, the second round of WGD that occurred around 13 Mya was exclusive to the genus *Glycine*, as far as the latest records are concerned, whereby both the species of the genus i.e., *G. max* and *G. soja* share duplicated genome as diplodized tetraploid (Shultz et al., 2006). The two genomes have only 0.31% of SNP variations (Kim et al., 2010). Although studies suggest that intensive domestication events in *G. max* have resulted in massive genomic rearrangements and inclusion of large number of transposable elements in its genetic reservoir (Kim et al., 2010; Sedivy et al., 2017), *G. max* and *G. soja* shares long regions of sequence conservation and instances of chromosomal synteny, along with the gene family expansion. Every node of the *GmBBX* and *GsBBX* cladogram terminated into a leaf node with orthologous *GmBBX* and *GsBBX* (Fig. 3). Each *GmBBX*:*GsBBX* ortholog pair were more closely related to each other than the corresponding *GmBBX* and *GsBBX* paralogs, respectively, justifying the projection of ortholog conjecture.

Apart from ortholog clustering, there were regions of paralog clusters in both the genomes. Most of these genes in the clusters on one chromosome were equidistant to *BBX* gene cluster on another chromosome of the same genome. These gene clusters could be presumably be segmentally duplicated paralogs. Moreover, long stretches of internal synteny between these chromosomal pairs in the two species had a conserved collinear gene order across the two species chromosomes (Cannon and Shoemaker, 2012). As the genes of these gene clusters were phylogenetically closer, we presumed the instances of segmental duplication in the *Glycine* genome.

Furthermore, we detected a chromosomal segment translocation event between the ‘q’ arm in each of the Gm13 and Gs11 chromosomes. The translocation events have been identified in different species of other sub-genus of the genus *Glycine*, which includes a genomic sequence reorganization between Gm13 and Gs11 in *G. max* and *G. soja*, respectively (Zhuang et al., 2022). The phylogenetic analysis revealed the orthologous clustering between Gm13 bearing gene cluster *GmBBX13d-13h* and Gs11 bearing gene cluster *GsBBX11e-11i* (Fig. 4). For instance the gene pair *GmBBX13d:GsBBX11e, GmBBX13e:GsBBX11f* always terminated in the leaf node in a phylogenetic classification. To confirm their relatedness, we first calculated the inter-genic distance between the two consecutive *BBX* genes in each gene cluster located on the two contrasting chromosomes. The *BBX* genes on the ‘q’ arm of Gm13 were equidistant to *BBX* genes on the ‘q’ arm of Gs11. To further ascertain our assumption, we carried out a microsynteny analysis of these regions using GEvo. The similarity results suggested that Gs11 chromosomal segment have been translocated to Gm13. This could have happened due to breakage of the chromosome 11 arm and its reattachment to chromosome 13, leading to the reduction of the arm of Gm11. The translocation event could have possibly occurred after or during the separation of *G. max* from *G. soja*. Hence, based on their evolutionary course, it seems more likely that a translocation event occurred in the Gm11. This altered phenotype caused by chromosomal segment translocation would probably have been favored by the local environment and might have been fixed in the population, through selective breeding, that could have resulted in desirable traits. However, as *G. max* evolved later to *G. soja*, there lies a higher probability of Gm13 gaining the translocated segment, rather than Gs11 losing the segment. Overall, these findings suggest an ancient segmental duplication leading to formation of paralogous genes in *Glycine*.

Eukaryotic duplications have resulted more likely in an autochthonous manner (Gogarten and Olendzenski, 1999). There has been substantial evidence that suggests the conservation of ortholog gene order between species attributed by their conserved long array of DNA sequence organization. This organization pattern contributed by their last common ancestor with sequence conservation after the speciation event, is called conserved linkages (Frazer et al., 2003). Lineage-specific gene gain and loss are frequently associated with *G. max* (Kofsky et al., 2018). With patches of internal synteny between chromosome Gm04 and Gm06, and Gs04 and Gs06, instances of segmental duplication in *G. max*, and *G. soja*, respectively have been clearly identified (Cannon and Shoemaker 2012). The microsyntenic genomic region in and around *BBX* genes in their corresponding chromosomal pairs, possibly suggests that these conserved linkages could have resulted due to segmental duplications, more likely, that it would have occurred in the *Glycine* ancestor. Moreover, there seems to be an insignificant reshuffling of the *BBX* genes after the split of *G. max* and *G. soja*. As the two genomes generate viable fertile hybrids, and differ only by a reciprocal translocation or by a paracentric inversion (Singh and Hymowitz 1988), they are likely to have shared similar genome. Following the identification of incidences of segmental duplication in the *GmBBX* and *GsBBX* gene family, and instances of shared genome synteny, we can suggest that the *BBX* members in each genome are a case of out-paralogs and not in-paralogs, originating from duplication event in the *Glycine* ancestral sequence. Genes duplicated through WGD tend to show lower expressional variation than the duplicated genes arising through tandem or segmental duplication (Casneuf et al., 2006; Rodgers-Melnick et al., 2012).

Conclusively, we can propose that although both the gene families show species-specific divergence among the paralogs, and there has been comparatively a lower rate of *Glycine*-specific divergence with respect to the *BBX* gene family. This observation additionally suggests that there was probably no gene duplication (or as we have not detected) after the speciation event in the *Glycine* sub-genus *soja*. As the *Glycine* genome underwent two rounds of WGD, such lineage-specific duplication can be only ascertained when more species of the genus *Glycine* and probably the sub-genus *glycine* is sequenced. Sharing equal number of *BBX* family members and the orthologous similarity is suggestive of that the *BBX* gene family diverged before the speciation of the genus *Glycine*, hence the members could be referred to as out-paralogs. However, it would be interesting to note that, the Papilionoideae clade underwent first WGD before the split of several other economically important legumes like *Cicer arietinum, Lotus japonicus, Cajanus cajan, Phaseolus vulgaris, Vigna radiata*, etc., we can expect these legumes to carry almost half the number i.e., around 20 to 30 of *BBX* genes as that of *Glycine* genus, depending upon the species-specific gain/loss in *BBX* gene family (Jin et al., 2020; Liu et al., 2021).

## Supplementary data

The supplementary data is available online

## Acknowledgement

This work was supported by the STARS-Ministry of Human Resource and Development (grant number MHRD/BIO/2019085).

