## Supplementary Fig. S for "Comparative analysis of *Glycine BBX* gene family reveals lineage-specific evolution and expansion": fig 1a.pdf

**sequence1 chr:1-4097928**

Supplementary Fig. S1A. The mVISTA plot of Gm04 and Gm06. The X axis represent Gm04, while Y axis represent Gm06.

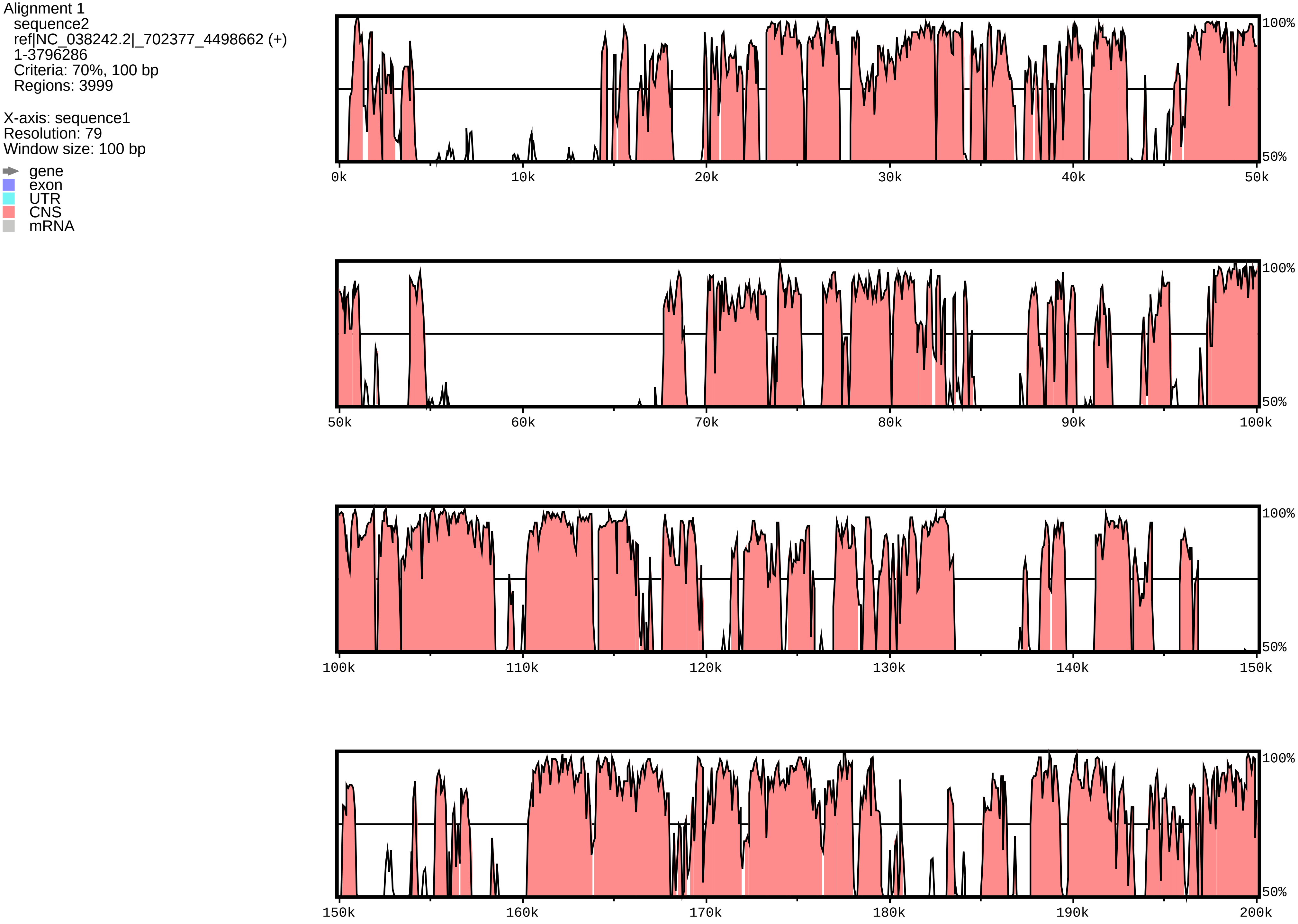

### sequence1 chr:1-4097928

Alignment 1
sequence2
ref|NC_038242.2|_702377_4498662 (+)
1-3796286
Criteria: 70%, 100 bp
Regions: 3999

X-axis: sequence1
Resolution: 79
Window size: 100 bp

gene
exon
UTR
CNS
mRNA

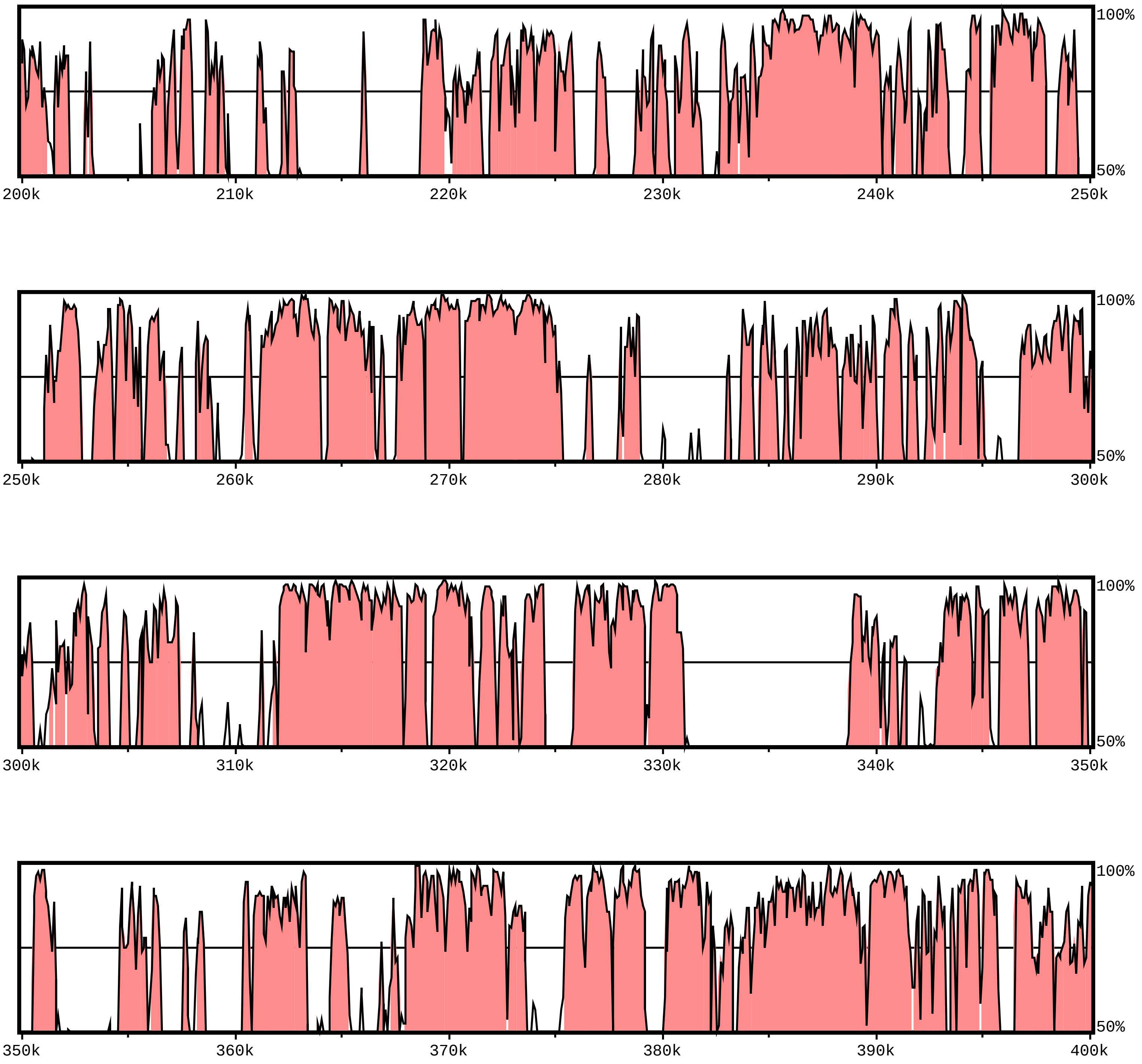

**sequence1 chr:1-4097928**

Alignment 1
sequence2
ref|NC_038242.2|_702377_4498662 (+)
1-3796286
Criteria: 70%, 100 bp
Regions: 3999

X-axis: sequence1
Resolution: 79
Window size: 100 bp

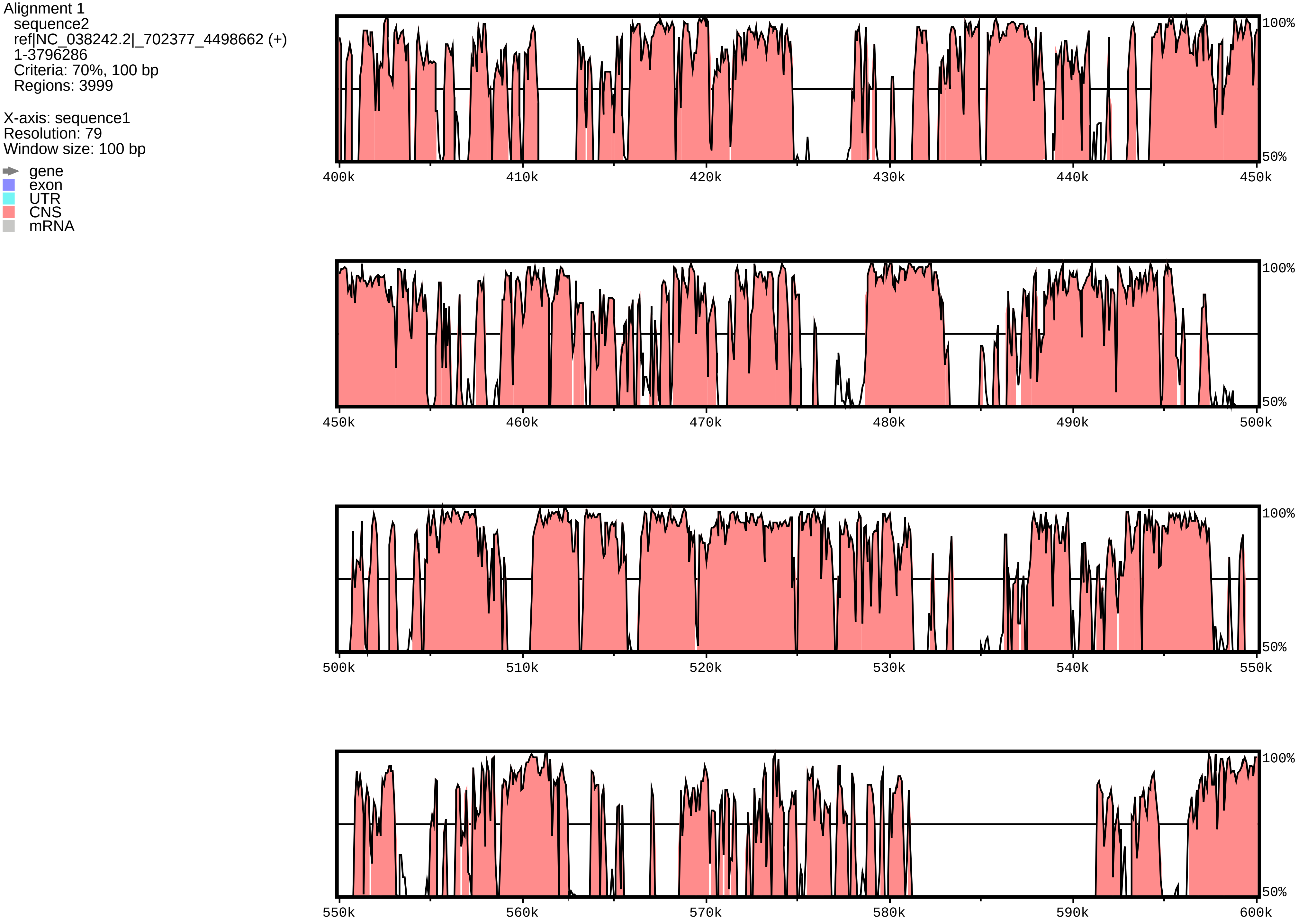

**sequence1 chr:1-4097928**

Alignment 1
sequence2
ref|NC_038242.2|_702377_4498662 (+)
1-3796286
Criteria: 70%, 100 bp
Regions: 3999

X-axis: sequence1
Resolution: 79
Window size: 100 bp

gene
exon
UTR
CNS
mRNA

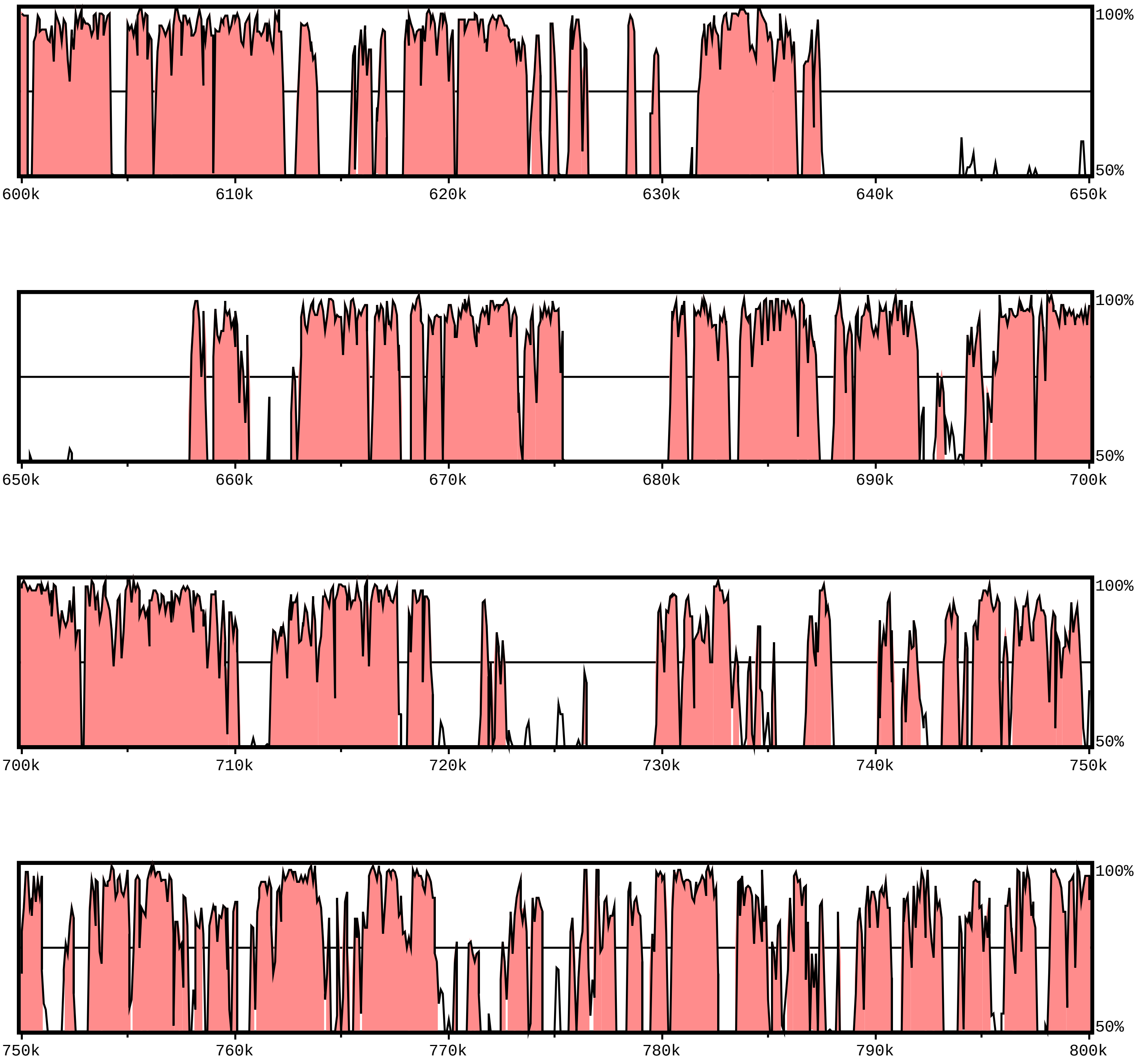

**sequence1 chr:1-4097928**

Alignment 1
sequence2
ref|NC_038242.2|_702377_4498662 (+)
1-3796286
Criteria: 70%, 100 bp
Regions: 3999

X-axis: sequence1
Resolution: 79
Window size: 100 bp

gene
exon
UTR
CNS
mRNA

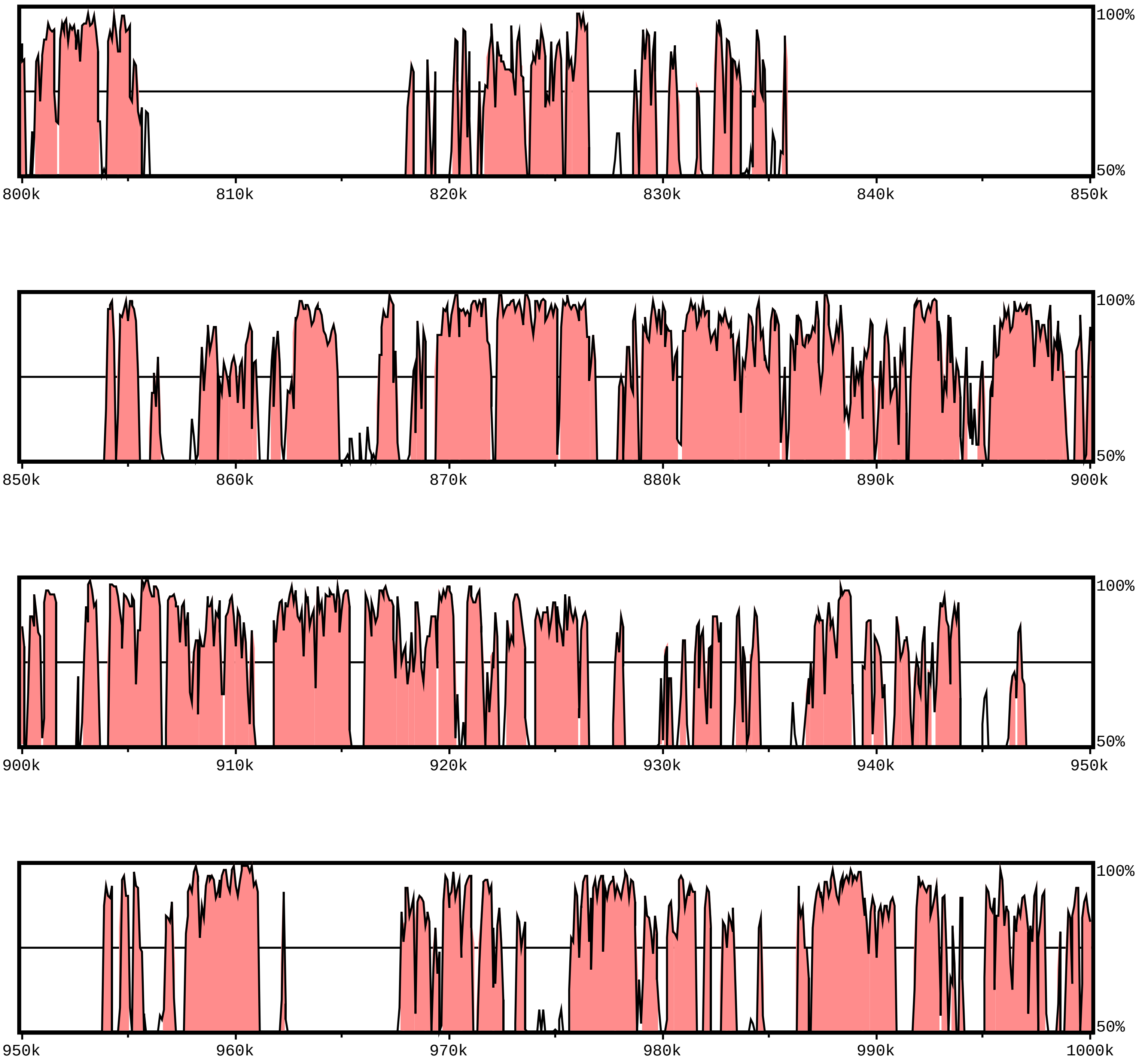

**sequence1 chr:1-4097928**

Alignment 1
sequence2
ref|NC_038242.2|_702377_4498662 (+)
1-3796286
Criteria: 70%, 100 bp
Regions: 3999

X-axis: sequence1
Resolution: 79
Window size: 100 bp

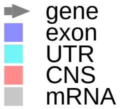


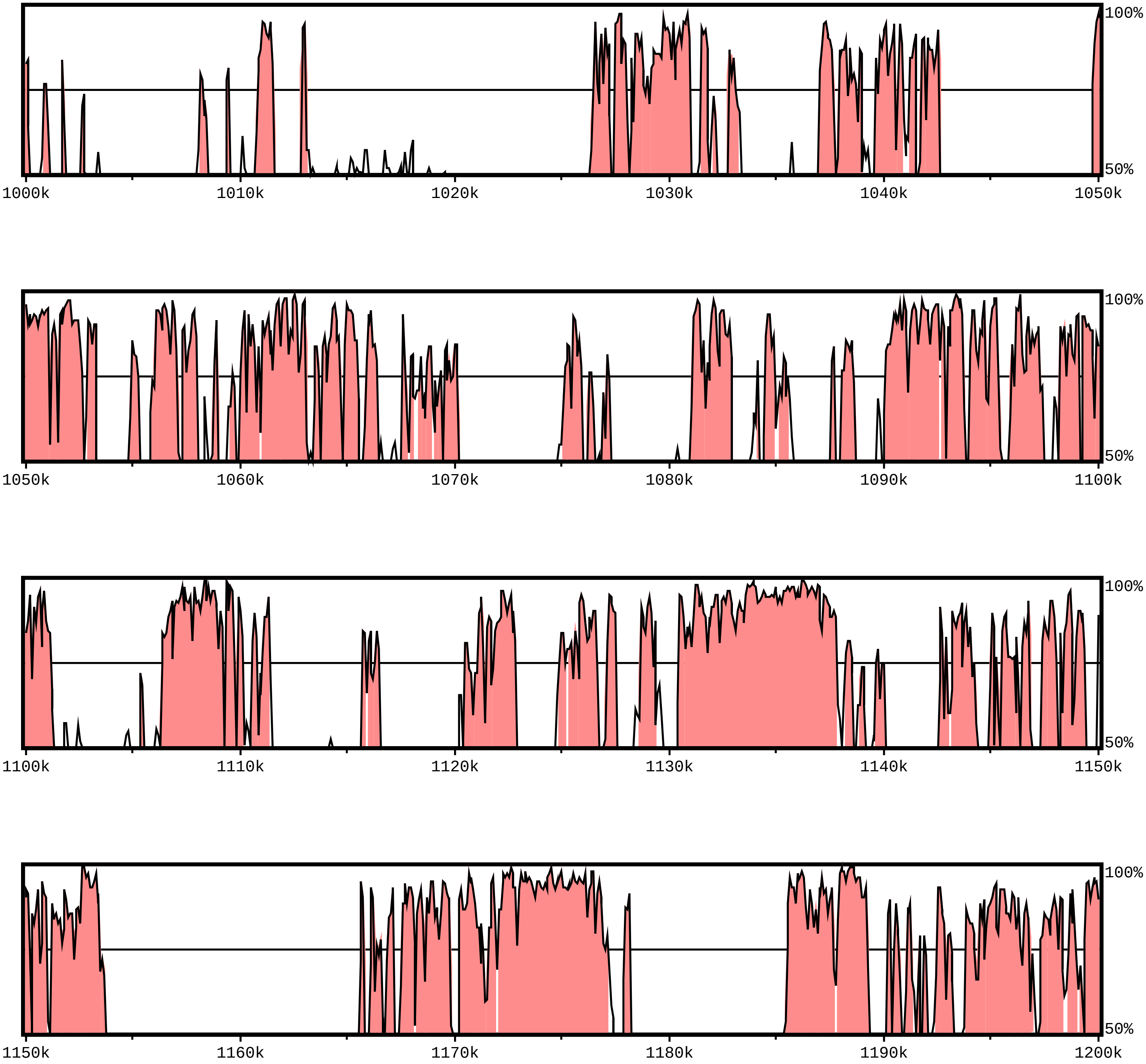

**sequence1 chr:1-4097928**

Alignment 1
sequence2
ref|NC_038242.2|_702377_4498662 (+)
1-3796286
Criteria: 70%, 100 bp
Regions: 3999

X-axis: sequence1
Resolution: 79
Window size: 100 bp

gene
exon
UTR
CNS
mRNA

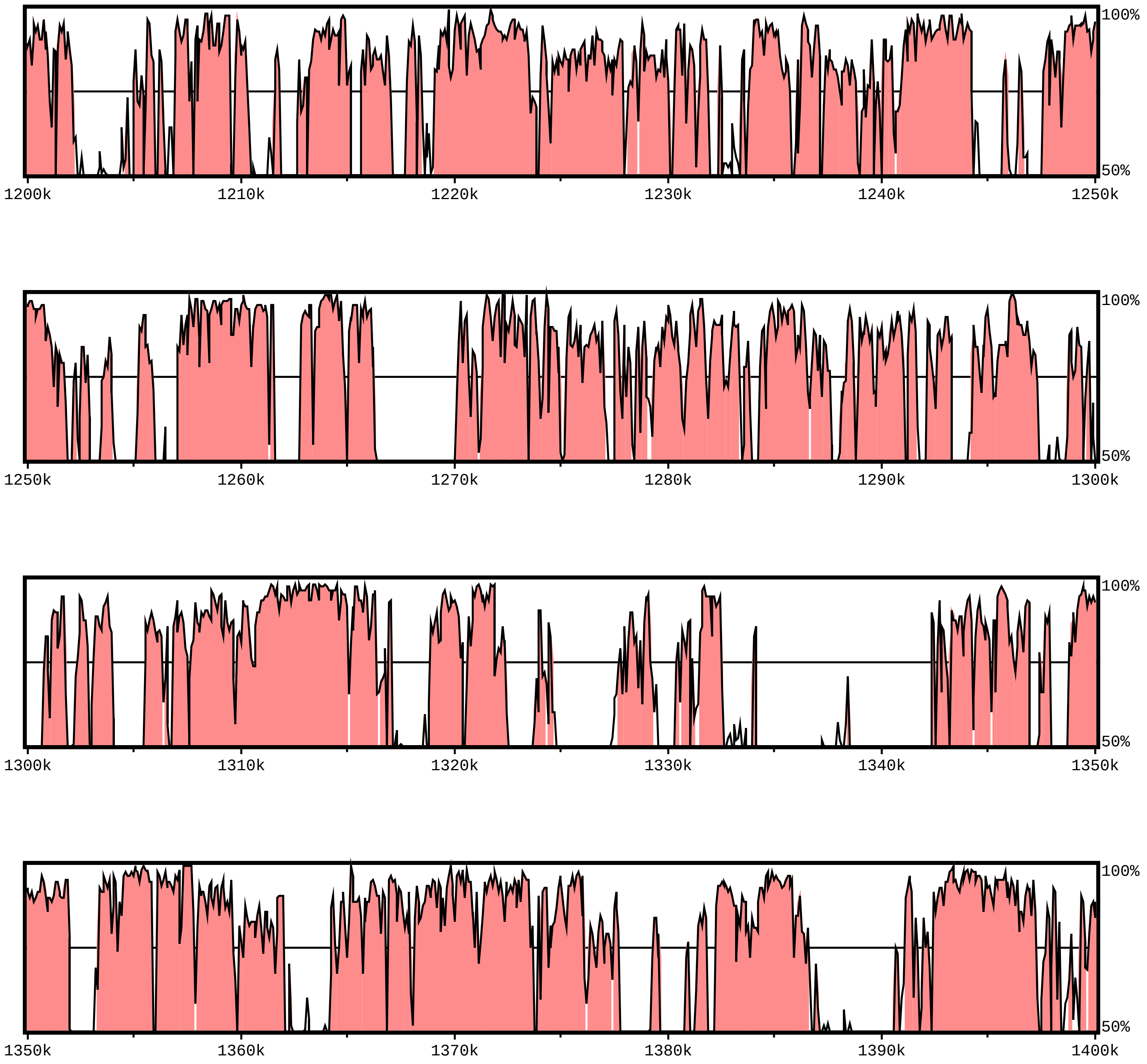

**sequence1 chr:1-4097928**

Alignment 1
sequence2
ref|NC_038242.2|_702377_4498662 (+)
1-3796286
Criteria: 70%, 100 bp
Regions: 3999

X-axis: sequence1
Resolution: 79
Window size: 100 bp

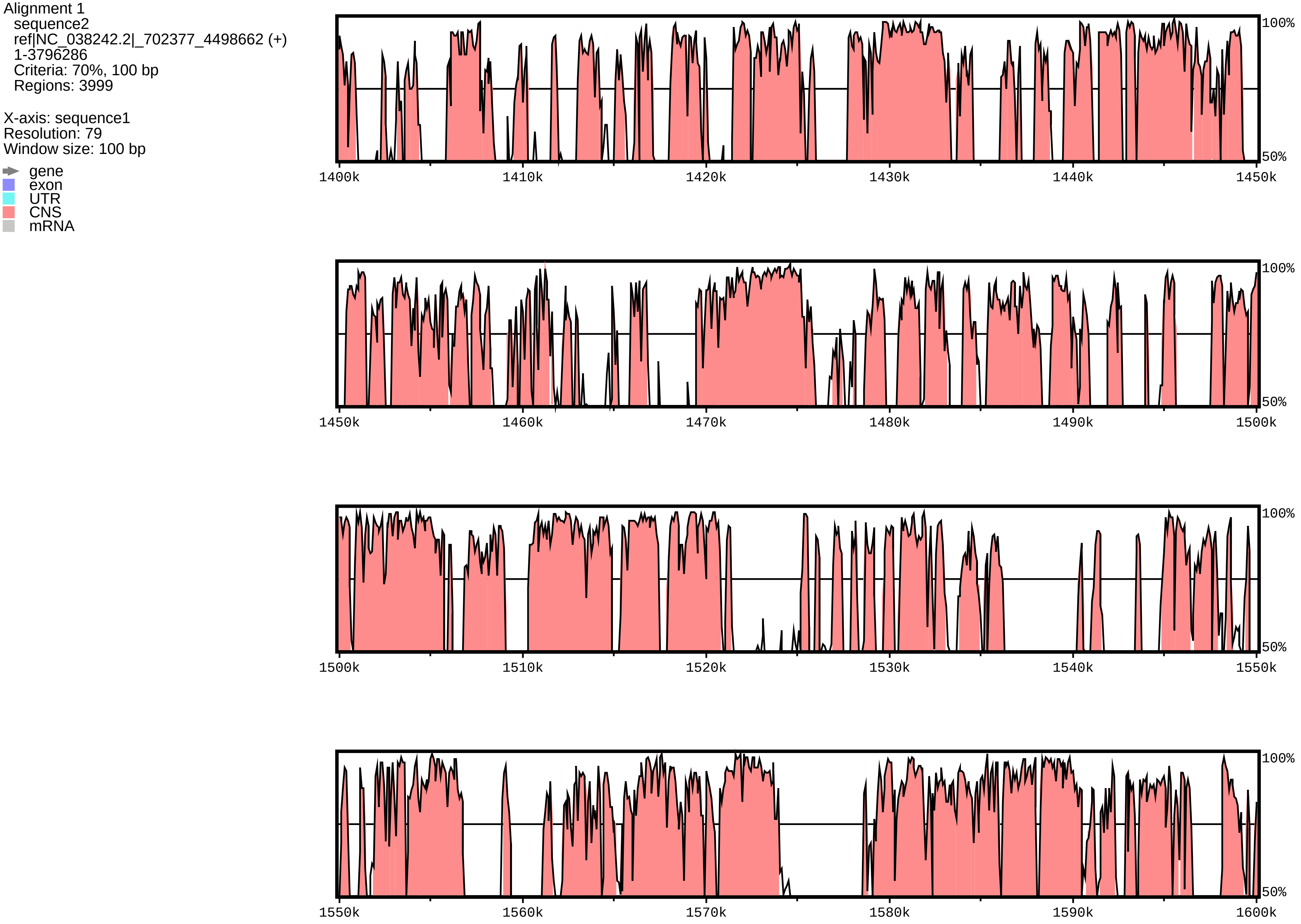

**sequence1 chr:1-4097928**

Alignment 1
sequence2
ref|NC_038242.2|_702377_4498662 (+)
1-3796286
Criteria: 70%, 100 bp
Regions: 3999

X-axis: sequence1
Resolution: 79
Window size: 100 bp

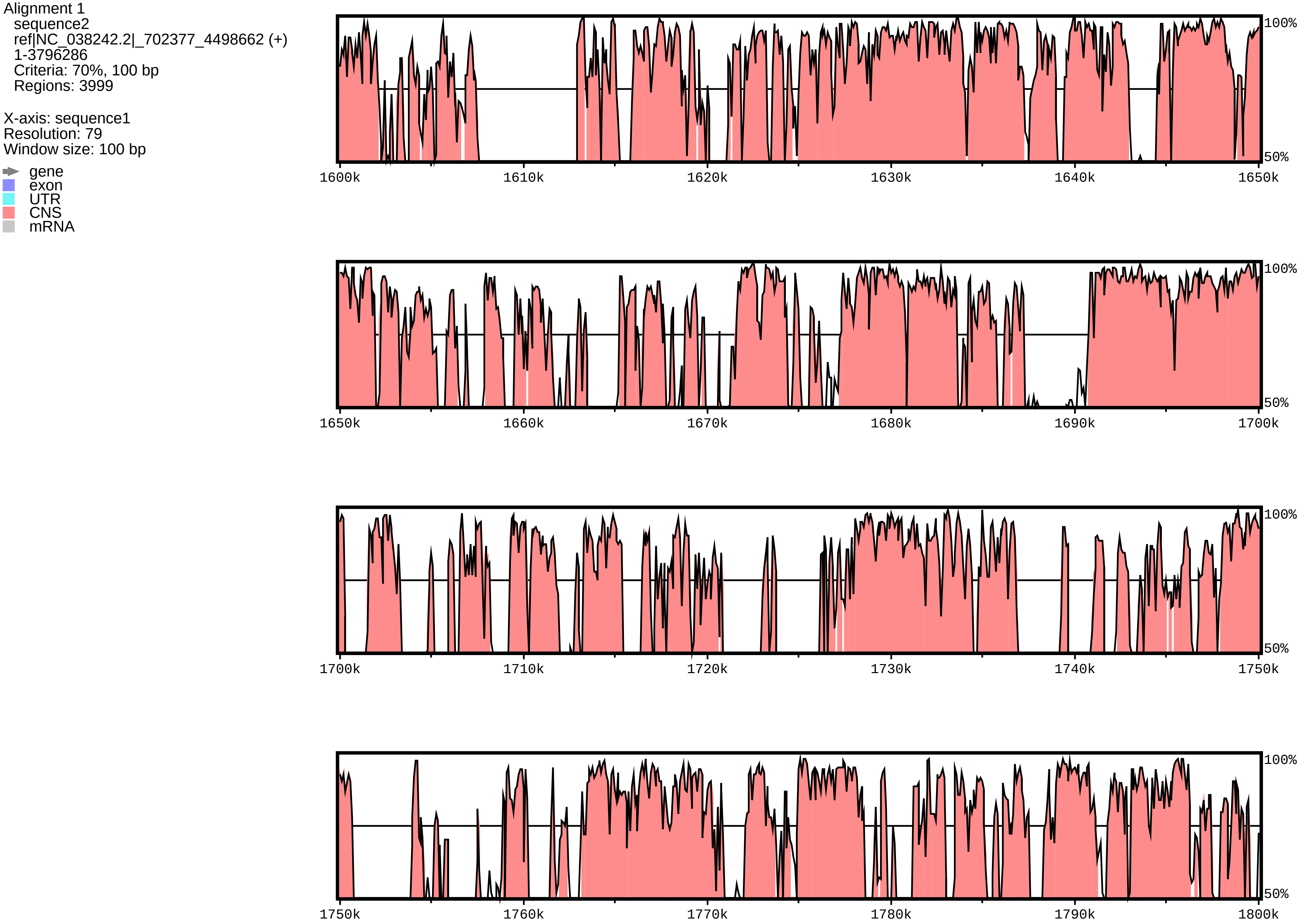

**sequence1 chr:1-4097928**

Alignment 1
sequence2
ref|NC_038242.2|_702377_4498662 (+)
1-3796286
Criteria: 70%, 100 bp
Regions: 3999

X-axis: sequence1
Resolution: 79
Window size: 100 bp

gene
exon
UTR
CNS
mRNA

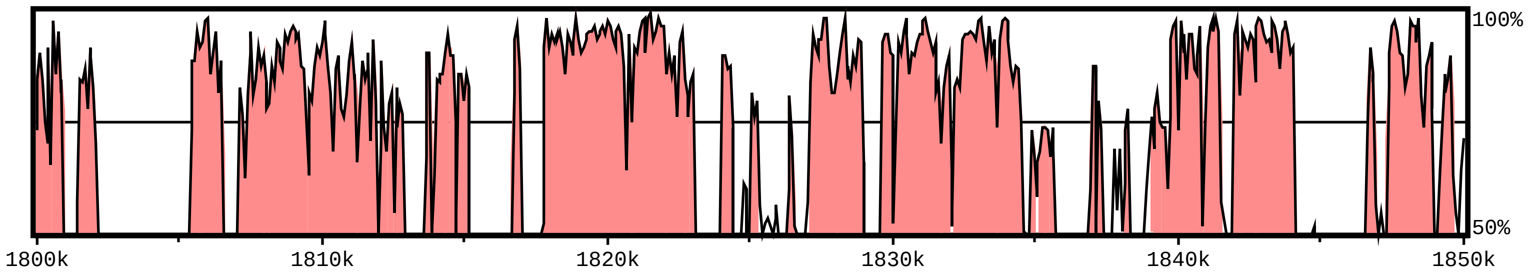


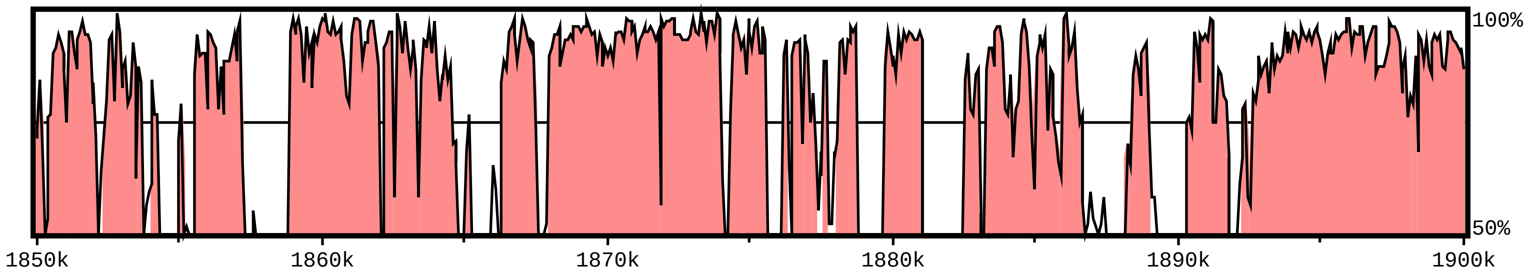


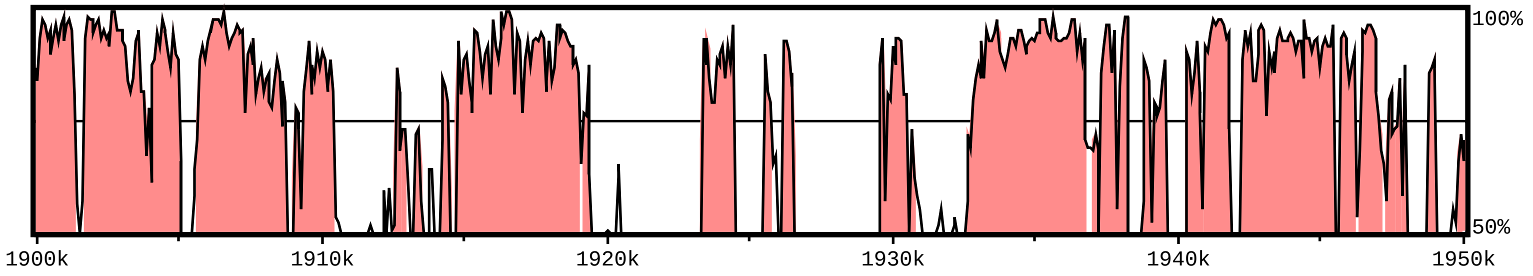


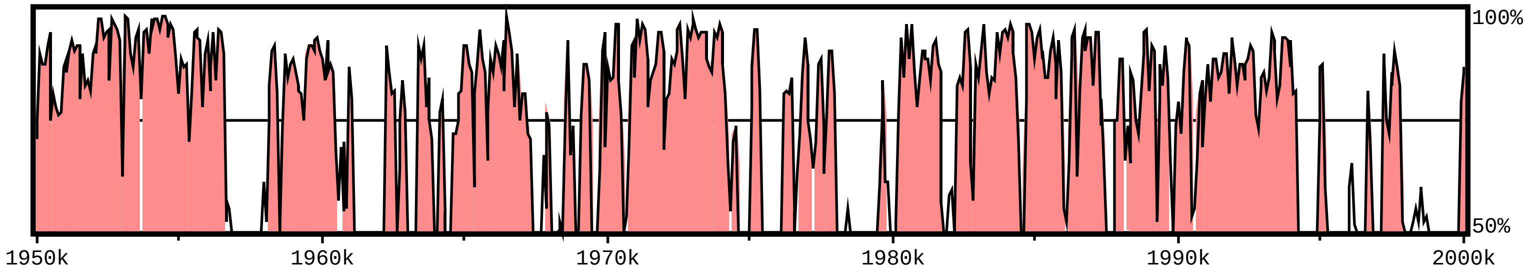

**sequence1 chr:1-4097928**

Alignment 1
sequence2
ref|NC_038242.2|_702377_4498662 (+)
1-3796286
Criteria: 70%, 100 bp
Regions: 3999

X-axis: sequence1
Resolution: 79
Window size: 100 bp

gene
exon
UTR
CNS
mRNA

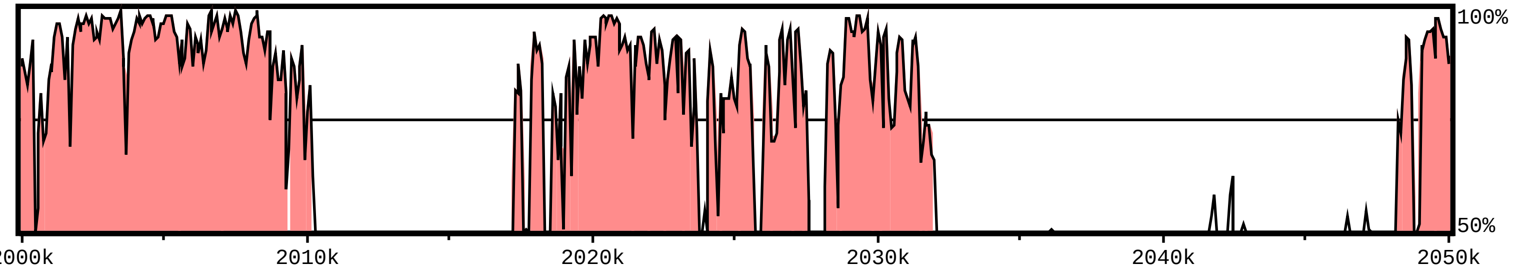


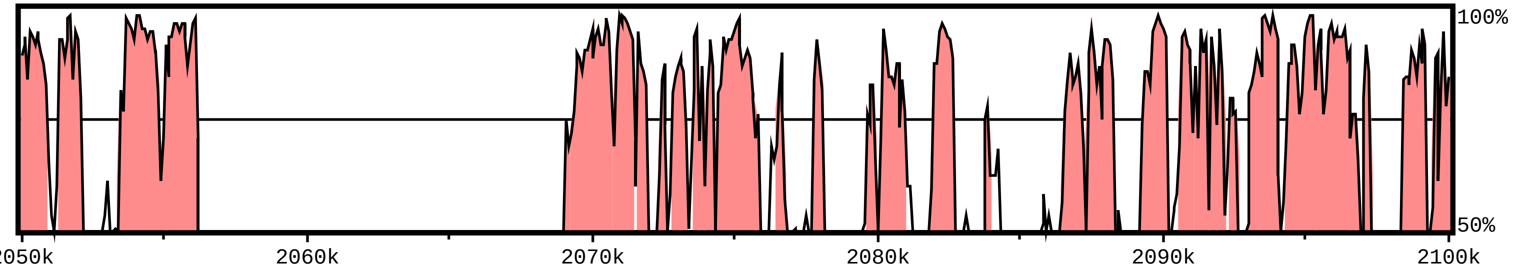


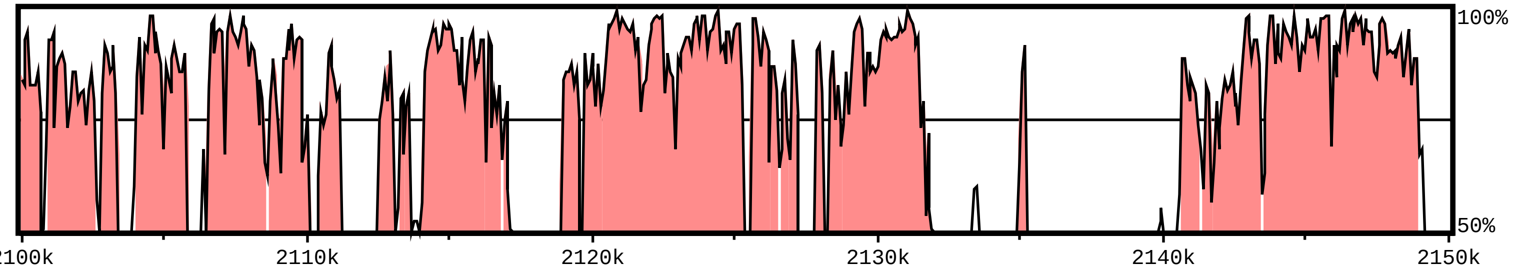


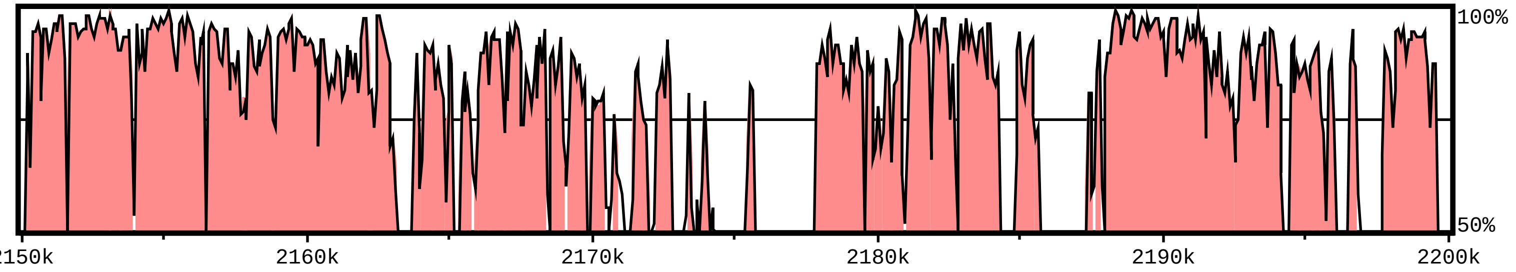

**sequence1 chr:1-4097928**

Alignment 1
sequence2
ref|NC_038242.2|_702377_4498662 (+)
1-3796286
Criteria: 70%, 100 bp
Regions: 3999

X-axis: sequence1
Resolution: 79
Window size: 100 bp

- gene
- exon
- UTR
- CNS
- mRNA

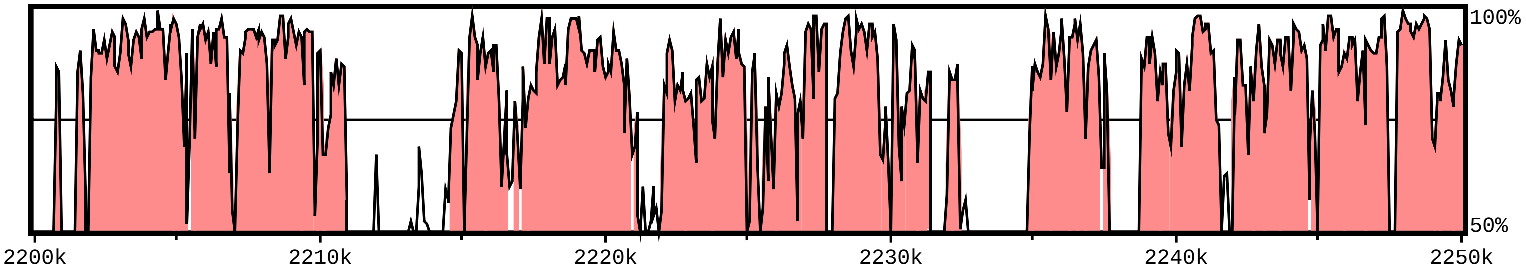


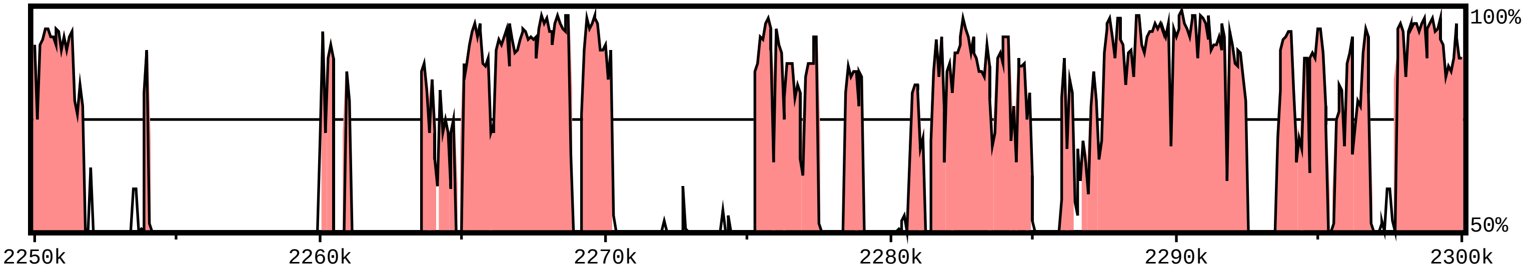


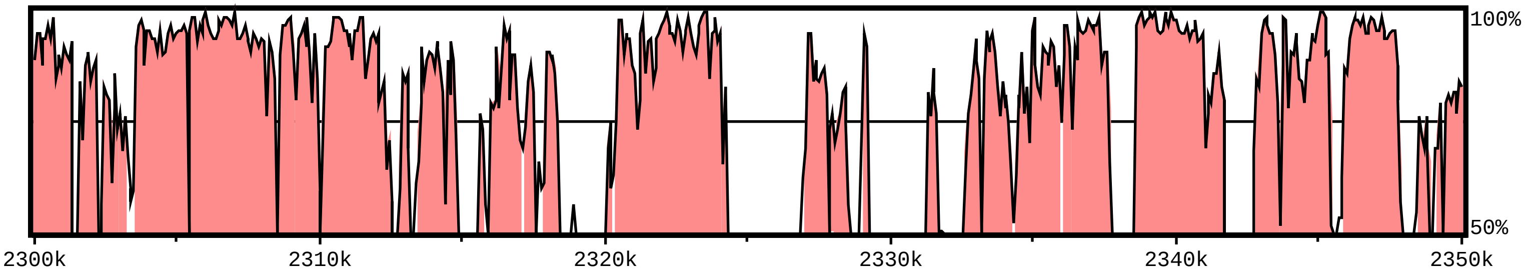


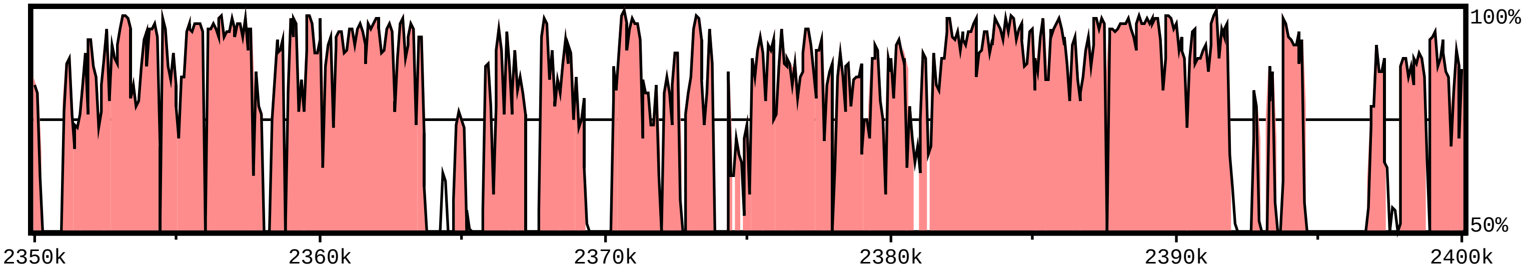

**sequence1 chr:1-4097928**

Alignment 1
sequence2
ref|NC_038242.2|_702377_4498662 (+)
1-3796286
Criteria: 70%, 100 bp
Regions: 3999

X-axis: sequence1
Resolution: 79
Window size: 100 bp

gene
exon
UTR
CNS
mRNA

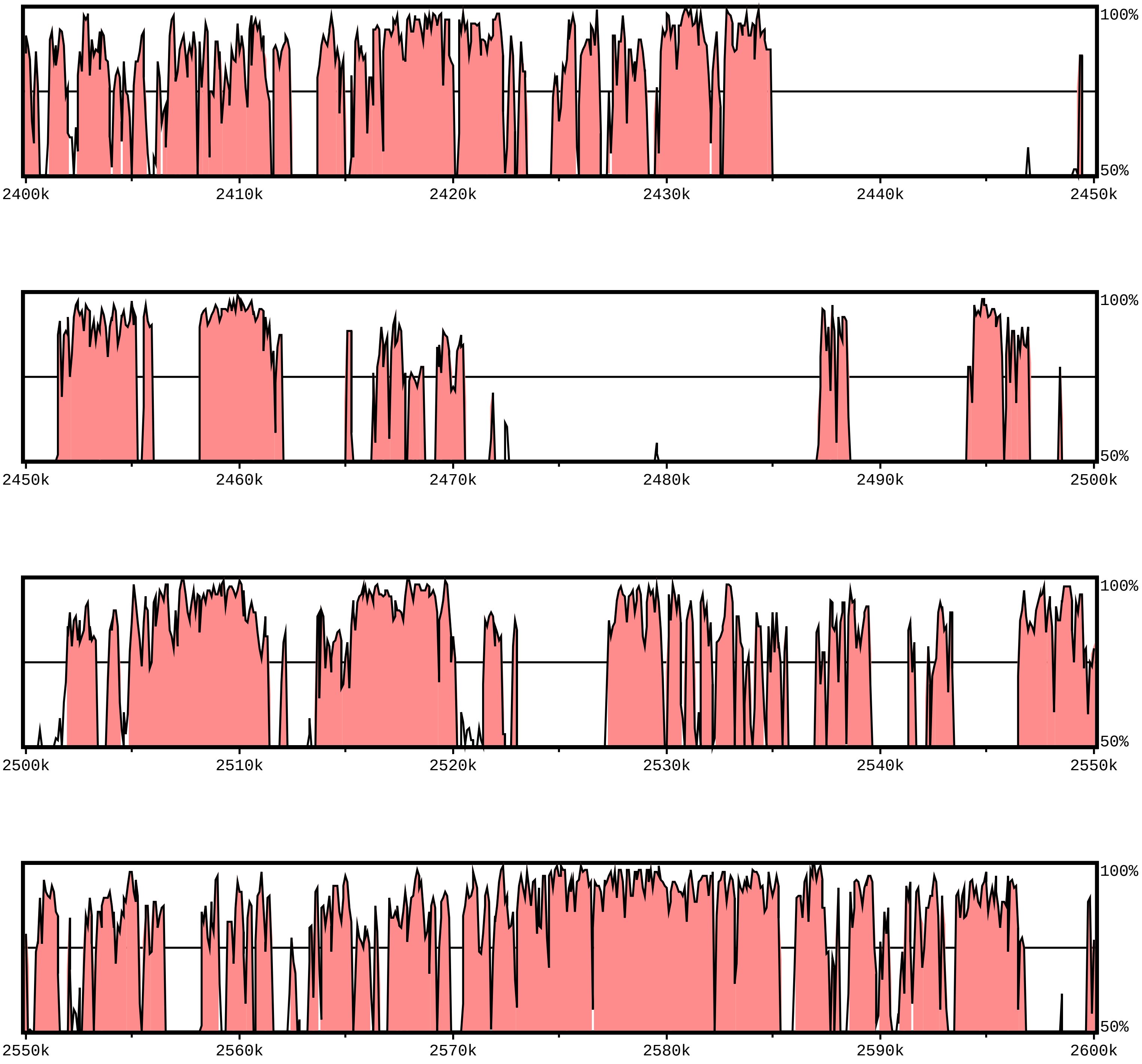

**sequence1 chr:1-4097928**

Alignment 1
sequence2
ref|NC_038242.2|_702377_4498662 (+)
1-3796286
Criteria: 70%, 100 bp
Regions: 3999

X-axis: sequence1
Resolution: 79
Window size: 100 bp

gene
exon
UTR
CNS
mRNA

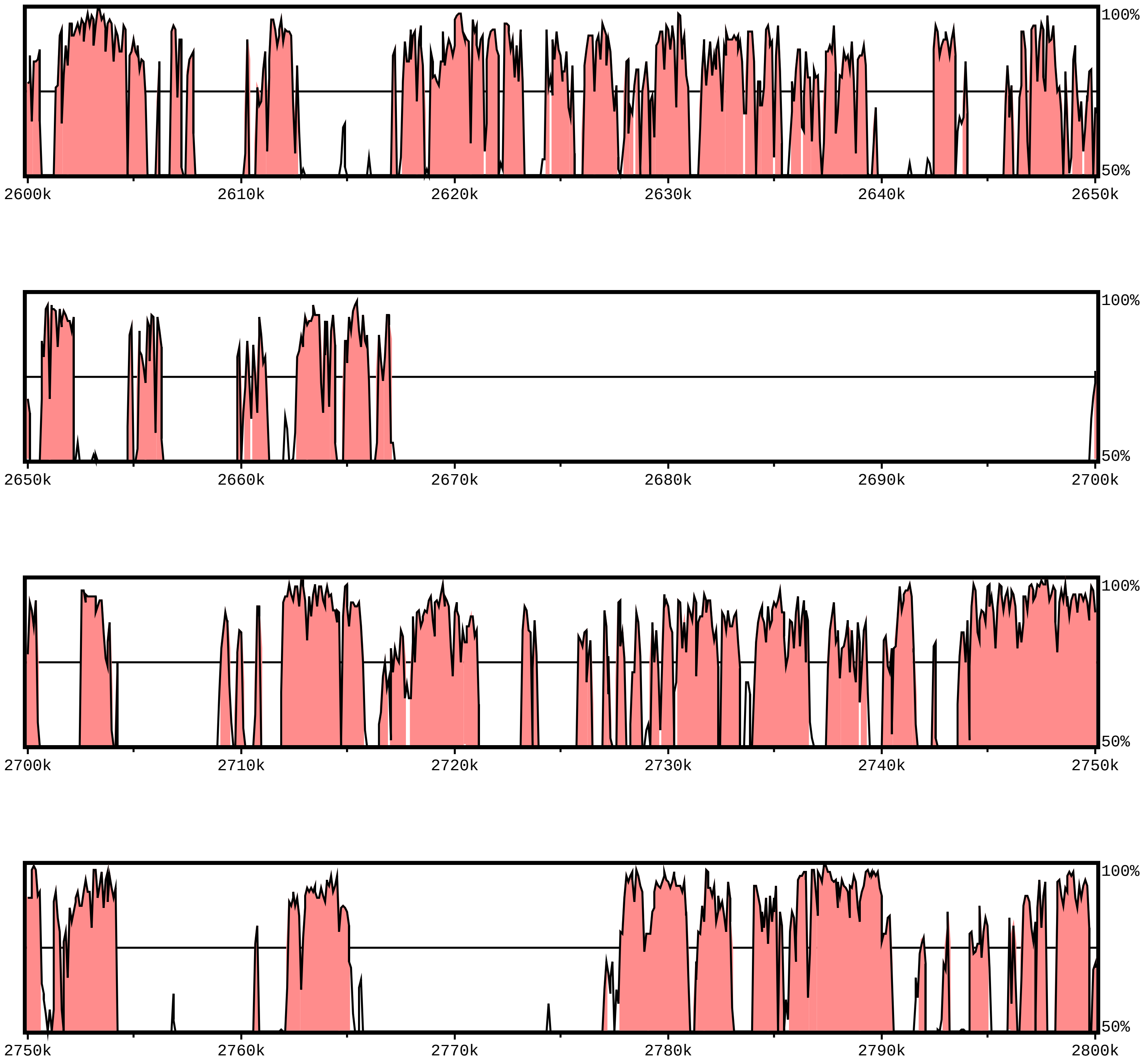

**sequence1 chr:1-4097928**

Alignment 1
sequence2
ref|NC_038242.2|_702377_4498662 (+)
1-3796286
Criteria: 70%, 100 bp
Regions: 3999

X-axis: sequence1
Resolution: 79
Window size: 100 bp

gene
exon
UTR
CNS
mRNA

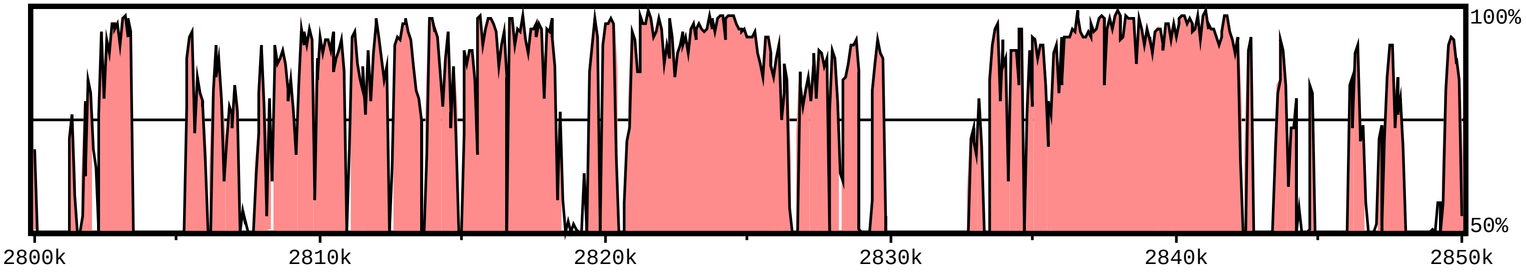


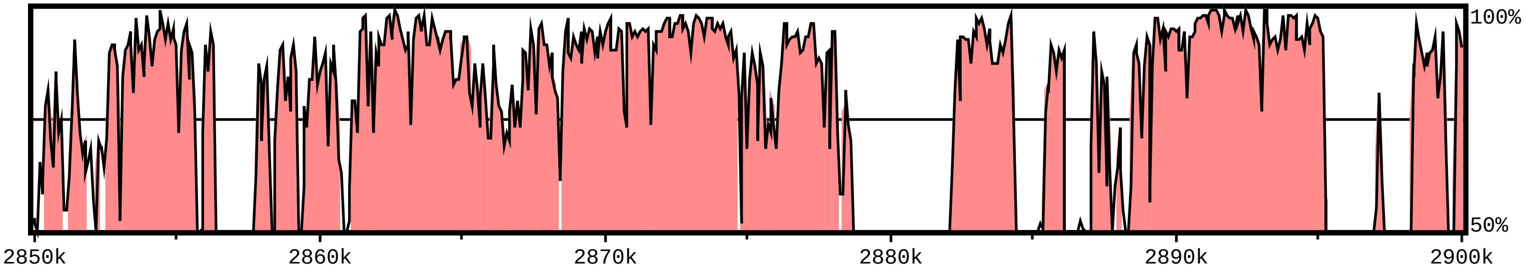


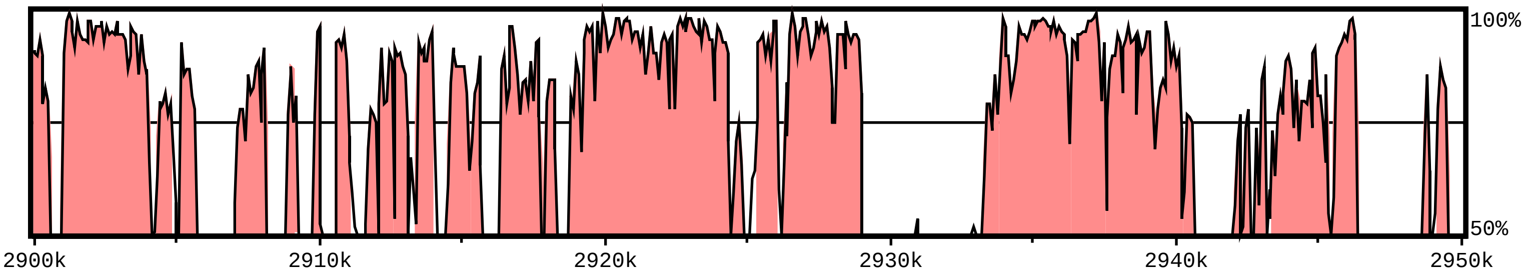


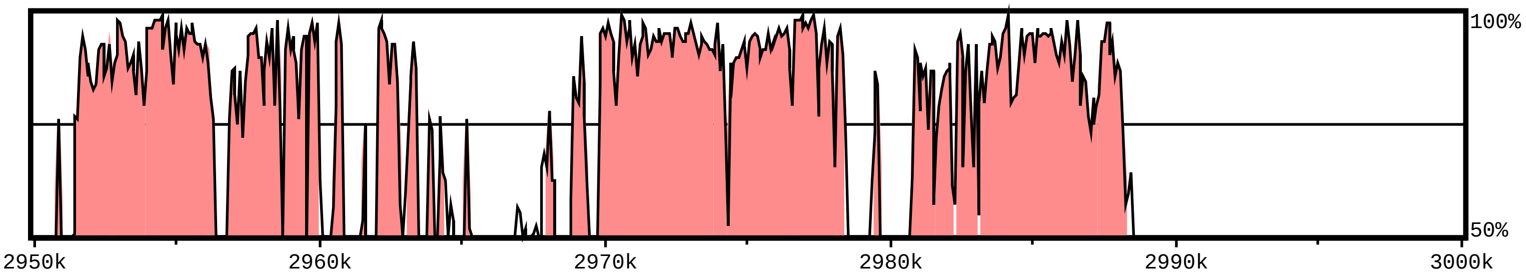

**sequence1 chr:1-4097928**

Alignment 1
sequence2
ref|NC_038242.2|_702377_4498662 (+)
1-3796286
Criteria: 70%, 100 bp
Regions: 3999

X-axis: sequence1
Resolution: 79
Window size: 100 bp

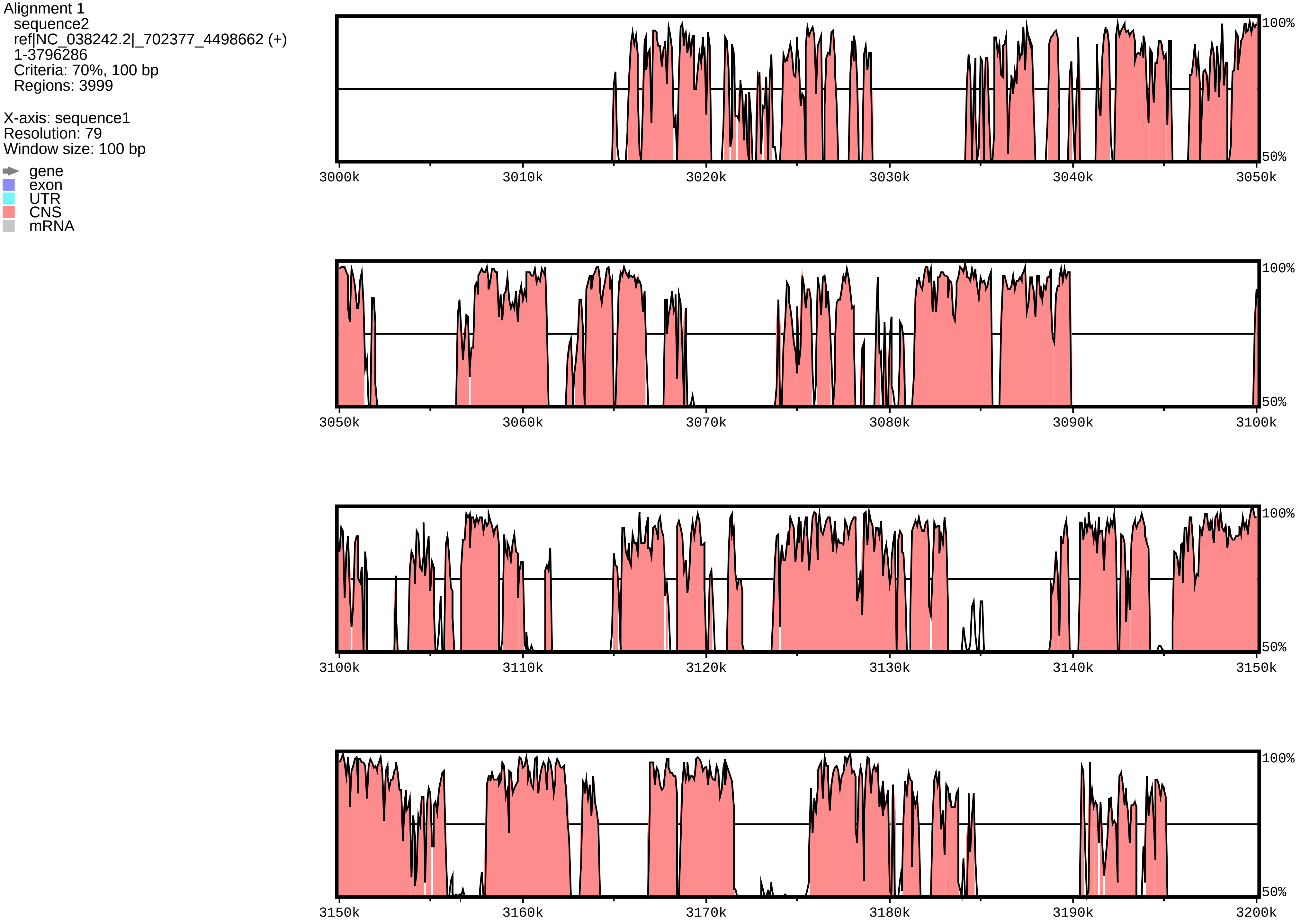

**sequence1 chr:1-4097928**

Alignment 1
sequence2
ref|NC_038242.2|_702377_4498662 (+)
1-3796286
Criteria: 70%, 100 bp
Regions: 3999

X-axis: sequence1
Resolution: 79
Window size: 100 bp

gene
exon
UTR
CNS
mRNA

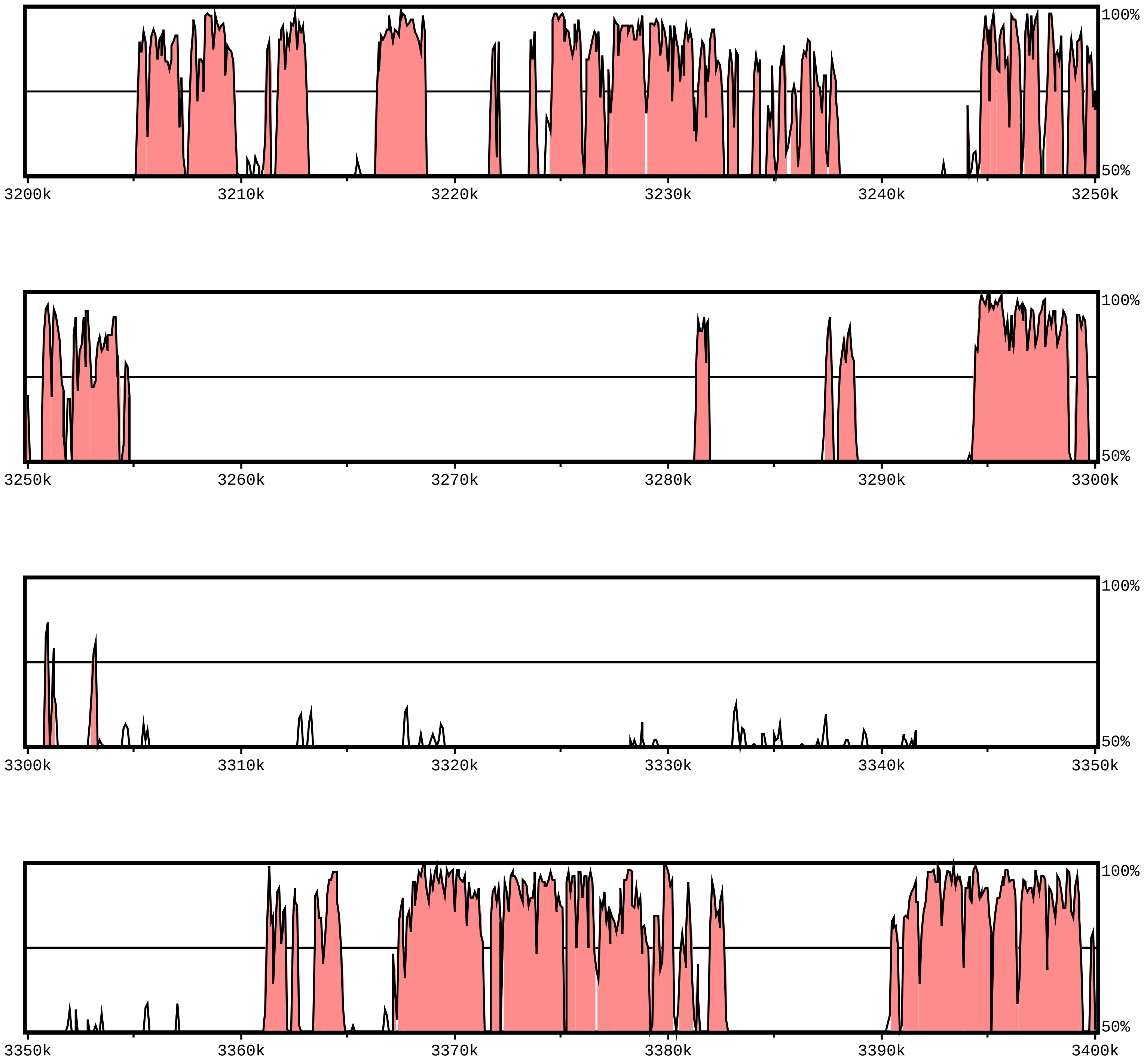

### sequence1 chr:1-4097928

Alignment 1
sequence2
ref|NC_038242.2|_702377_4498662 (+)
1-3796286
Criteria: 70%, 100 bp
Regions: 3999

X-axis: sequence1
Resolution: 79
Window size: 100 bp

- gene
- exon
- UTR
- CNS
- mRNA

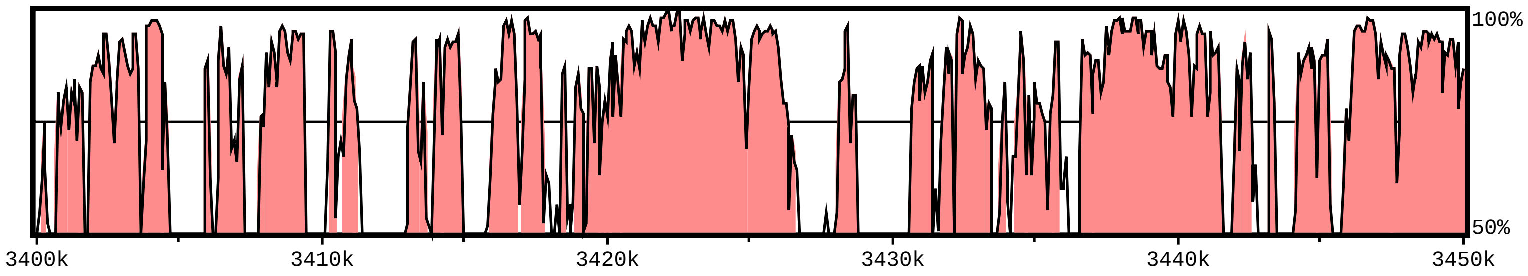


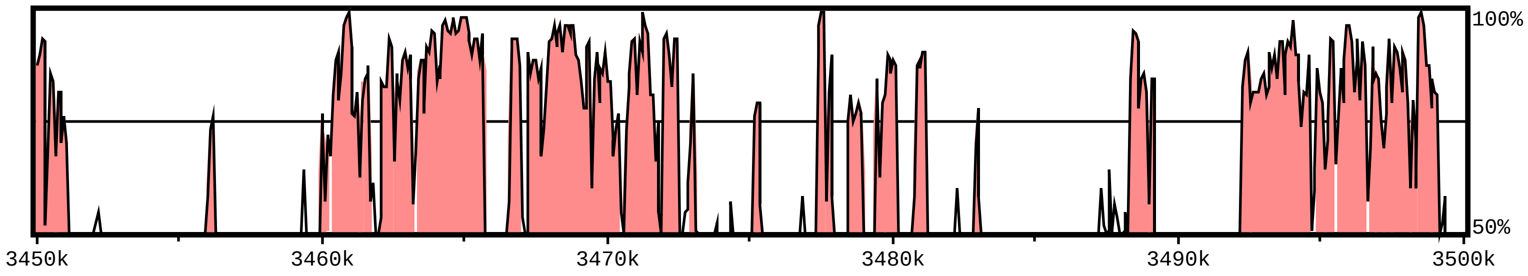


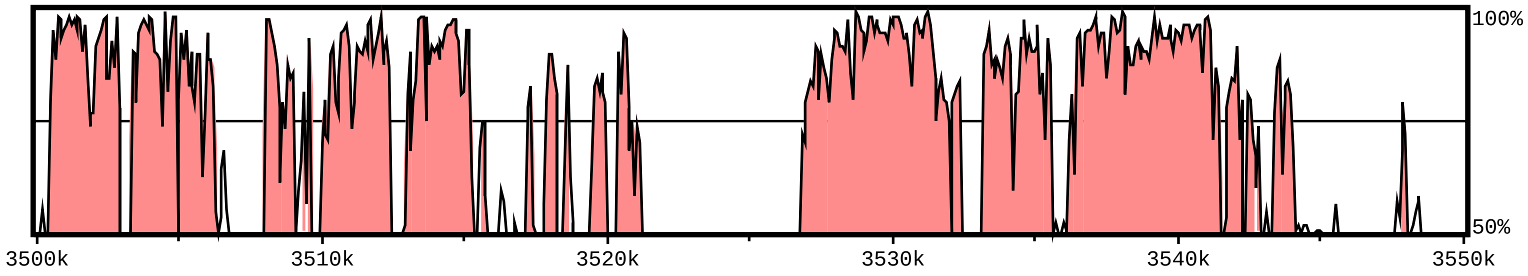


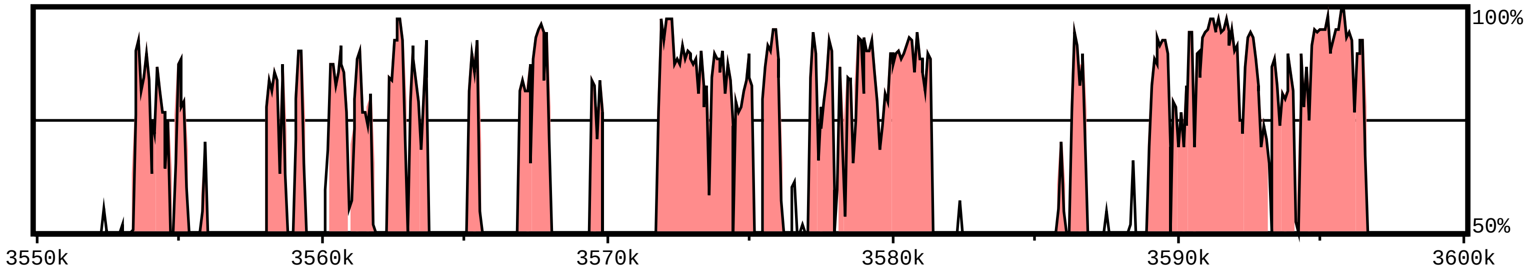

**sequence1 chr:1-4097928**

Alignment 1
sequence2
ref|NC_038242.2|_702377_4498662 (+)
1-3796286
Criteria: 70%, 100 bp
Regions: 3999

X-axis: sequence1
Resolution: 79
Window size: 100 bp

gene
exon
UTR
CNS
mRNA

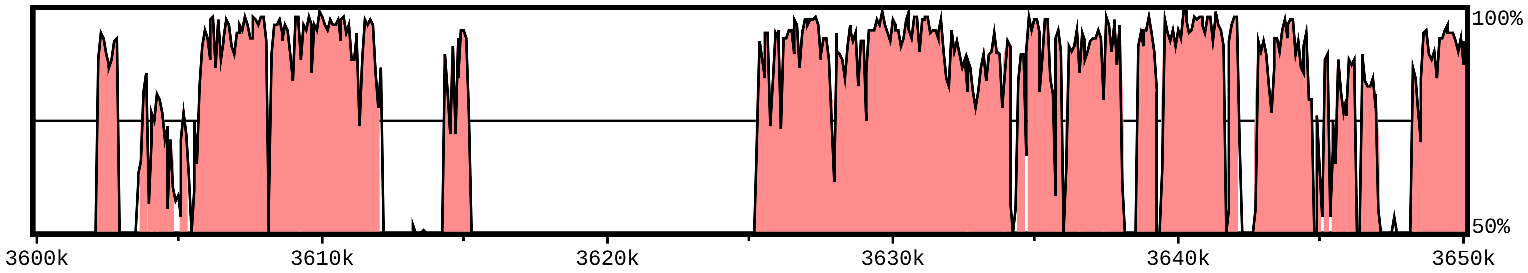


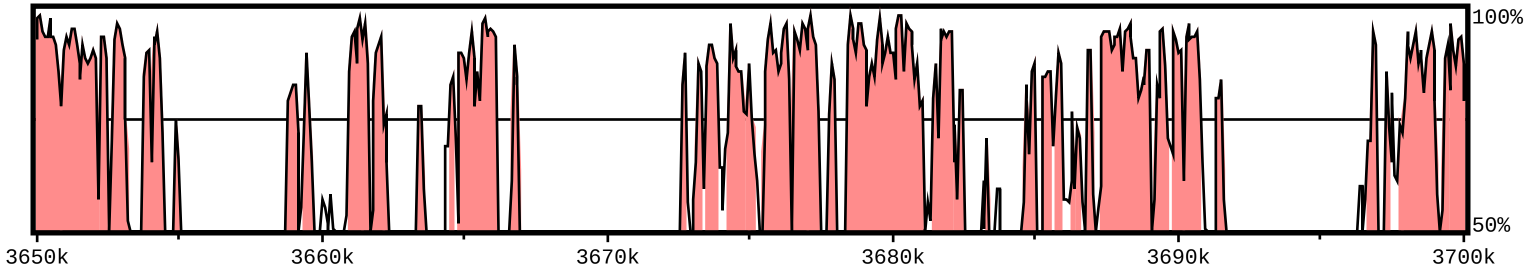


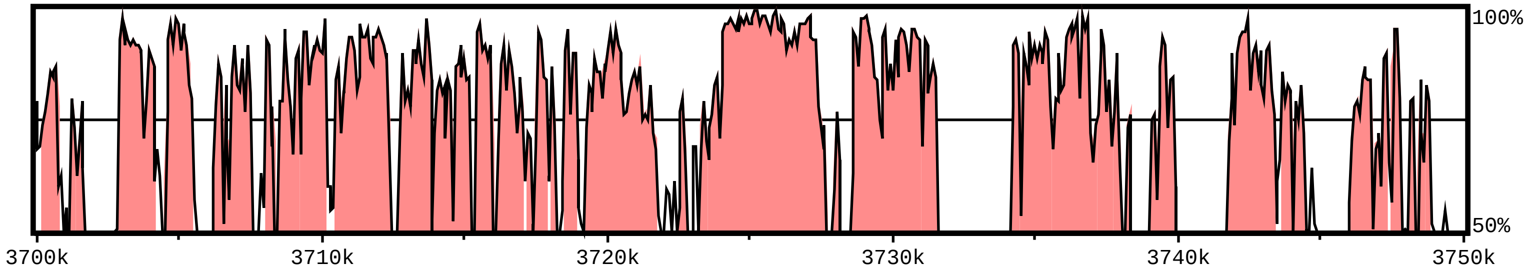


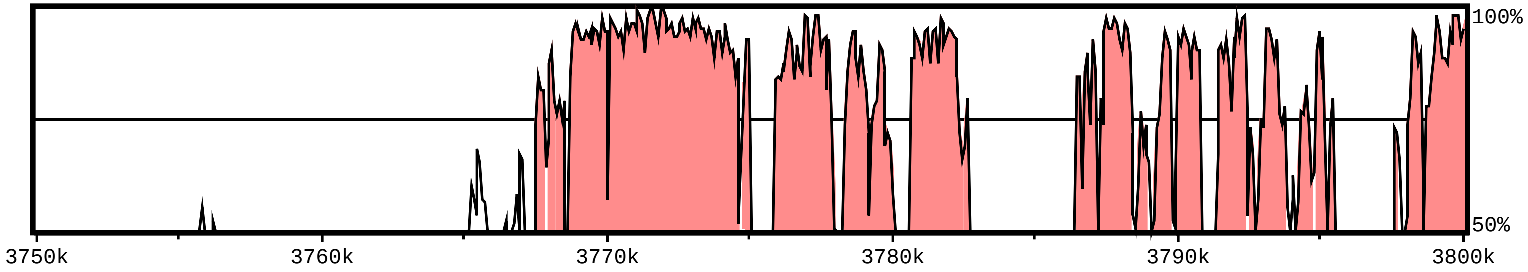

**sequence1 chr:1-4097928**

Alignment 1
sequence2
ref|NC_038242.2|_702377_4498662 (+)
1-3796286
Criteria: 70%, 100 bp
Regions: 3999

X-axis: sequence1
Resolution: 79
Window size: 100 bp

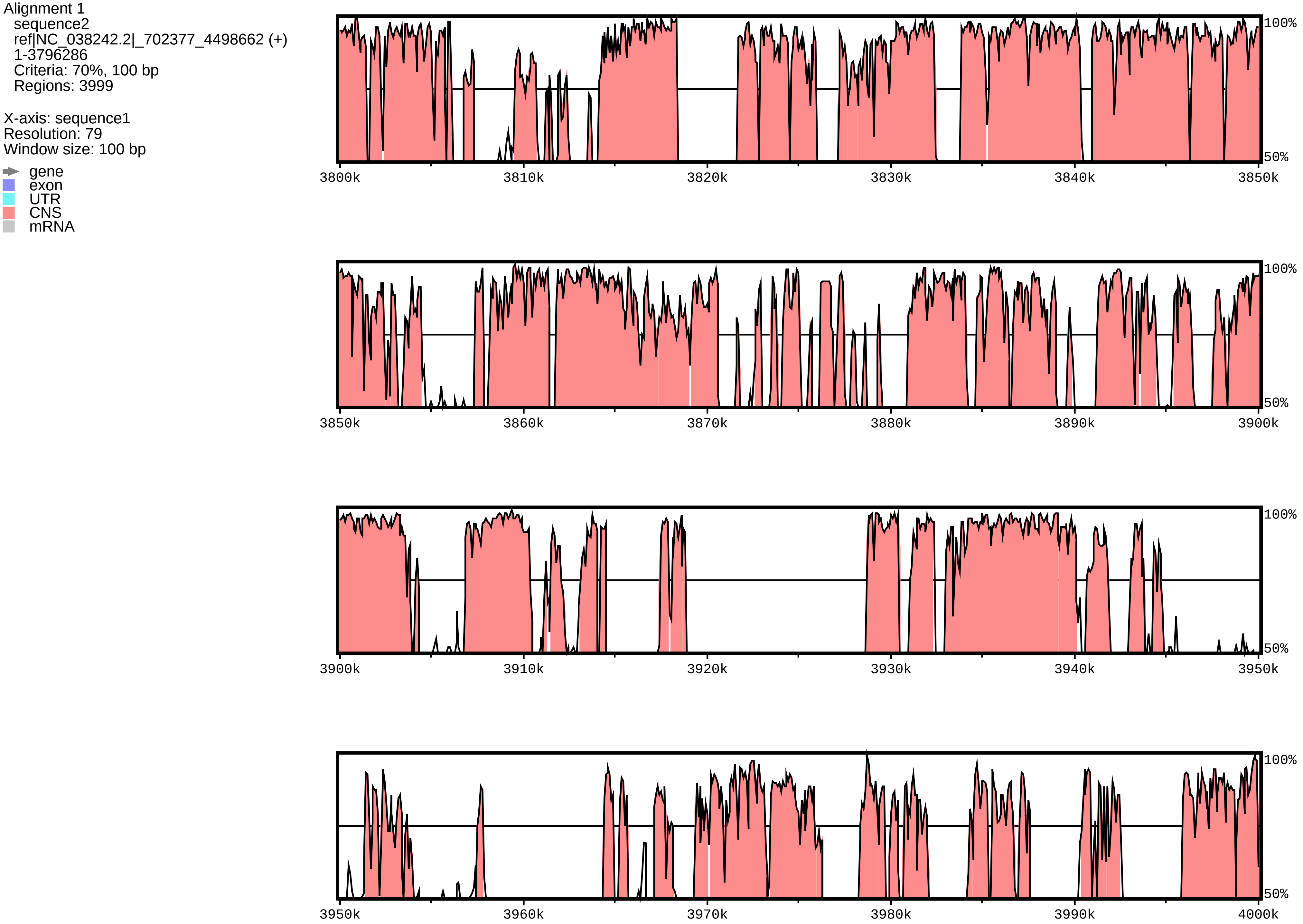

**sequence1 chr:1-4097928**

Alignment 1
sequence2
ref|NC_038242.2|_702377_4498662 (+)
1-3796286
Criteria: 70%, 100 bp
Regions: 3999

X-axis: sequence1
Resolution: 79
Window size: 100 bp

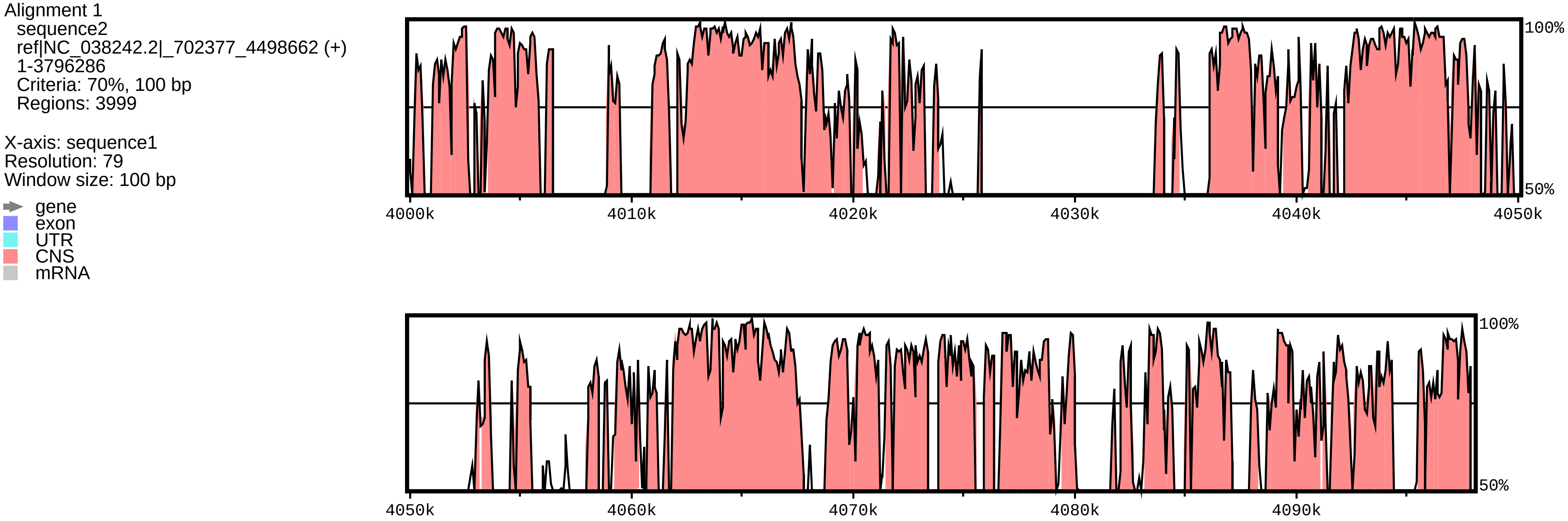
