## Supplementary Fig. S for "Comparative analysis of *Glycine BBX* gene family reveals lineage-specific evolution and expansion": fig 1b.pdf

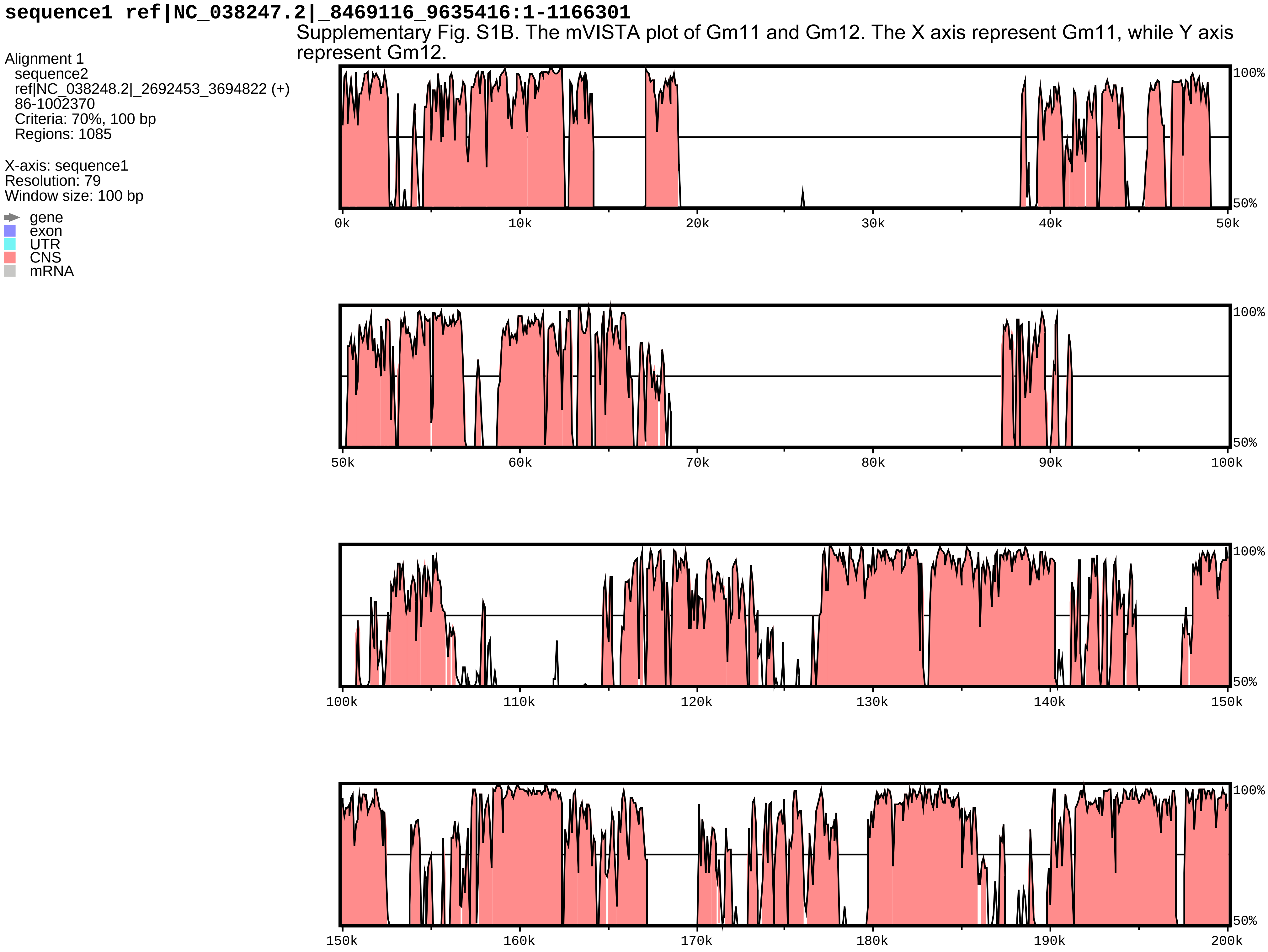


Supplementary Fig. S1B. The mVISTA plot of Gm11 and Gm12. The X axis represent Gm11, while Y axis represent Gm12.

**sequence1 ref|NC_038247.2|_8469116_9635416:1-1166301**

Alignment 1
sequence2
ref|NC_038248.2|_2692453_3694822 (+)
86-1002370
Criteria: 70%, 100 bp
Regions: 1085

X-axis: sequence1
Resolution: 79
Window size: 100 bp

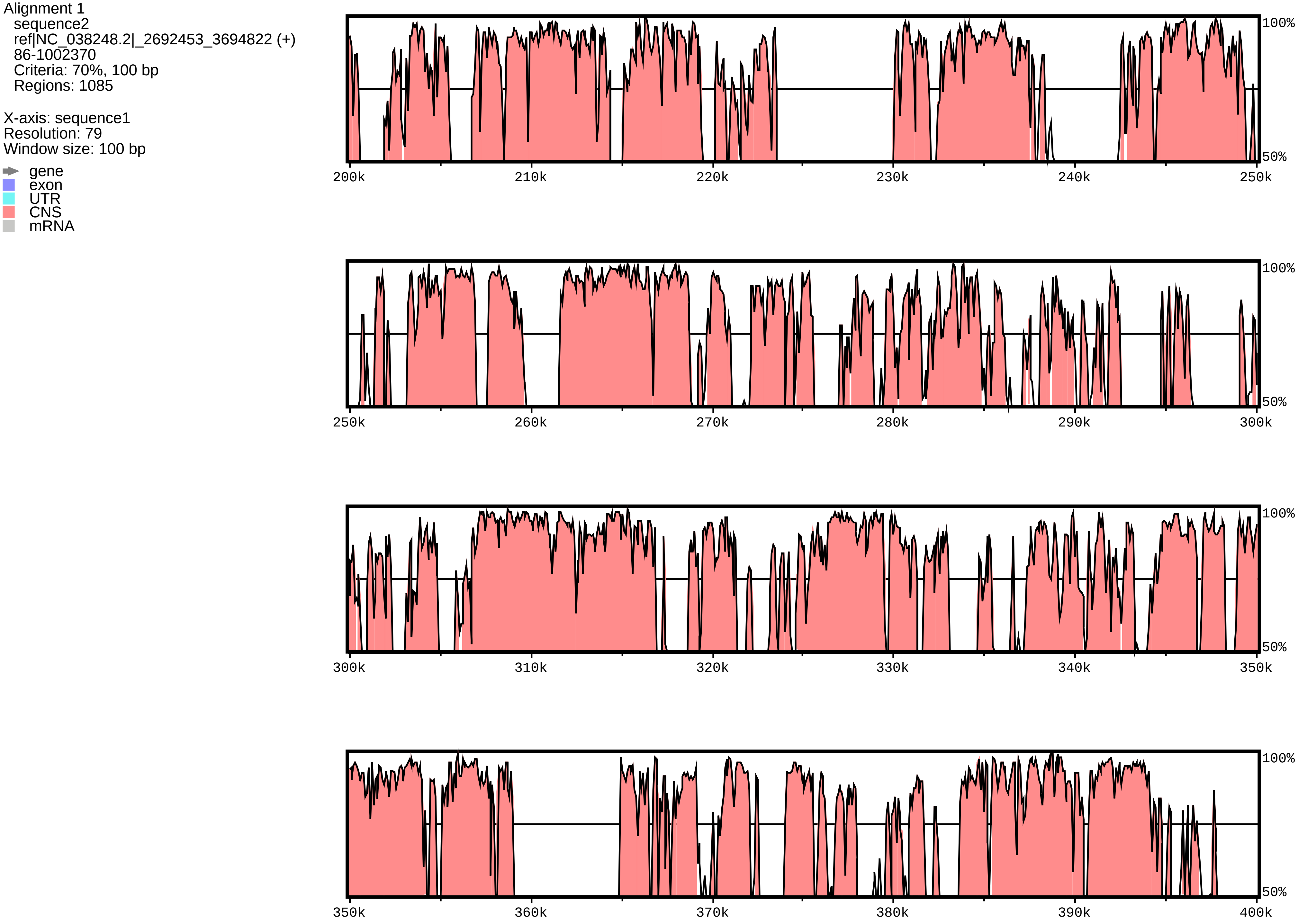

**sequence1 ref|NC_038247.2|_8469116_9635416:1-1166301**

Alignment 1
sequence2
ref|NC_038248.2|_2692453_3694822 (+)
86-1002370
Criteria: 70%, 100 bp
Regions: 1085

X-axis: sequence1
Resolution: 79
Window size: 100 bp

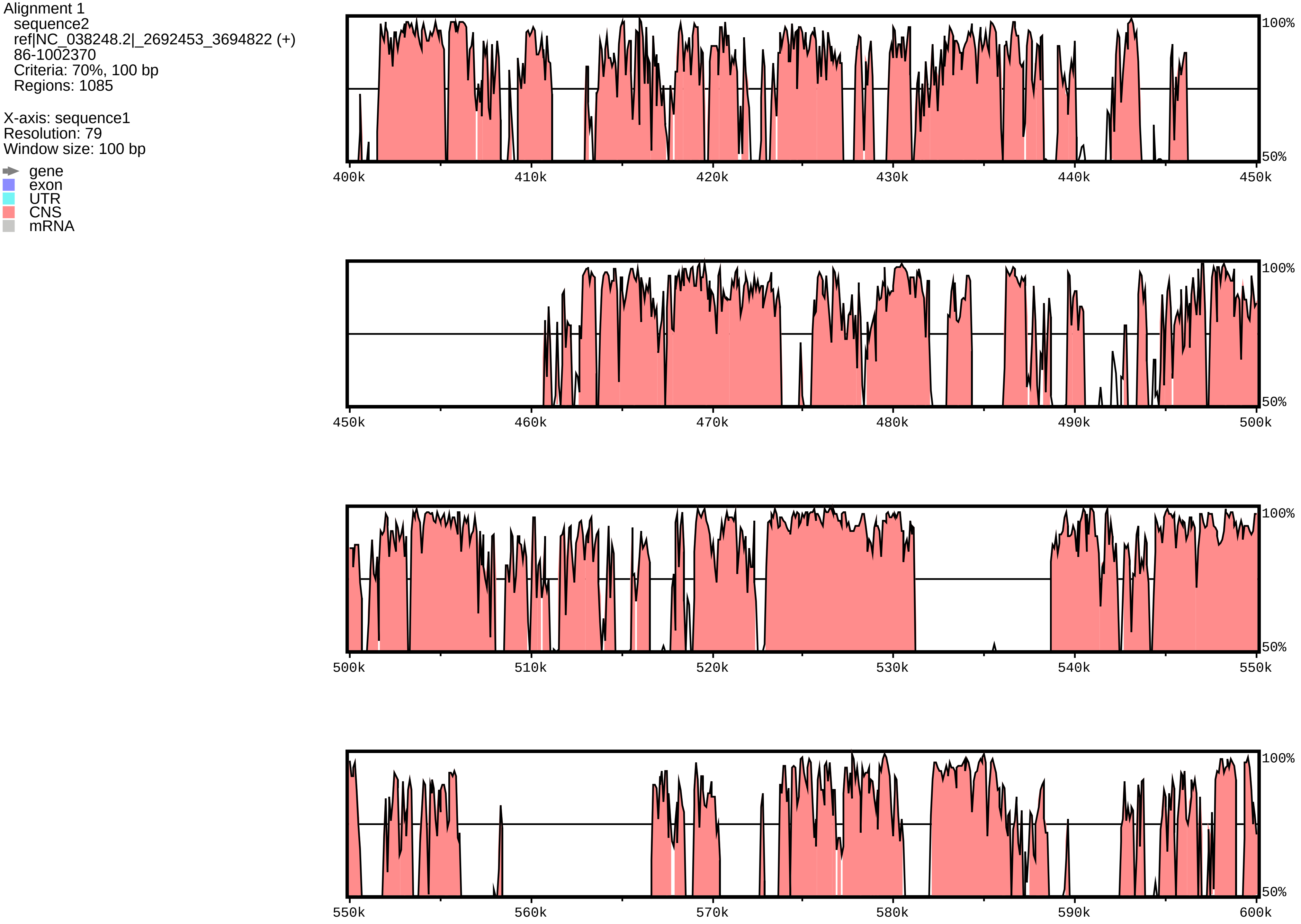

**sequence1 ref|NC_038247.2|_8469116_9635416:1-1166301**

Alignment 1
sequence2
ref|NC_038248.2|_2692453_3694822 (+)
86-1002370
Criteria: 70%, 100 bp
Regions: 1085

X-axis: sequence1
Resolution: 79
Window size: 100 bp

gene
exon
UTR
CNS
mRNA

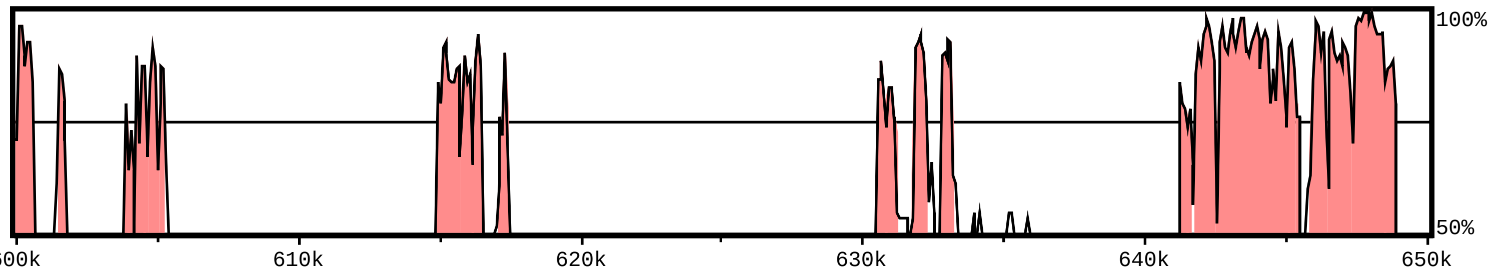


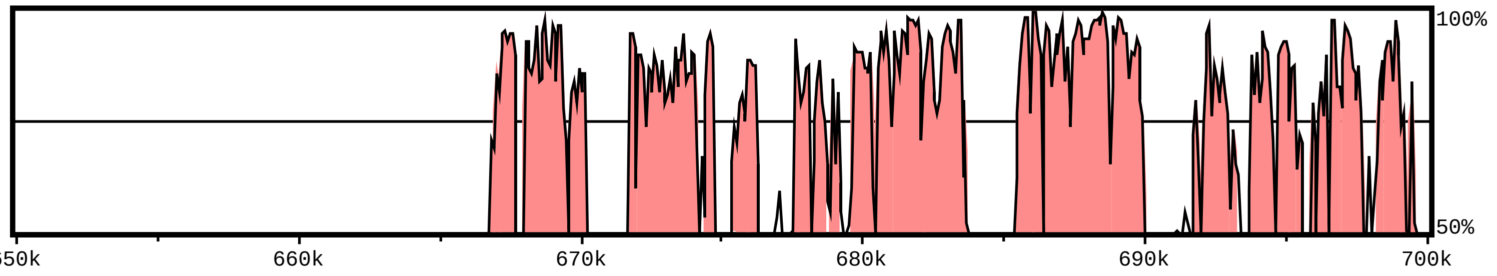


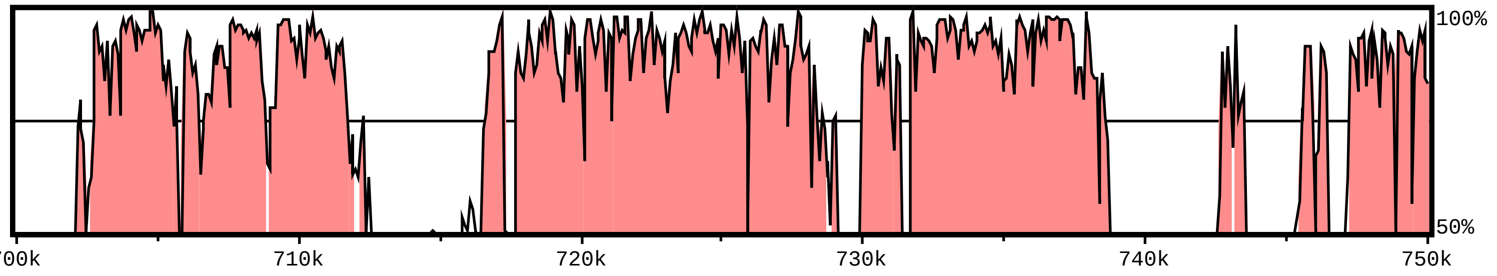


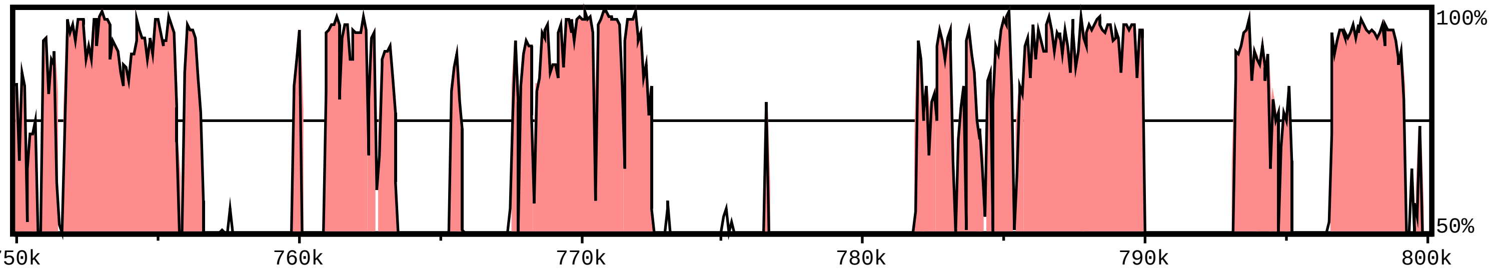

**sequence1 ref|NC_038247.2|_8469116_9635416:1-1166301**

Alignment 1
sequence2
ref|NC_038248.2|_2692453_3694822 (+)
86-1002370
Criteria: 70%, 100 bp
Regions: 1085

X-axis: sequence1
Resolution: 79
Window size: 100 bp

gene
exon
UTR
CNS
mRNA

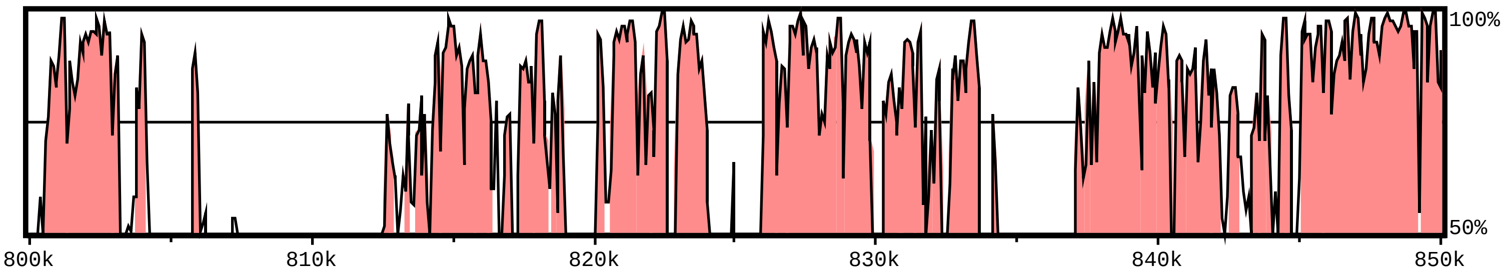


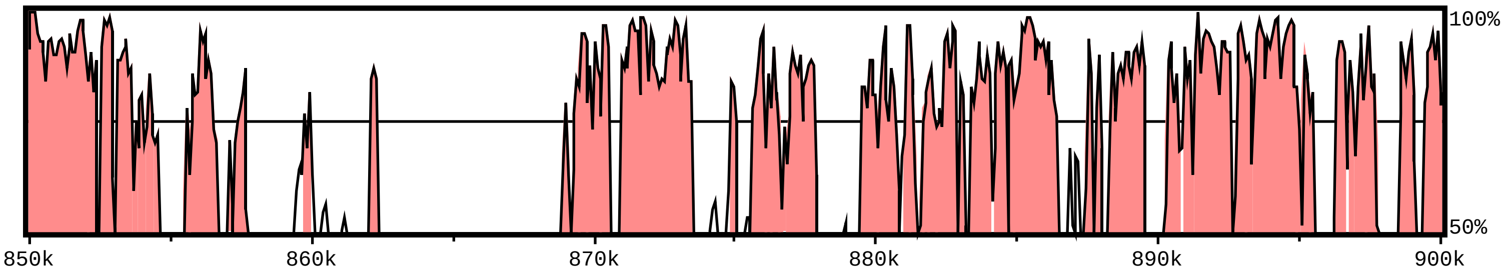


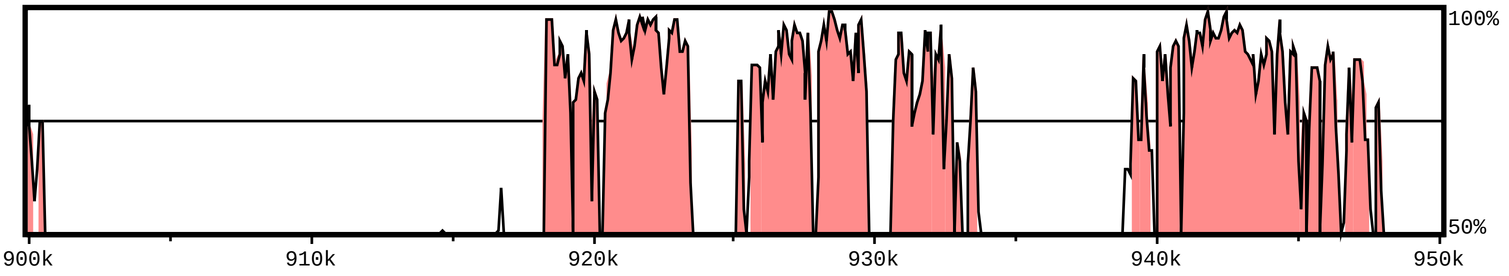


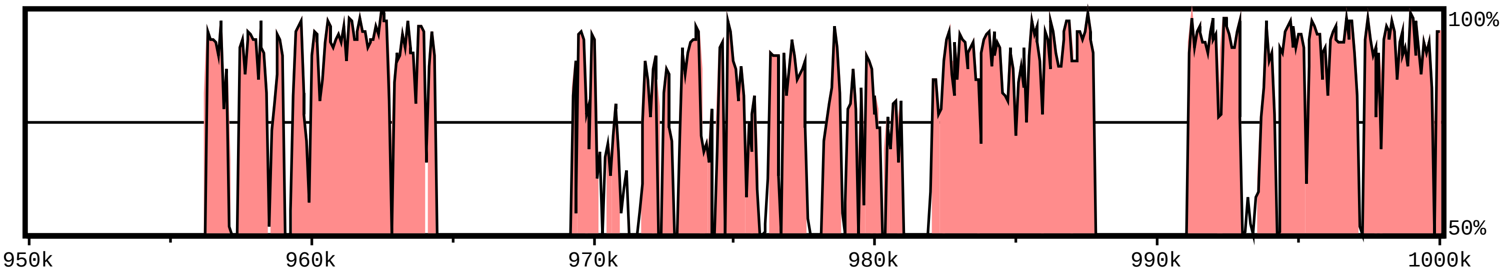

**sequence1 ref|NC_038247.2|_8469116_9635416:1-1166301**

Alignment 1
sequence2
ref|NC_038248.2|_2692453_3694822 (+)
86-1002370
Criteria: 70%, 100 bp
Regions: 1085

X-axis: sequence1
Resolution: 79
Window size: 100 bp

gene
exon
UTR
CNS
mRNA

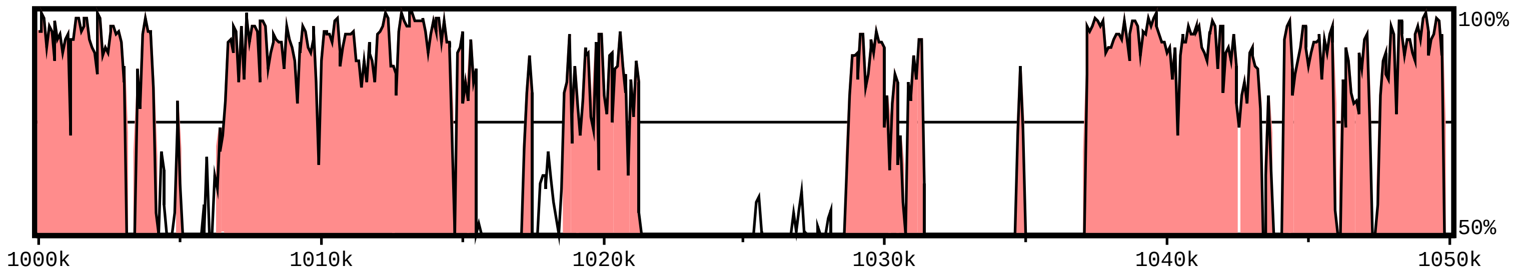


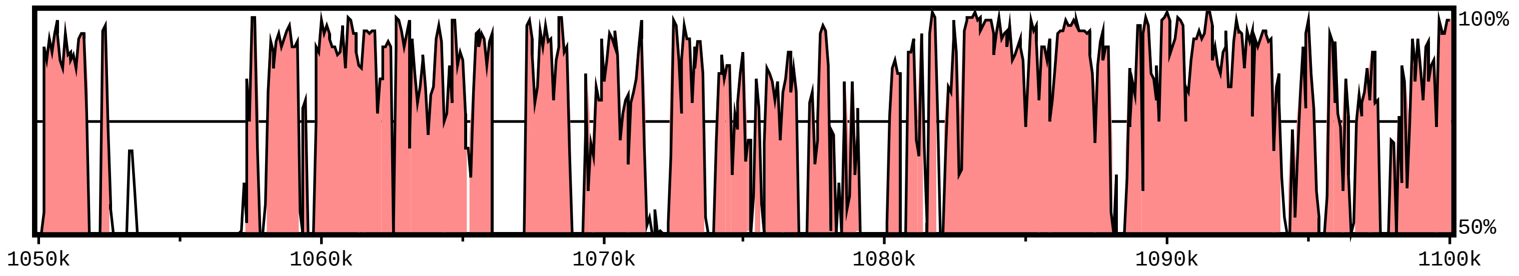


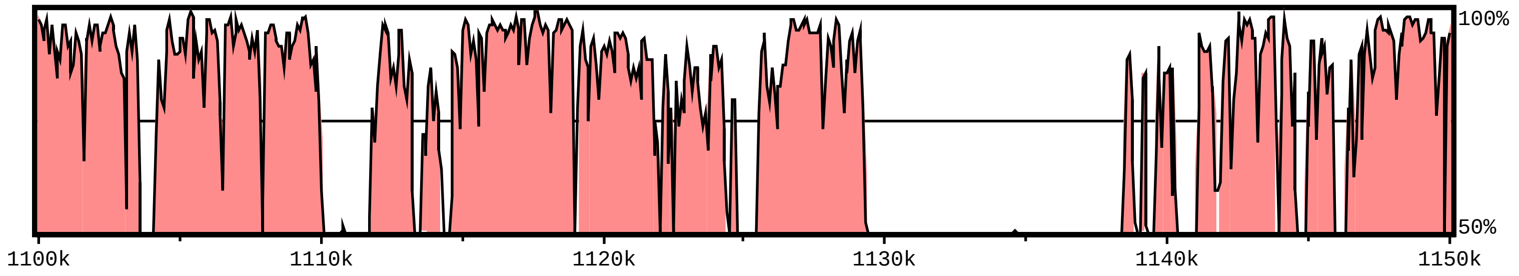


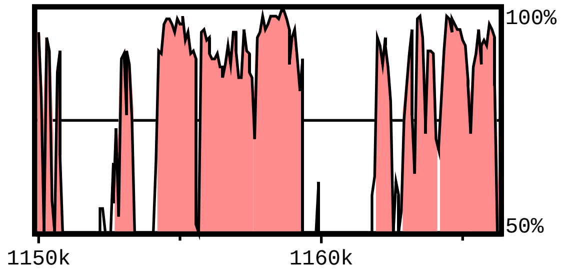
