## Supplementary Fig. S for "Comparative analysis of *Glycine BBX* gene family reveals lineage-specific evolution and expansion": figure 6.pdf

Supplementary Fig. S6. Multiple sequence alignment of GmBBX and GsBBX proteins. (a) The alignment shows only the regions of indels in GmBBX and GsBBX proteins, viz., GmBBX14b:GsBBX14b and GmBBX12g:GsBBX12g. (b) and (c) shows the BBX1 and BBX2 domain, respectively. Protein sequences are named to the left.

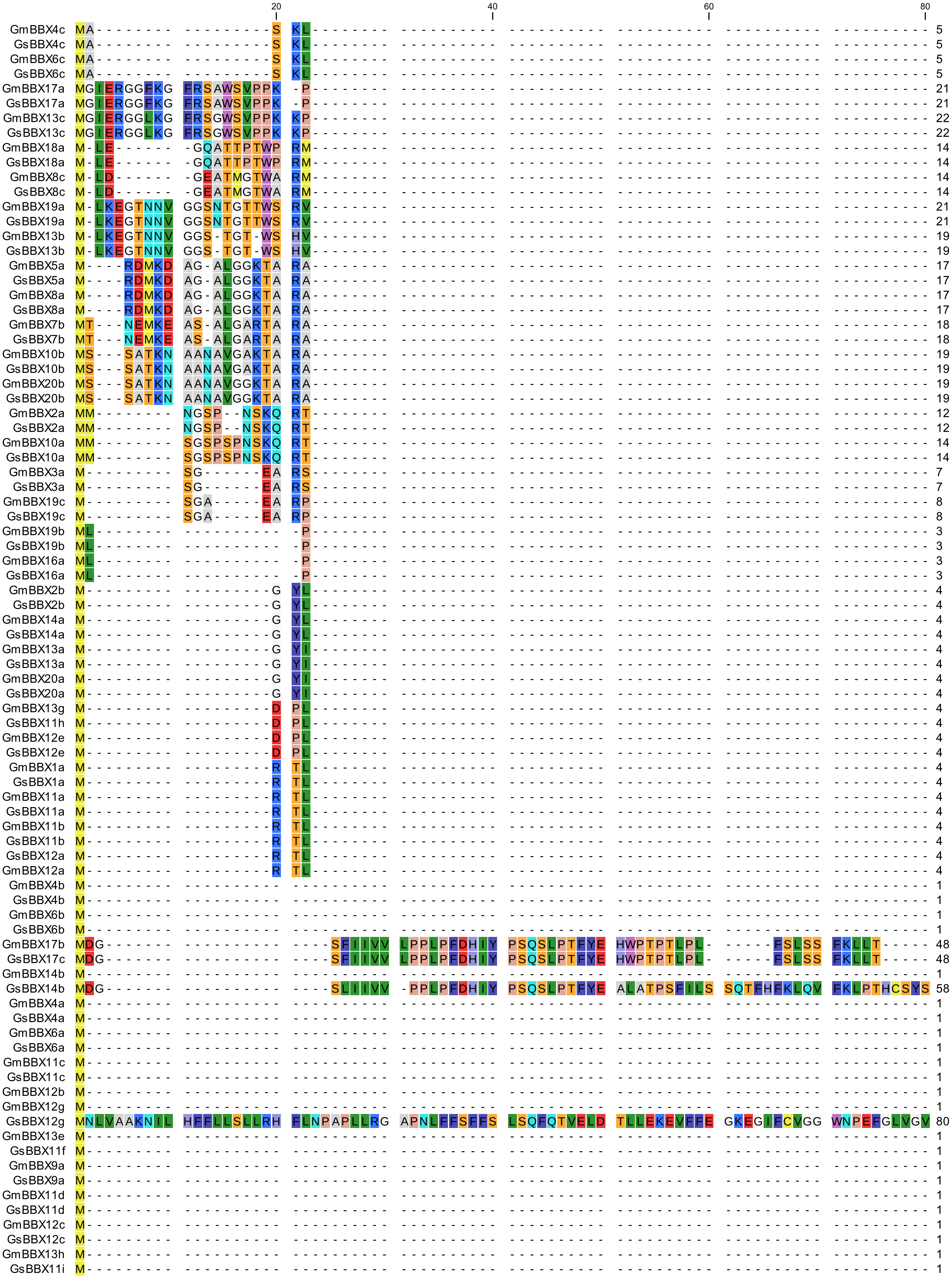

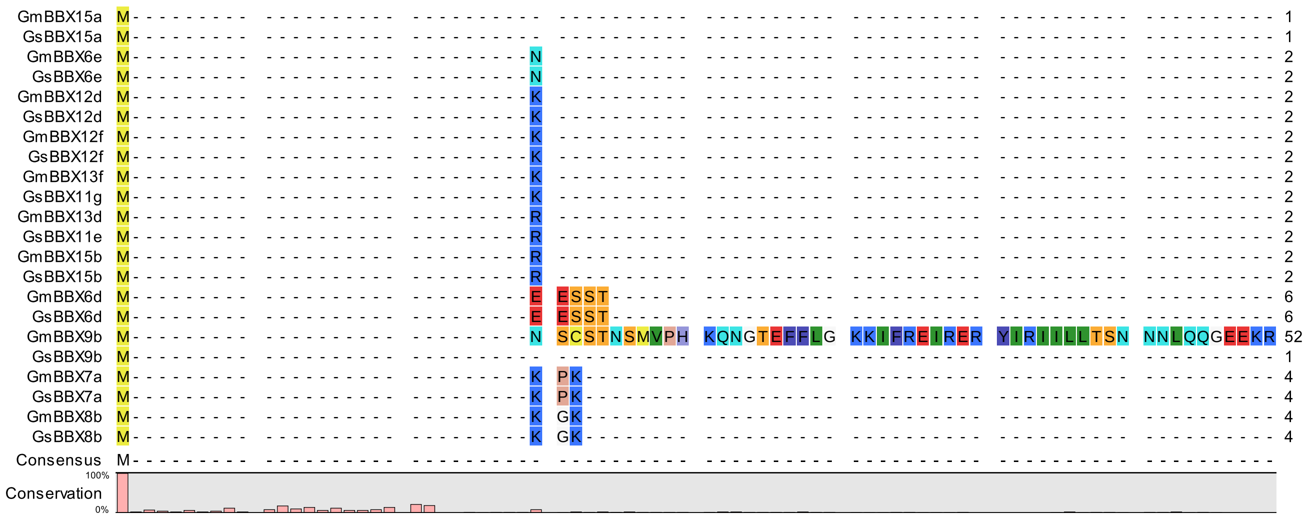
GmBBX15a
GsBBX15a
GmBBX6e
GsBBX6e
GmBBX12d
GsBBX12d
GmBBX12f
GsBBX12f
GmBBX13f
GsBBX11g
GmBBX13d
GsBBX11e
GmBBX15b
GsBBX15b
GmBBX6d
GsBBX6d
GmBBX9b
GsBBX9b
GmBBX7a
GsBBX7a
GmBBX8b
GsBBX8b
Consensus
Conservation
100%
0%

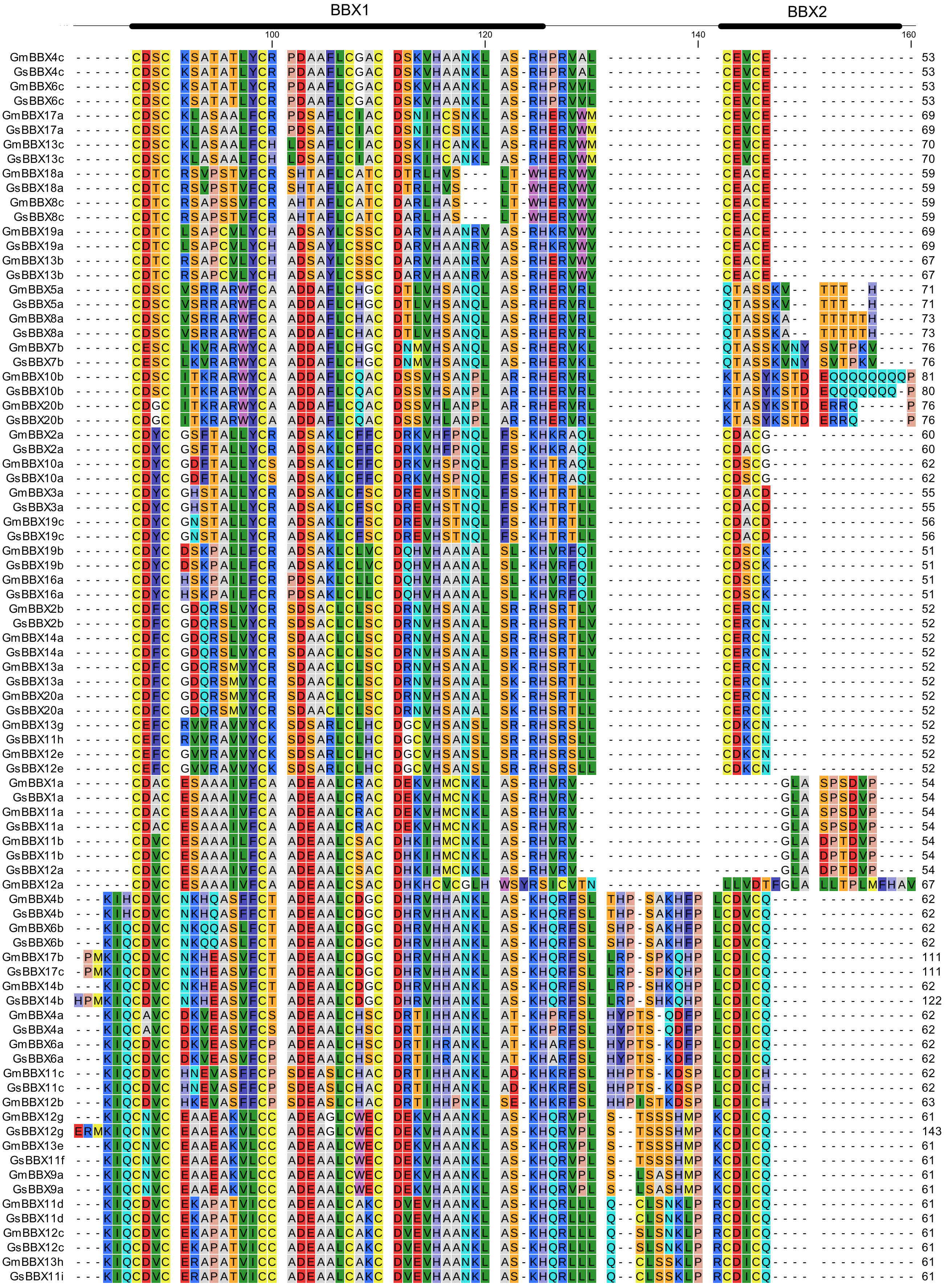
BBX1
BBX2
100
120
140
160

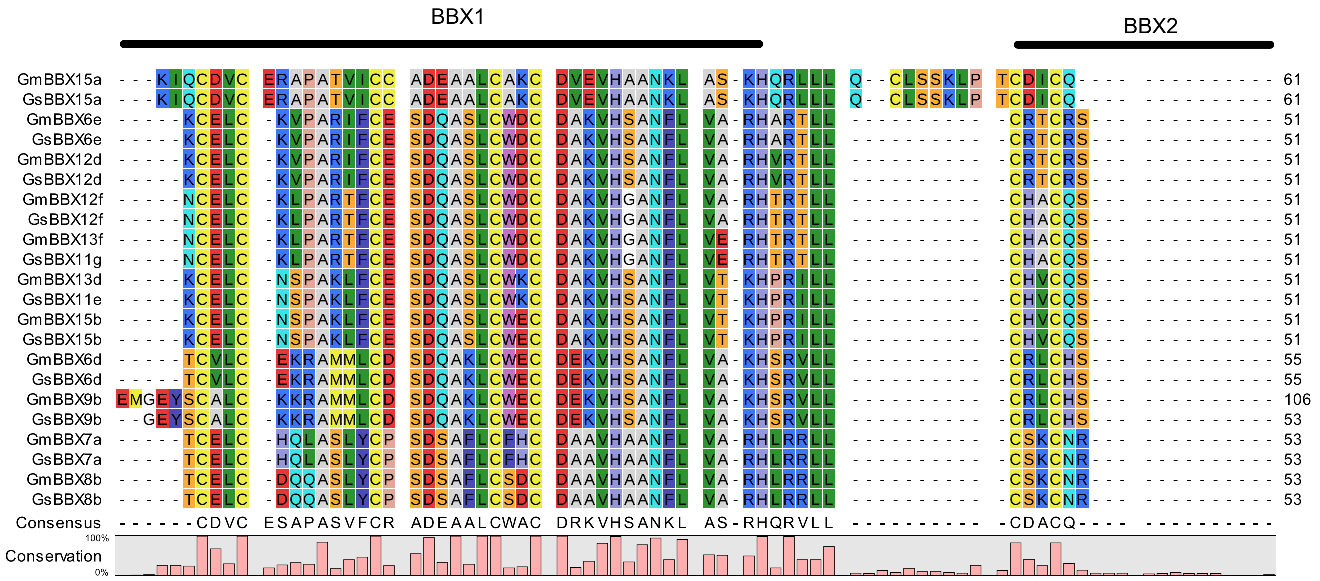
BBX1
BBX2
Consensus
Conservation
100%
0%

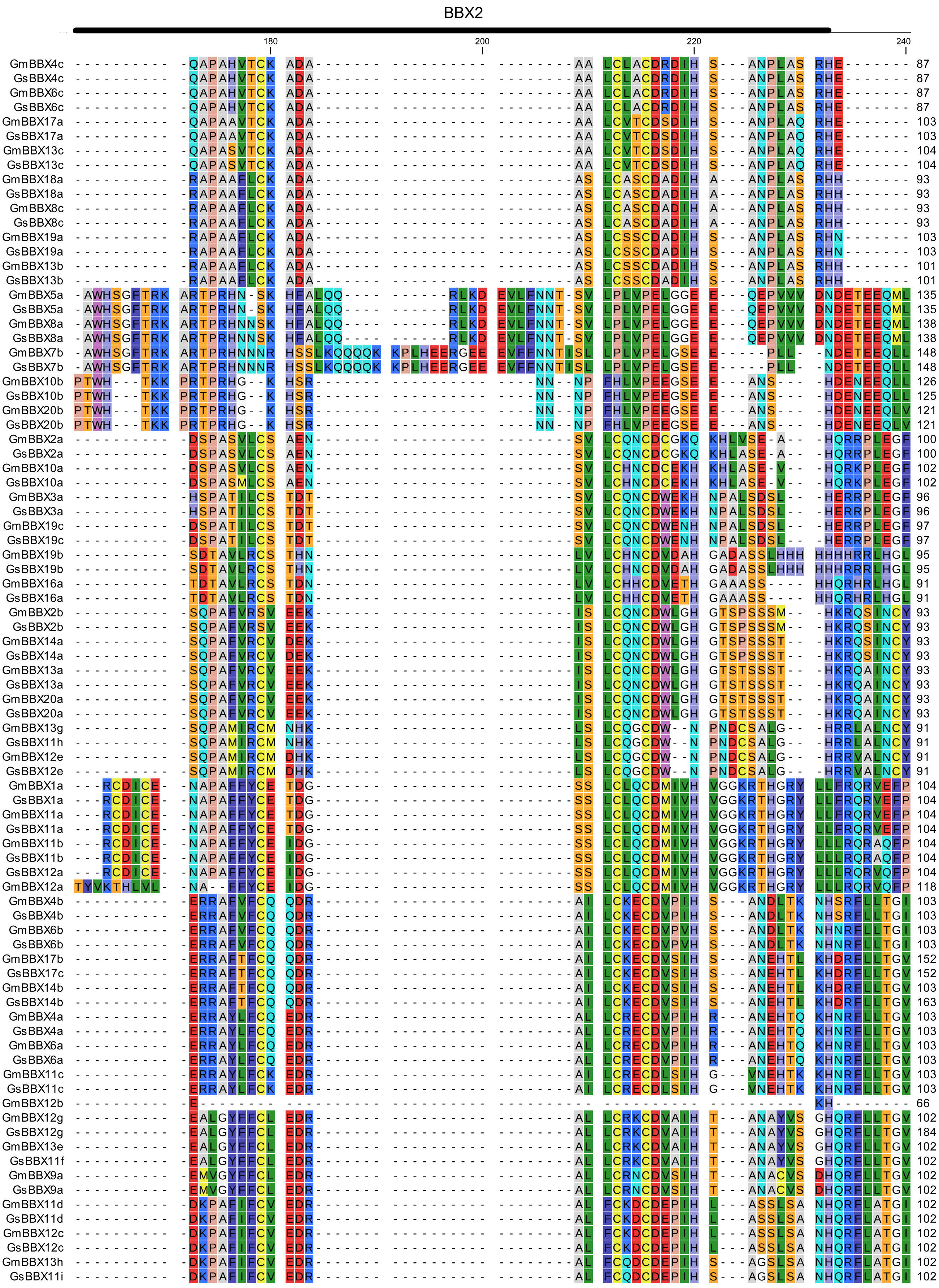
BBX2
180
200
220
240

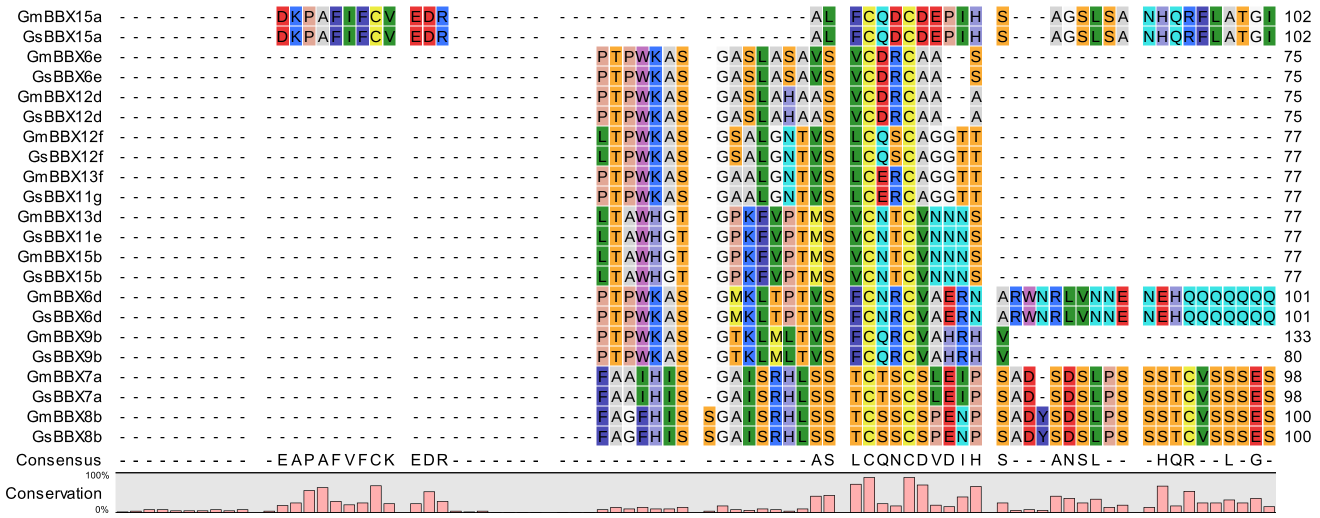
GmBBX15a ---------- -DKPAFIFCV EDR------- ---------- --------AL FCQDCDEPIH S---AGSLSA NHQRFLATGI 102
GsBBX15a ---------- -DKPAFIFCV EDR------- ---------- --------AL FCQDCDEPIH S---AGSLSA NHQRFLATGI 102
GmBBX6e ---------- ---------- ---------- ---PTPWKAS -GASLASAVS VCDRCAA--S ---------- ---------- 75
GsBBX6e ---------- ---------- ---------- ---PTPWKAS -GASLASAVS VCDRCAA--S ---------- ---------- 75
GmBBX12d ---------- ---------- ---------- ---PTPWKAS -GASLAHAAS VCDRCAA--A ---------- ---------- 75
GsBBX12d ---------- ---------- ---------- ---PTPWKAS -GASLAHAAS VCDRCAA--A ---------- ---------- 75
GmBBX12f ---------- ---------- ---------- ---LTPWKAS -GSALGNTVS LCQSCAGGTT ---------- ---------- 77
GsBBX12f ---------- ---------- ---------- ---LTPWKAS -GSALGNTVS LCQSCAGGTT ---------- ---------- 77
GmBBX13f ---------- ---------- ---------- ---PTPWKAS -GAALGNTVS LCERCAGGTT ---------- ---------- 77
GsBBX11g ---------- ---------- ---------- ---PTPWKAS -GAALGNTVS LCERCAGGTT ---------- ---------- 77
GmBBX13d ---------- ---------- ---------- ---LTAWHGT -GPKFVPTMS VCNTCVNNNS ---------- ---------- 77
GsBBX11e ---------- ---------- ---------- ---LTAWHGT -GPKFVPTMS VCNTCVNNNS ---------- ---------- 77
GmBBX15b ---------- ---------- ---------- ---LTAWHGT -GPKFVPTMS VCNTCVNNNS ---------- ---------- 77
GsBBX15b ---------- ---------- ---------- ---LTAWHGT -GPKFVPTMS VCNTCVNNNS ---------- ---------- 77
GmBBX6d ---------- ---------- ---------- ---PTPWKAS -GMKLTPTVS FCNRCVAERN ARWNRLVNNE NEHQQQQQQQ 101
GsBBX6d ---------- ---------- ---------- ---PTPWKAS -GMKLTPTVS FCNRCVAERN ARWNRLVNNE NEHQQQQQQQ 101
GmBBX9b ---------- ---------- ---------- ---PTPWKAS -GTKLMLTVS FCQRCVAHRH V--------- ---------- 133
GsBBX9b ---------- ---------- ---------- ---PTPWKAS -GTKLMLTVS FCQRCVAHRH V--------- ---------- 80
GmBBX7a ---------- ---------- ---------- ---FAAIHIS -GAISRHLSS TCTSCSLEIP SAD-SDSLPS SSTCVSSSES 98
GsBBX7a ---------- ---------- ---------- ---FAAIHIS -GAISRHLSS TCTSCSLEIP SAD-SDSLPS SSTCVSSSES 98
GmBBX8b ---------- ---------- ---------- ---FAGFHIS SGAISRHLSS TCSSCSPENP SADYSDSLPS SSTCVSSSES 100
GsBBX8b ---------- ---------- ---------- ---FAGFHIS SGAISRHLSS TCSSCSPENP SADYSDSLPS SSTCVSSSES 100
Consensus ---------- -EAPAFVFCK EDR------- ---------- --------AS LCQNCDVDIH S---ANSL-- -HQR--L-G-
Conservation
100%
0%

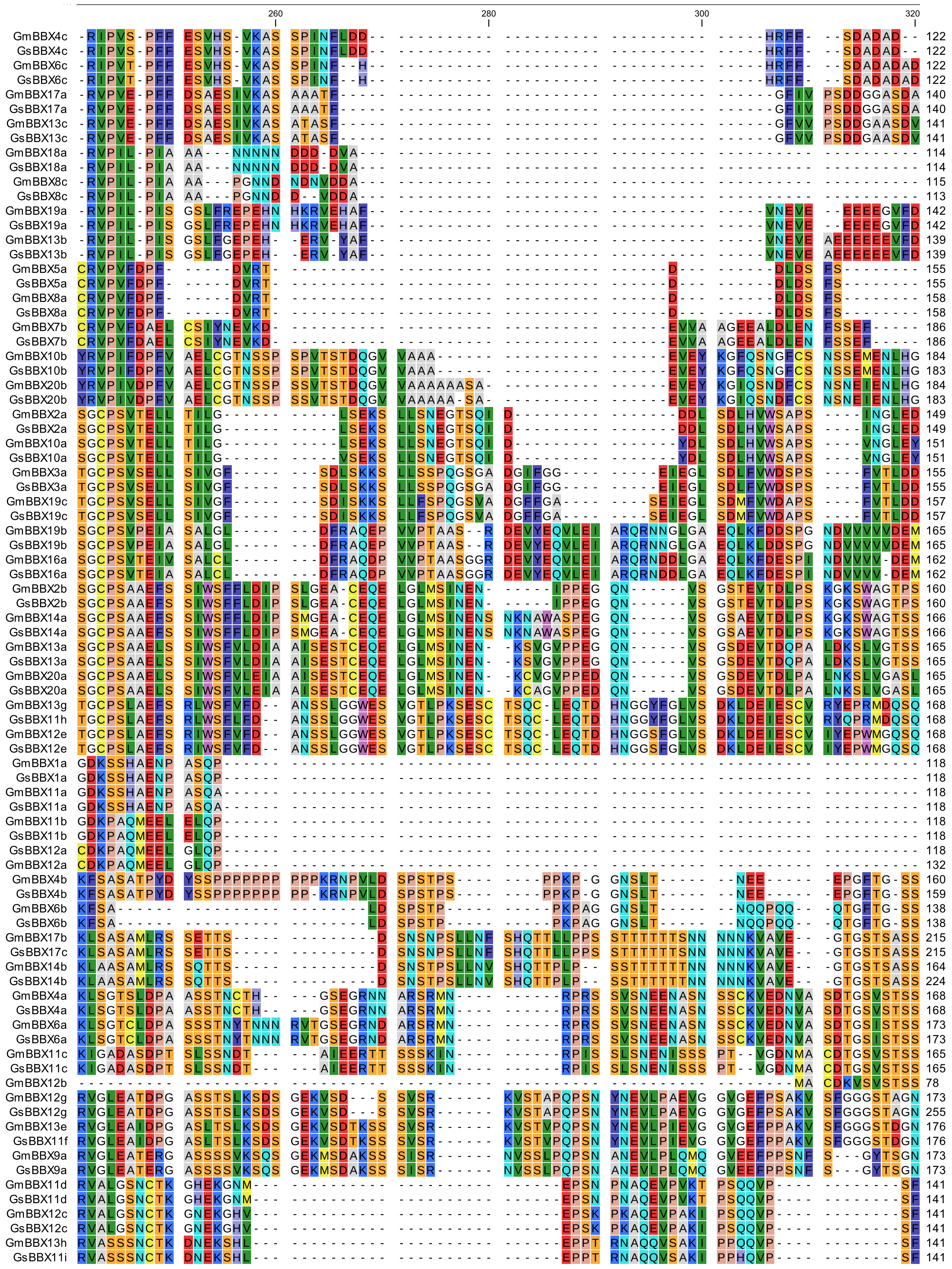

260
280
300
320
GmBBX4c 122
GsBBX4c 122
GmBBX6c 122
GsBBX6c 122
GmBBX17a 140
GsBBX17a 140
GmBBX13c 141
GsBBX13c 141
GmBBX18a 114
GsBBX18a 114
GmBBX8c 115
GsBBX8c 113
GmBBX19a 142
GsBBX19a 142
GmBBX13b 139
GsBBX13b 139
GmBBX5a 155
GsBBX5a 155
GmBBX8a 158
GsBBX8a 158
GmBBX7b 186
GsBBX7b 186
GmBBX10b 184
GsBBX10b 183
GmBBX20b 184
GsBBX20b 183
GmBBX2a 149
GsBBX2a 149
GmBBX10a 151
GsBBX10a 151
GmBBX3a 155
GsBBX3a 155
GmBBX19c 157
GsBBX19c 157
GmBBX19b 165
GsBBX19b 165
GmBBX16a 162
GsBBX16a 162
GmBBX2b 160
GsBBX2b 160
GmBBX14a 166
GsBBX14a 166
GmBBX13a 165
GsBBX13a 165
GmBBX20a 165
GsBBX20a 165
GmBBX13g 168
GsBBX11h 168
GmBBX12e 168
GsBBX12e 168
GmBBX1a 118
GsBBX1a 118
GmBBX11a 118
GsBBX11a 118
GmBBX11b 118
GsBBX11b 118
GsBBX12a 118
GmBBX12a 132
GmBBX4b 160
GsBBX4b 159
GmBBX6b 138
GsBBX6b 138
GmBBX17b 215
GsBBX17c 215
GmBBX14b 164
GsBBX14b 224
GmBBX4a 168
GsBBX4a 168
GmBBX6a 173
GsBBX6a 173
GmBBX11c 165
GsBBX11c 165
GmBBX12b 78
GmBBX12g 173
GsBBX12g 255
GmBBX13e 176
GsBBX11f 176
GmBBX9a 173
GsBBX9a 173
GmBBX11d 141
GsBBX11d 141
GmBBX12c 141
GsBBX12c 141
GmBBX13h 141
GsBBX11i 141

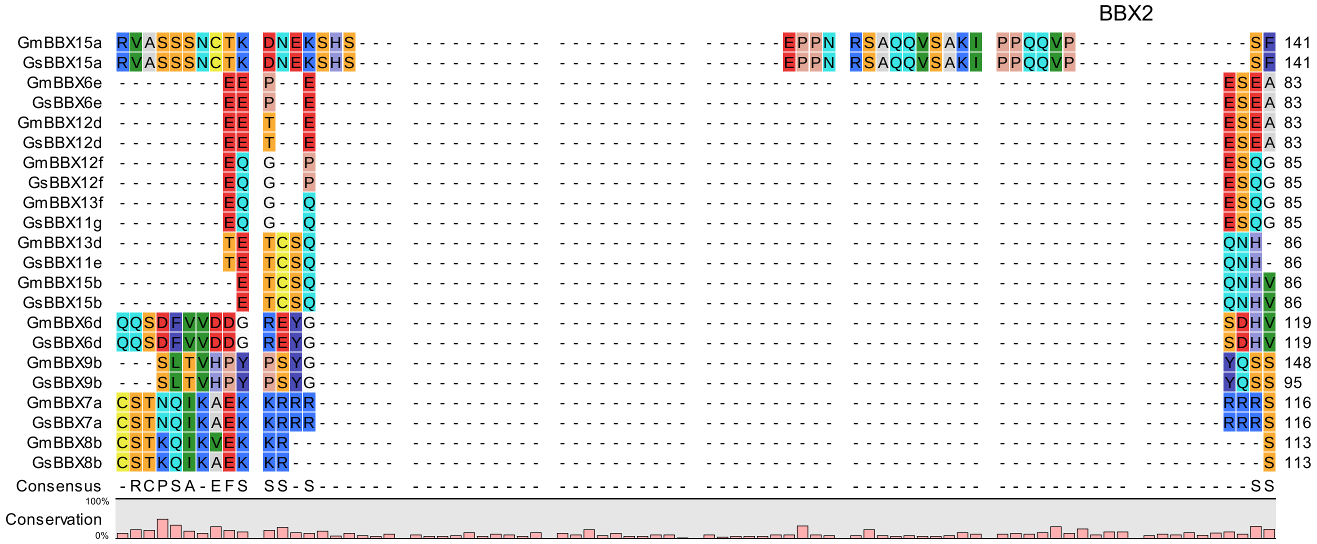
BBX2
GmBBX15a RVASSSNCTK DNEKSHS EPPN RSAQQVSAKI PPQQVP SF 141
GsBBX15a RVASSSNCTK DNEKSHS EPPN RSAQQVSAKI PPQQVP SF 141
GmBBX6e EE P E ESEA 83
GsBBX6e EE P E ESEA 83
GmBBX12d EE T E ESEA 83
GsBBX12d EE T E ESEA 83
GmBBX12f EQ G P ESQG 85
GsBBX12f EQ G P ESQG 85
GmBBX13f EQ G Q ESQG 85
GsBBX11g EQ G Q ESQG 85
GmBBX13d TE TCSQ QNH 86
GsBBX11e TE TCSQ QNH 86
GmBBX15b E TCSQ QNHV 86
GsBBX15b E TCSQ QNHV 86
GmBBX6d QQSDFVVDDG REYG SDHV 119
GsBBX6d QQSDFVVDDG REYG SDHV 119
GmBBX9b SLTVHPY PSYG YQSS 148
GsBBX9b SLTVHPY PSYG YQSS 95
GmBBX7a CSTNQIKAEK KRRR RRRS 116
GsBBX7a CSTNQIKAEK KRRR RRRS 116
GmBBX8b CSTKQIKVEK KR S 113
GsBBX8b CSTKQIKAEK KR S 113
Consensus -RCPSA-EFS SS-S SS
Conservation
100%
0%

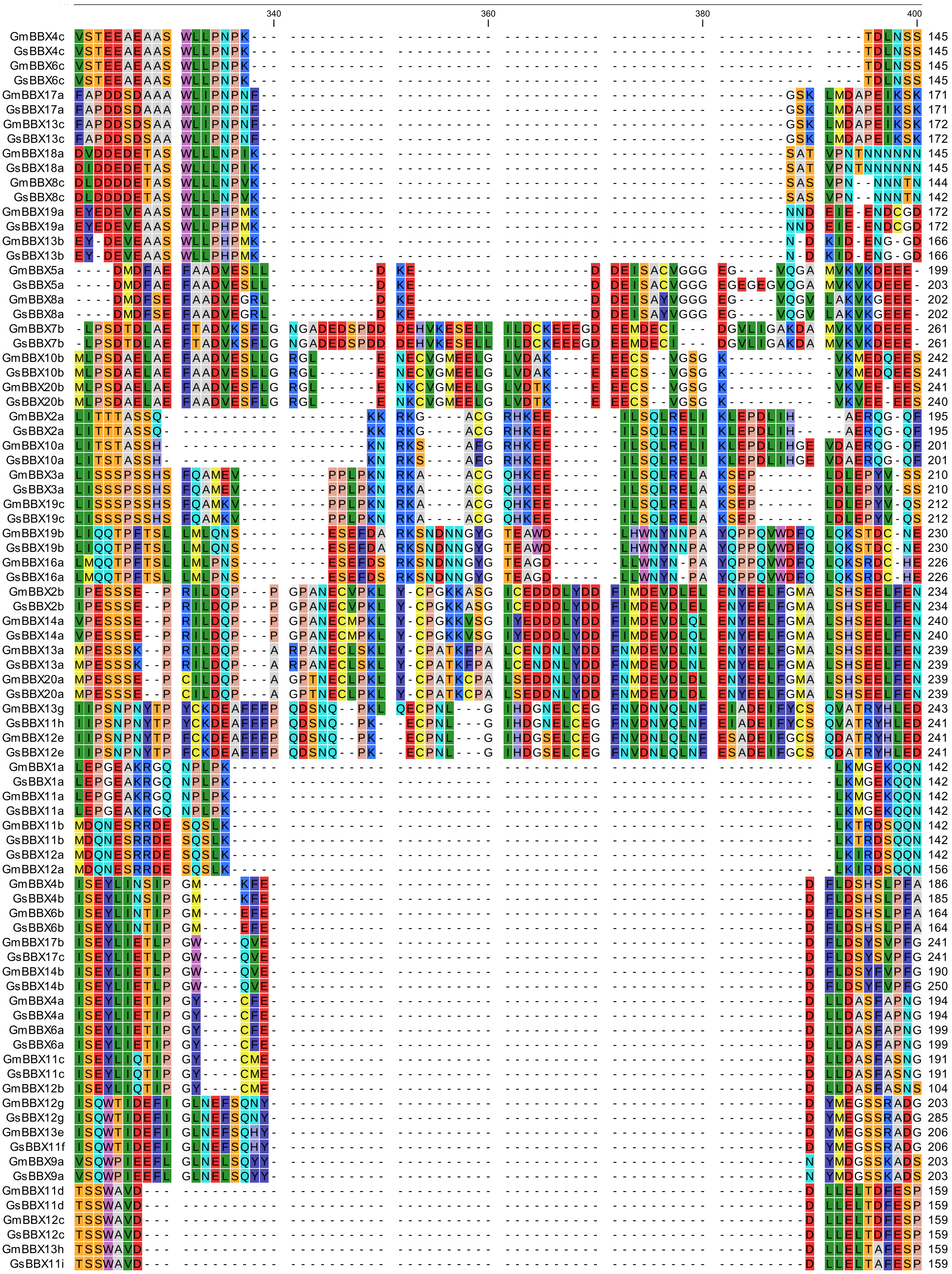
340
360
380
400
GmBBX4c 145
GsBBX4c 145
GmBBX6c 145
GsBBX6c 145
GmBBX17a 171
GsBBX17a 171
GmBBX13c 172
GsBBX13c 172
GmBBX18a 145
GsBBX18a 145
GmBBX8c 144
GsBBX8c 142
GmBBX19a 172
GsBBX19a 172
GmBBX13b 166
GsBBX13b 166
GmBBX5a 199
GsBBX5a 203
GmBBX8a 202
GsBBX8a 202
GmBBX7b 261
GsBBX7b 261
GmBBX10b 242
GsBBX10b 241
GmBBX20b 241
GsBBX20b 240
GmBBX2a 195
GsBBX2a 195
GmBBX10a 201
GsBBX10a 201
GmBBX3a 210
GsBBX3a 210
GmBBX19c 212
GsBBX19c 212
GmBBX19b 230
GsBBX19b 230
GmBBX16a 226
GsBBX16a 226
GmBBX2b 234
GsBBX2b 234
GmBBX14a 240
GsBBX14a 240
GmBBX13a 239
GsBBX13a 239
GmBBX20a 239
GsBBX20a 239
GmBBX13g 243
GsBBX11h 241
GmBBX12e 241
GsBBX12e 241
GmBBX1a 142
GsBBX1a 142
GmBBX11a 142
GsBBX11a 142
GmBBX11b 142
GsBBX11b 142
GsBBX12a 142
GmBBX12a 156
GmBBX4b 186
GsBBX4b 185
GmBBX6b 164
GsBBX6b 164
GmBBX17b 241
GsBBX17c 241
GmBBX14b 190
GsBBX14b 250
GmBBX4a 194
GsBBX4a 194
GmBBX6a 199
GsBBX6a 199
GmBBX11c 191
GsBBX11c 191
GmBBX12b 104
GmBBX12g 203
GsBBX12g 285
GmBBX13e 206
GsBBX11f 206
GmBBX9a 203
GsBBX9a 203
GmBBX11d 159
GsBBX11d 159
GmBBX12c 159
GsBBX12c 159
GmBBX13h 159
GsBBX11i 159

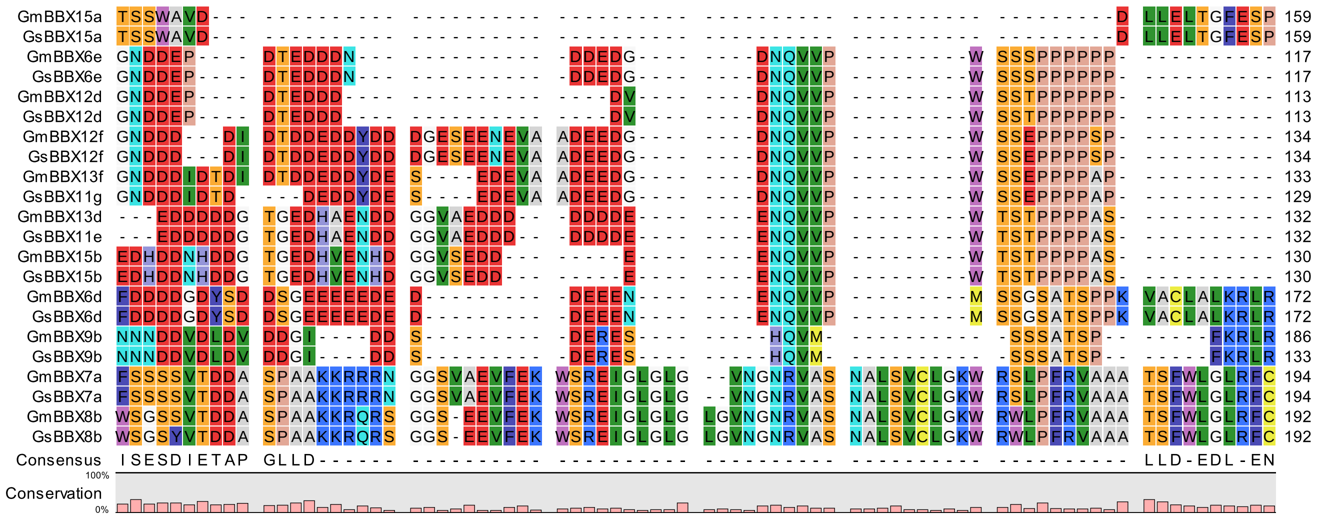
GmBBX15a 159
GsBBX15a 159
GmBBX6e 117
GsBBX6e 117
GmBBX12d 113
GsBBX12d 113
GmBBX12f 134
GsBBX12f 134
GmBBX13f 133
GsBBX11g 129
GmBBX13d 132
GsBBX11e 132
GmBBX15b 130
GsBBX15b 130
GmBBX6d 172
GsBBX6d 172
GmBBX9b 186
GsBBX9b 133
GmBBX7a 194
GsBBX7a 194
GmBBX8b 192
GsBBX8b 192
Consensus
Conservation
100%
0%

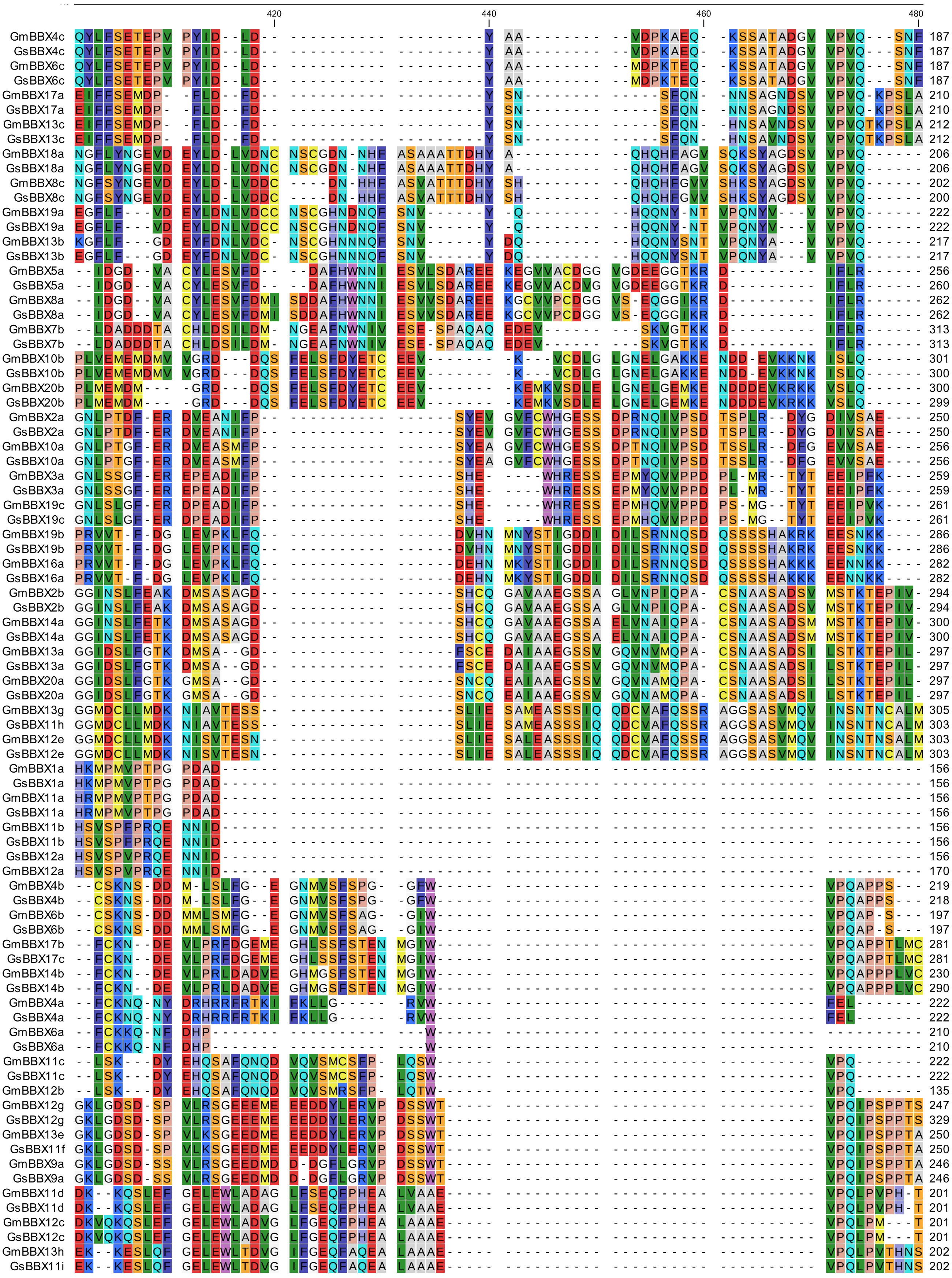

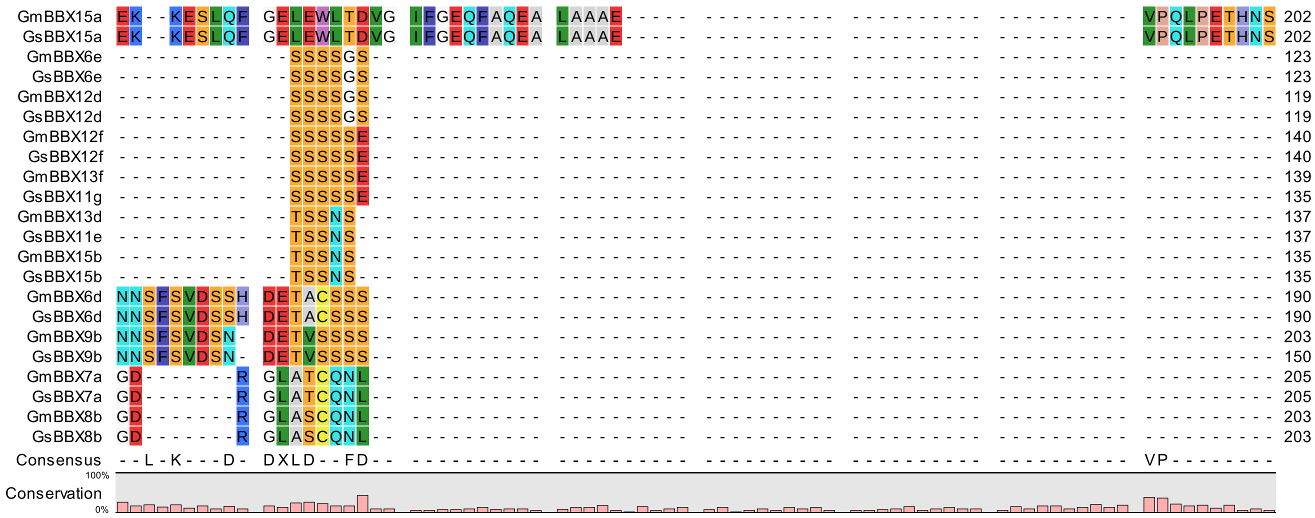

GmBBX15a EK--KESLQF GELEWLTDVG IFGEQFAQEA LAAAE----- ---------- ---------- ---------- VPQLPETHNS 202
GsBBX15a EK--KESLQF GELEWLTDVG IFGEQFAQEA LAAAE----- ---------- ---------- ---------- VPQLPETHNS 202
GmBBX6e ---------- --SSSSGS-- ---------- ---------- ---------- ---------- ---------- ---------- 123
GsBBX6e ---------- --SSSSGS-- ---------- ---------- ---------- ---------- ---------- ---------- 123
GmBBX12d ---------- --SSSSGS-- ---------- ---------- ---------- ---------- ---------- ---------- 119
GsBBX12d ---------- --SSSSGS-- ---------- ---------- ---------- ---------- ---------- ---------- 119
GmBBX12f ---------- --SSSSSE-- ---------- ---------- ---------- ---------- ---------- ---------- 140
GsBBX12f ---------- --SSSSSE-- ---------- ---------- ---------- ---------- ---------- ---------- 140
GmBBX13f ---------- --SSSSSE-- ---------- ---------- ---------- ---------- ---------- ---------- 139
GsBBX11g ---------- --SSSSSE-- ---------- ---------- ---------- ---------- ---------- ---------- 135
GmBBX13d ---------- --TSSNS--- ---------- ---------- ---------- ---------- ---------- ---------- 137
GsBBX11e ---------- --TSSNS--- ---------- ---------- ---------- ---------- ---------- ---------- 137
GmBBX15b ---------- --TSSNS--- ---------- ---------- ---------- ---------- ---------- ---------- 135
GsBBX15b ---------- --TSSNS--- ---------- ---------- ---------- ---------- ---------- ---------- 135
GmBBX6d NNSFSVDSSH DETACSSS-- ---------- ---------- ---------- ---------- ---------- ---------- 190
GsBBX6d NNSFSVDSSH DETACSSS-- ---------- ---------- ---------- ---------- ---------- ---------- 190
GmBBX9b NNSFSVDSN- DETVSSSS-- ---------- ---------- ---------- ---------- ---------- ---------- 203
GsBBX9b NNSFSVDSN- DETVSSSS-- ---------- ---------- ---------- ---------- ---------- ---------- 150
GmBBX7a GD-------R GLATCQNL-- ---------- ---------- ---------- ---------- ---------- ---------- 205
GsBBX7a GD-------R GLATCQNL-- ---------- ---------- ---------- ---------- ---------- ---------- 205
GmBBX8b GD-------R GLASCQNL-- ---------- ---------- ---------- ---------- ---------- ---------- 203
GsBBX8b GD-------R GLASCQNL-- ---------- ---------- ---------- ---------- ---------- ---------- 203
Consensus --L-K---D- DXLD--FD-- ---------- ---------- ---------- ---------- ---------- VP--------
100%
Conservation
0%

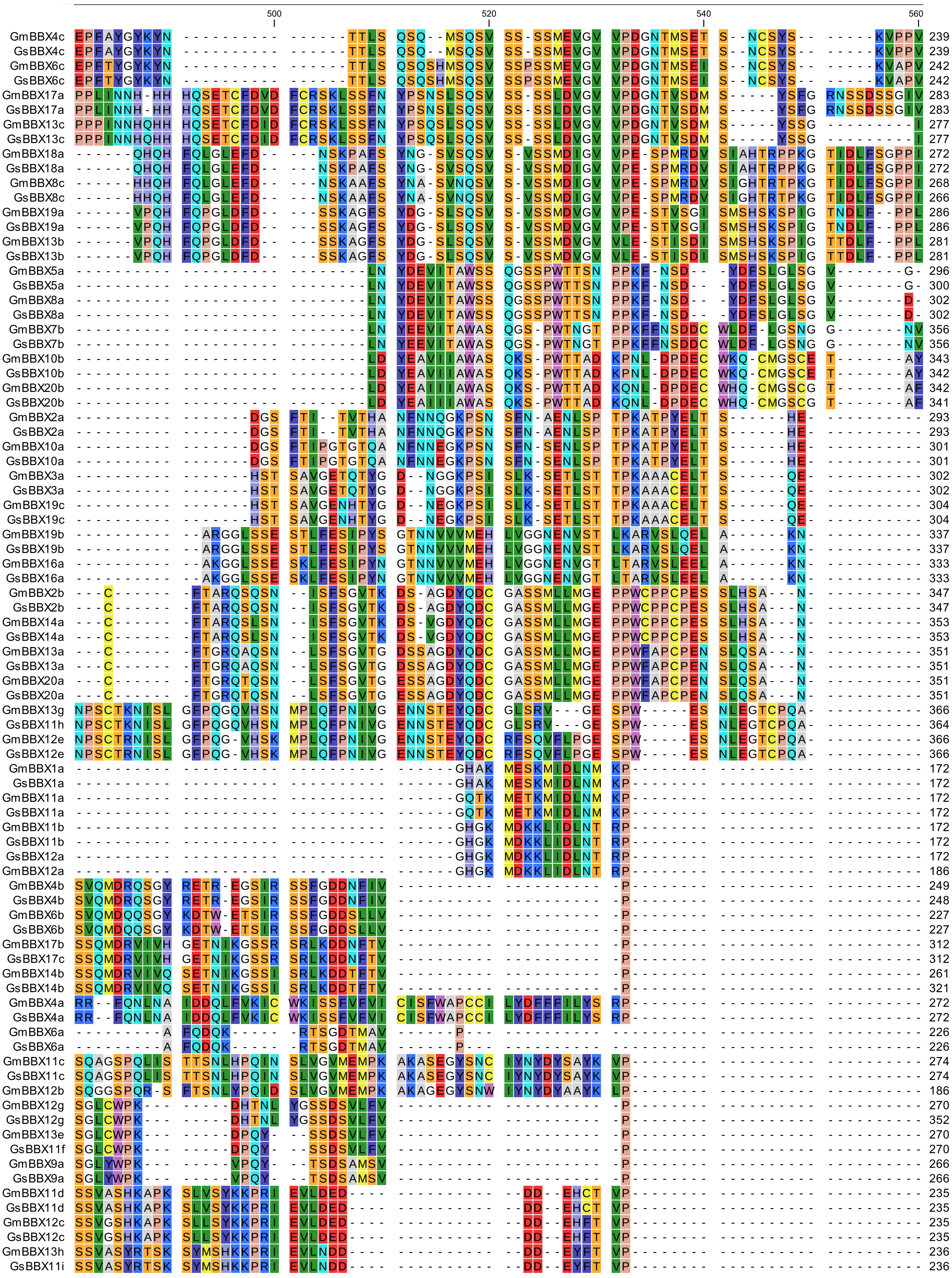

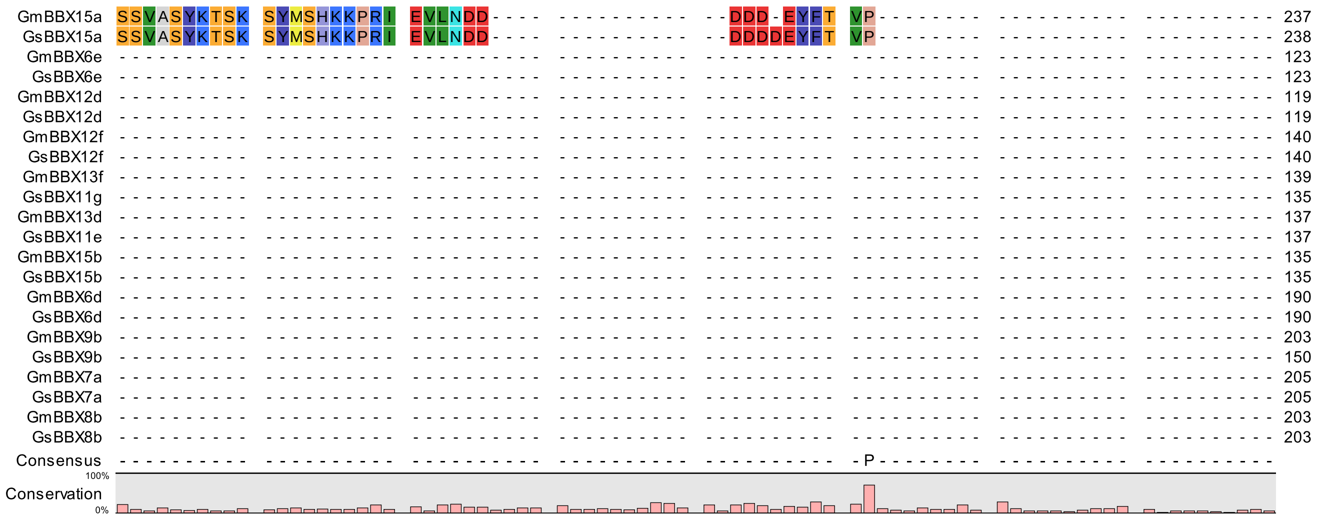

GmBBX15a SSVASYKTSK SYMSHKKPRI EVLNDD---- ---------- --DDD-EYFT VP-------- ---------- ---------- 237
GsBBX15a SSVASYKTSK SYMSHKKPRI EVLNDD---- ---------- --DDDDEYFT VP-------- ---------- ---------- 238
GmBBX6e 123
GsBBX6e 123
GmBBX12d 119
GsBBX12d 119
GmBBX12f 140
GsBBX12f 140
GmBBX13f 139
GsBBX11g 135
GmBBX13d 137
GsBBX11e 137
GmBBX15b 135
GsBBX15b 135
GmBBX6d 190
GsBBX6d 190
GmBBX9b 203
GsBBX9b 150
GmBBX7a 205
GsBBX7a 205
GmBBX8b 203
GsBBX8b 203
Consensus -P
Conservation
100%
0%

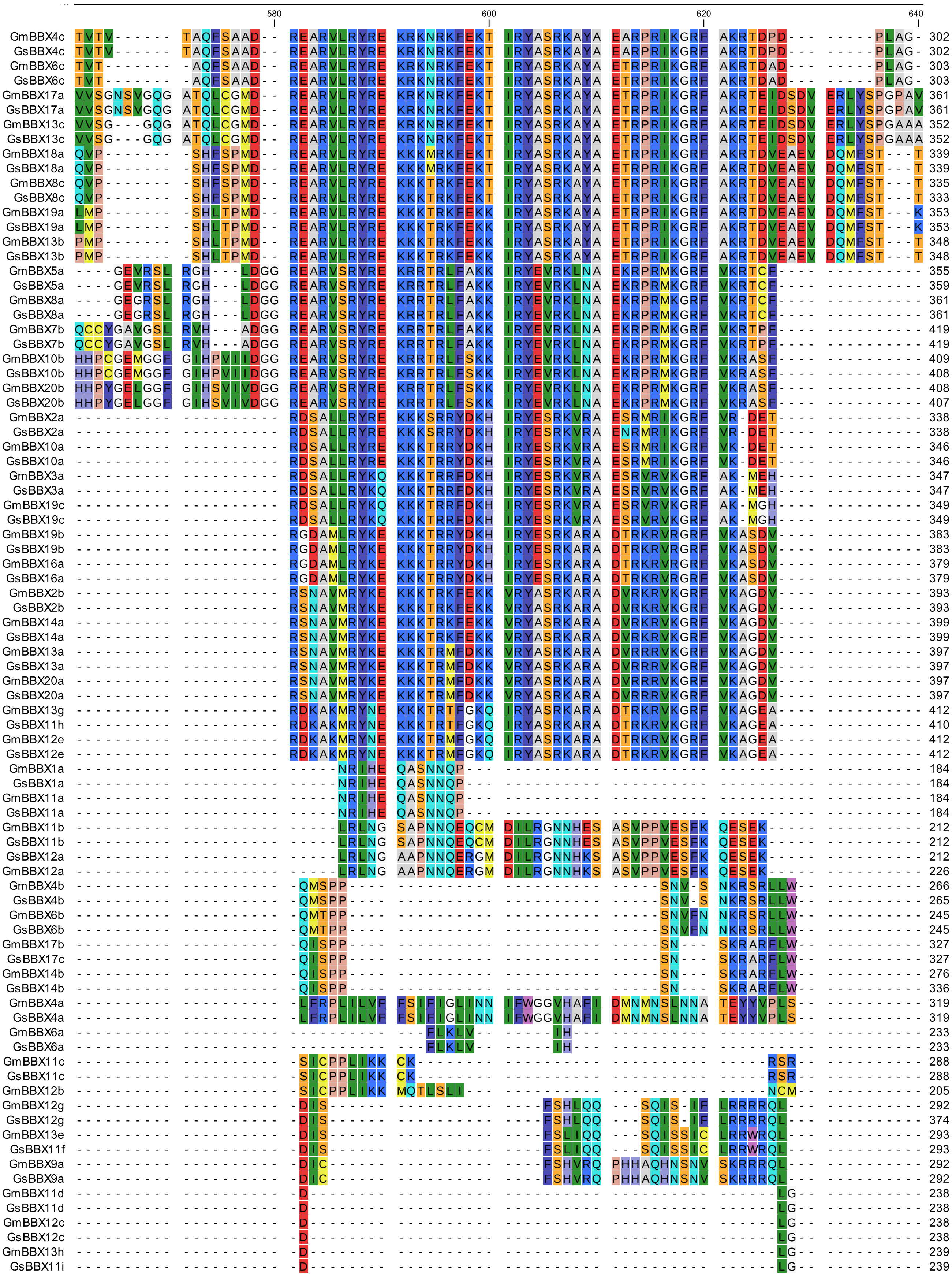
580
600
620
640

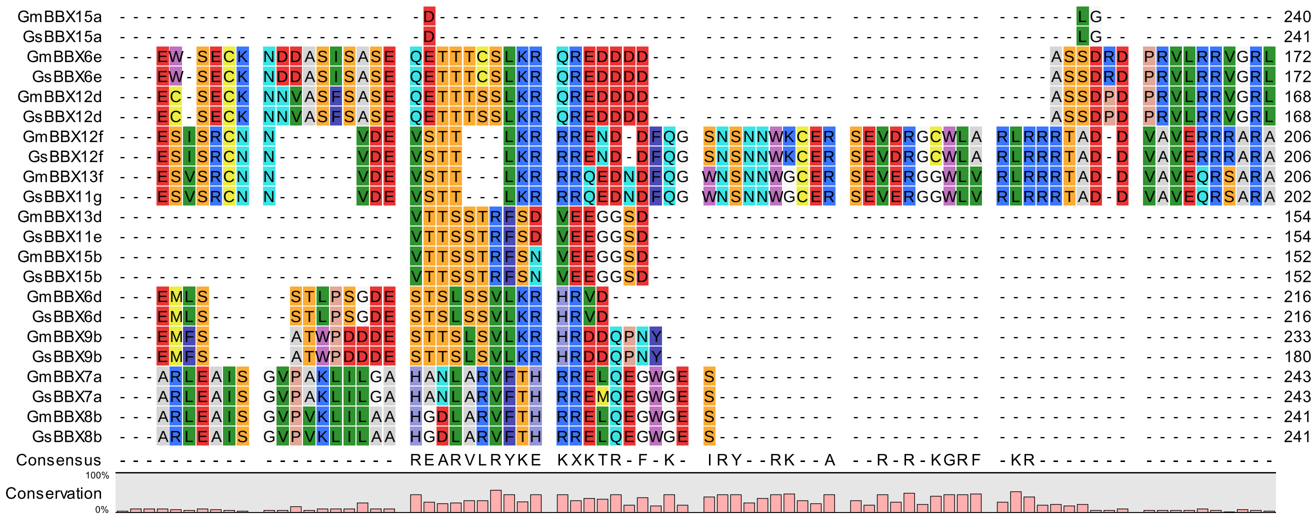
GmBBX15a 240
GsBBX15a 241
GmBBX6e 172
GsBBX6e 172
GmBBX12d 168
GsBBX12d 168
GmBBX12f 206
GsBBX12f 206
GmBBX13f 206
GsBBX11g 202
GmBBX13d 154
GsBBX11e 154
GmBBX15b 152
GsBBX15b 152
GmBBX6d 216
GsBBX6d 216
GmBBX9b 233
GsBBX9b 180
GmBBX7a 243
GsBBX7a 243
GmBBX8b 241
GsBBX8b 241
Consensus
REARVLRYKE KXKTR-F-K- IRY--RK--A --R-R-KGRF -KR
Conservation
100%
0%

```
GmBBX4c   ------YGVV PSC-- 309
GsBBX4c   ------YGVV PSC-- 309
GmBBX6c   ------YGVV PSC-- 310
GsBBX6c   ------YGVV PSC-- 310
GmBBX17a  LMLDTPYGVV PSF-- 374
GsBBX17a  LMLDTPYGVV PSF-- 374
GmBBX13c  LMLDTPYGVV PTF-- 365
GsBBX13c  LMLDTPYGVV PSF-- 365
GmBBX18a  LITEVGYGIV PSF-- 352
GsBBX18a  LITEVGYGIV PSF-- 352
GmBBX8c   LITEVGYGIV PSF-- 348
GsBBX8c   LITEVGYGIV PSF-- 346
GmBBX19a  LFNEVGGSIF PTF-- 366
GsBBX19a  LFNEVGGSIF PTF-- 366
GmBBX13b  LFTEVGGSIF PTF-- 361
GsBBX13b  LFTEVGGSIF PTF-- 361
GmBBX5a   ----VGANAF PAYH- 365
GsBBX5a   ----VGANAF PAYH- 369
GmBBX8a   ----VGANAF PAYH- 371
GsBBX8a   ----VGANAF PAYH- 371
GmBBX7b   ----VGATAL PA--- 427
GsBBX7b   ----VGATAL PA--- 427
GmBBX10b  ----APPT-F PLLNK 419
GsBBX10b  ----APPT-F PLLNK 418
GmBBX20b  ----APPT-F PLLNK 418
GsBBX20b  ----APPT-F PLLNK 417
GmBBX2a   ---------- ---QK 340
GsBBX2a   ---------- ---QK 340
GmBBX10a  ---------- ---QK 348
GsBBX10a  ---------- ---QK 348
GmBBX3a   ---------- ---EH 349
GsBBX3a   ---------- ---EH 349
GmBBX19c  ---------- ---EH 351
GsBBX19c  ---------- ---EH 351
GmBBX19b  ---------- ---QA 385
GsBBX19b  ---------- ---QA 385
GmBBX16a  ---------- ---QA 381
GsBBX16a  ---------- ---QA 381
GmBBX2b   ---YDYDPLN QTRSY 405
GsBBX2b   ---YDYDPLN QTRSY 405
GmBBX14a  ---YDYDPLN QTRSC 411
GsBBX14a  ---YDYDPLN QTRSC 411
GmBBX13a  ---YDYDPLS TTRSF 409
GsBBX13a  ---YDYDPLS TTRSF 409
GmBBX20a  ---YDYDPLS TTRSC 409
GsBBX20a  ---YDYDPLS TTRSC 409
GmBBX13g  ---YDYDPLG -TRDI 423
GsBBX11h  ---YDYDPLG -TRDI 421
GmBBX12e  ---YDYDPLG -TRDI 423
GsBBX12e  ---YDYDPLG -TRDI 423
GmBBX1a   ---------- ----- 184
GsBBX1a   ---------- ----- 184
GmBBX11a  ---------- ----- 184
GsBBX11a  ---------- ----- 184
GmBBX11b  ---------- ----- 212
GsBBX11b  ---------- ----- 212
GsBBX12a  ---------- ----- 212
GmBBX12a  ---------- ----- 226
GmBBX4b   ---------- ----- 266
GsBBX4b   ---------- ----- 265
GmBBX6b   ---------- ----- 245
GsBBX6b   ---------- ----- 245
GmBBX17b  ---------- ----- 327
GsBBX17c  ---------- ----- 327
GmBBX14b  ---------- ----- 276
GsBBX14b  ---------- ----- 336
GmBBX4a   ---------- ----- 319
GsBBX4a   ---------- ----- 319
GmBBX6a   ---------- ----- 233
GsBBX6a   ---------- ----- 233
GmBBX11c  ---------- ----- 288
GsBBX11c  ---------- ----- 288
GmBBX12b  ---------- ----- 205
GmBBX12g  ---------- ----- 292
GsBBX12g  ---------- ----- 374
GmBBX13e  ---------- ----- 293
GsBBX11f  ---------- ----- 293
GmBBX9a   ---------- ----- 292
GsBBX9a   ---------- ----- 292
GmBBX11d  ---------- ----- 238
GsBBX11d  ---------- ----- 238
GmBBX12c  ---------- ----- 238
GsBBX12c  ---------- ----- 238
GmBBX13h  ---------- ----- 239
GsBBX11i  ---------- ----- 239
```

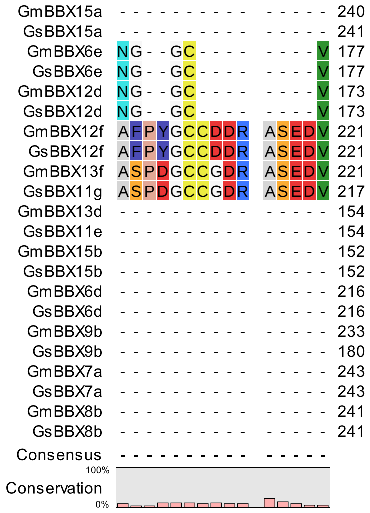
GmBBX15a - - - - - - - - - - - - - - - 240
GsBBX15a - - - - - - - - - - - - - - - 241
GmBBX6e NG - - GC - - - - - - - - V 177
GsBBX6e NG - - GC - - - - - - - - V 177
GmBBX12d NG - - GC - - - - - - - - V 173
GsBBX12d NG - - GC - - - - - - - - V 173
GmBBX12f AFPYGCCDDR ASEDV 221
GsBBX12f AFPYGCCDDR ASEDV 221
GmBBX13f ASPDGCCGDR ASEDV 221
GsBBX11g ASPDGCCGDR ASEDV 217
GmBBX13d - - - - - - - - - - - - - - - 154
GsBBX11e - - - - - - - - - - - - - - - 154
GmBBX15b - - - - - - - - - - - - - - - 152
GsBBX15b - - - - - - - - - - - - - - - 152
GmBBX6d - - - - - - - - - - - - - - - 216
GsBBX6d - - - - - - - - - - - - - - - 216
GmBBX9b - - - - - - - - - - - - - - - 233
GsBBX9b - - - - - - - - - - - - - - - 180
GmBBX7a - - - - - - - - - - - - - - - 243
GsBBX7a - - - - - - - - - - - - - - - 243
GmBBX8b - - - - - - - - - - - - - - - 241
GsBBX8b - - - - - - - - - - - - - - - 241
Consensus - - - - - - - - - - - - - - -
100%
Conservation
0%
