## Supplementary Fig. S for "Comparative analysis of *Glycine BBX* gene family reveals lineage-specific evolution and expansion": Supplementary fig 2-5.pdf

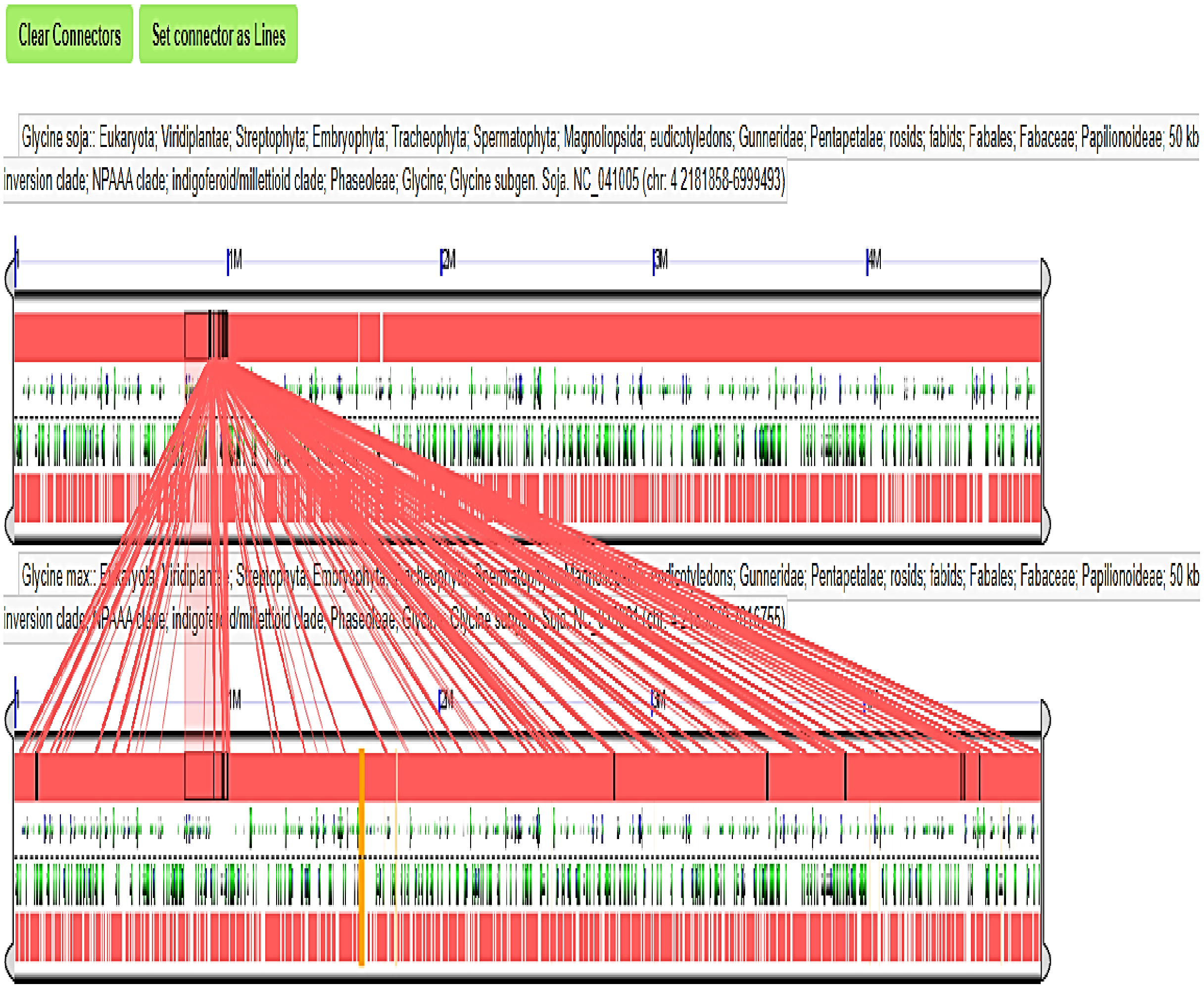


Supplementary Fig. S2. Microsynteny analysis between *G. max* and *G. soja* regions carrying *BBX* genes on *Gm04* and *Gs04*, respectively viz., *GmBBX4a*, *4b*, *4c* and *GsBBX4a*, *4b*, *4c*, using GEvo on a CoGe platform. (a) shows the region on Gs04. (b) shows the region on Gm04. The connector lines indicate regions of similarity. The black box represents the whole similarity box between the two genomic regions. The green and blue cylindrical structures represent coding regions within the genomic segment.

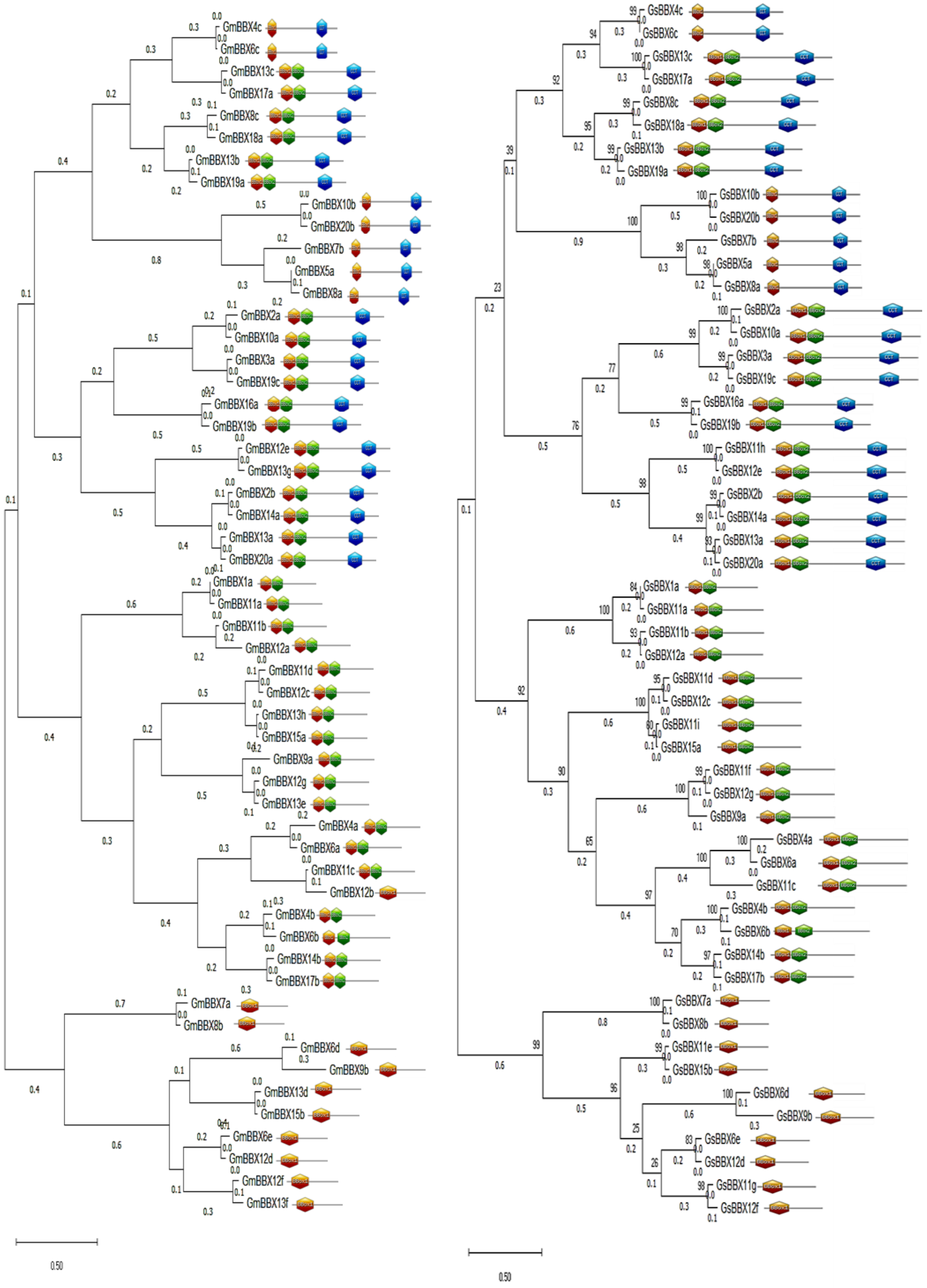


Supplementary Fig. S3. The phylogenetic classification of GmBBX and GsBBX proteins displaying member classification based on the BBX domain organization.

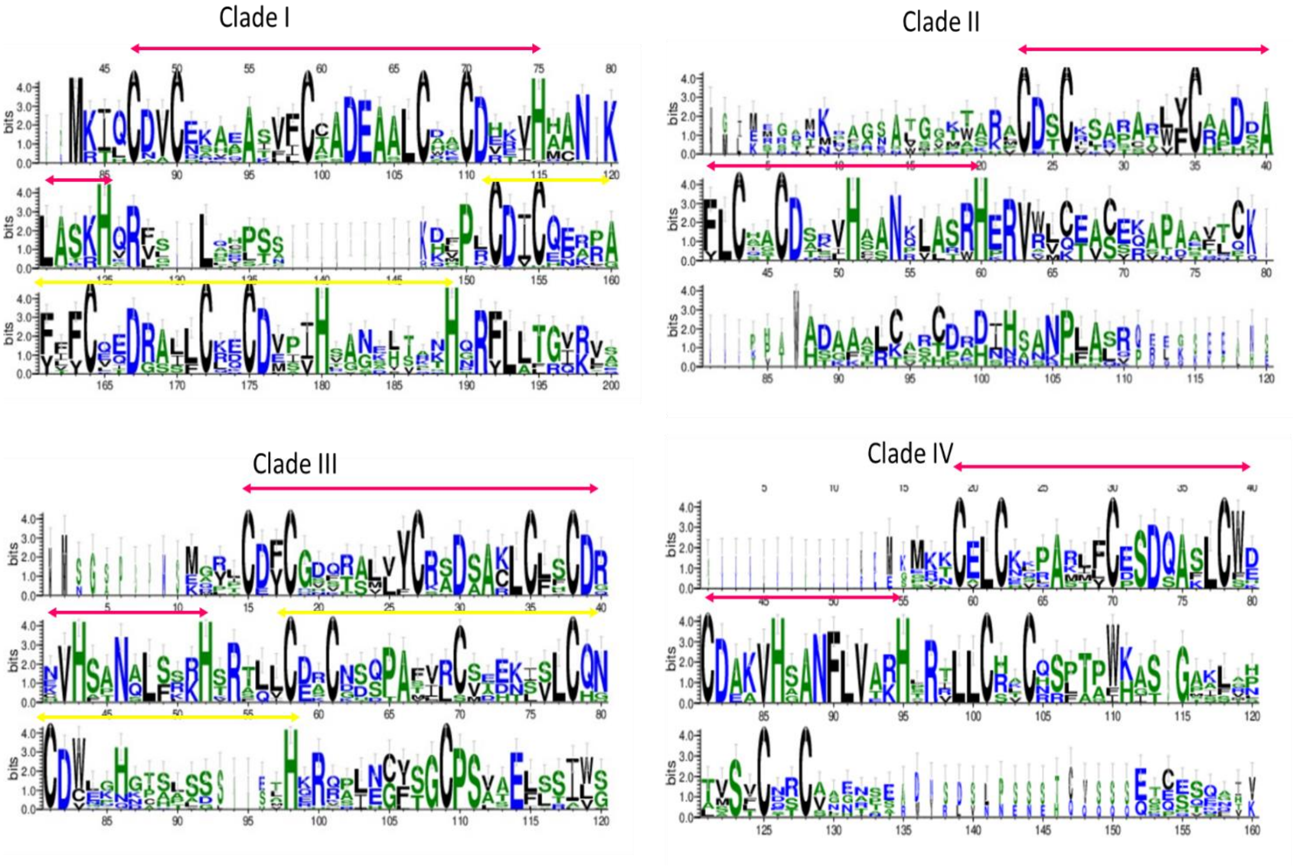


Supplementary Fig. S4. The clade classification of GmBBX and GsBBX proteins motifs using WebLogo.

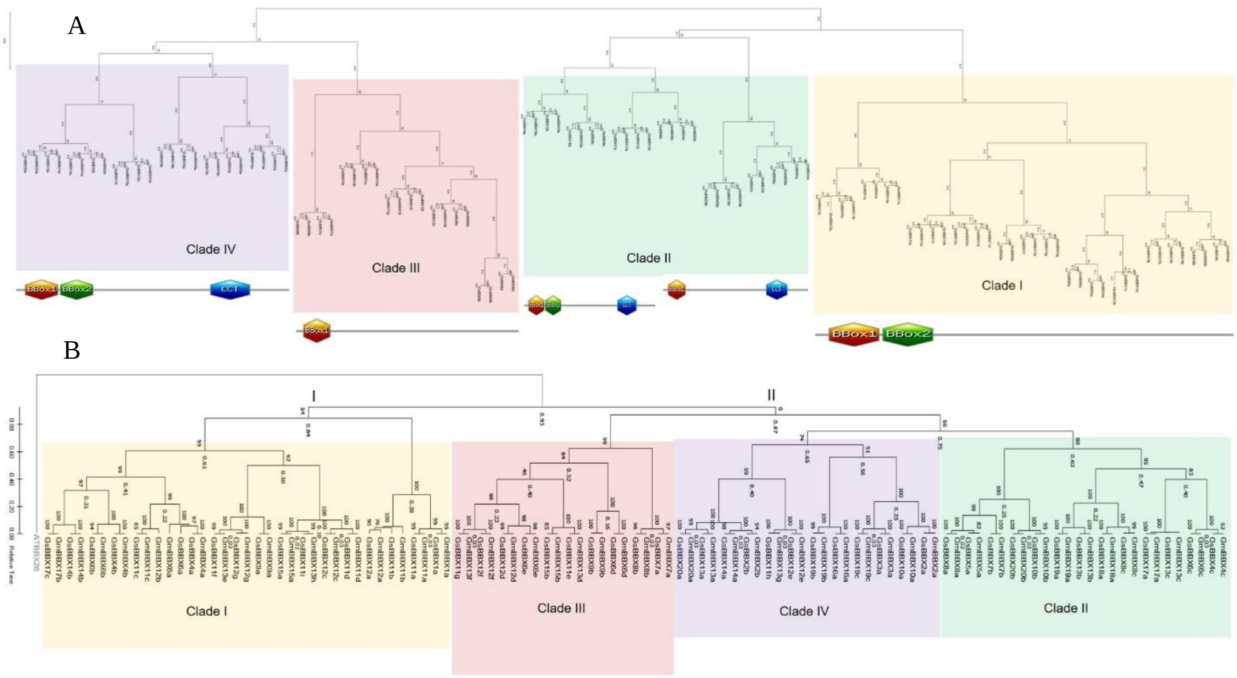


Supplementary Fig. S5. Phylogenetic classification of GmBBX and GsBBX proteins identifying BBX domains in different clades. (A) The phylogenetic clustering *Glycine* BBX proteins using ML. (B) The relative time tree of GmBBX and GsBBX proteins using AtBBX as an outgroup.
