## Supplementary Table for "Comparative analysis of *Glycine BBX* gene family reveals lineage-specific evolution and expansion": supplementary table 2.docx

| Group classification | GmBBX | GsBBX |
| --- | --- | --- |
| Group 1 (BBX1) | GmBBX7a, 8b, 6d, 9b, 13d, 15b, 6e, 12d, 12f, 13f | GsBBX7a, 8b, 6d, 9b, 11e, 15b, 6e, 12d, 12f, 11g |
| Group 2 (BBX1 and BBX2) | GmBBX1a, 11a, 11b, 12a, 11d, 12c, 13h, 15a, 9a, 12g, 13e, 4a, 6a, 11c, 12b, 4b, 6b, 14b, 17b | GsBBX1a, 11a, 11b, 12a, 11d, 12c, 11i, 15a, 9a, 12g, 11f, 4a, 6a, 11c, 4b, 6b, 14b, 17b |
| Group 3 (BBX1, BBX2 and CCT) | GmBBX13c, 17a, 8c, 18a, 13b, 19a, 2a, 10a, 3a, 19c, 16a, 19b, 2e, 13g, 2b, 14a, 13a, 20a, | GsBBX13c, 17a, 8c, 18a, 13b, 19a, 2a, 10a, 3a, 19c, 16a, 19b, 2e, 11h, 2b, 14a, 13a, 20a, |
| Group 4 (BBX1 and CCT) | GmBBX4c, 6c, 10b, 20b, 7b, 5a, 8a | GsBBX4c, 6c, 10b, 20b, 7b, 5a, 8a |

**Table 2** Classification of GmBBX and GsBBX proteins based upon the presence of one/two BBX domains
